# Genomic architecture and introgression shape a butterfly radiation

**DOI:** 10.1101/466292

**Authors:** Nathaniel B. Edelman, Paul B. Frandsen, Michael Miyagi, Bernardo Clavijo, John Davey, Rebecca Dikow, Gonzalo García-Accinelli, Steven van Belleghem, Nick Patterson, Daniel E. Neafsey, Richard Challis, Sujai Kumar, Gilson Moreira, Camilo Salazar, Mathieu Chouteau, Brian Counterman, Riccardo Papa, Mark Blaxter, Robert D. Reed, Kanchon Dasmahapatra, Marcus Kronforst, Mathieu Joron, Chris D. Jiggins, W. Owen McMillan, Federica Di Palma, Andrew J. Blumberg, John Wakeley, David Jaffe, James Mallet

## Abstract

We here pioneer a low-cost assembly strategy for 20 Heliconiini genomes to characterize the evolutionary history of the rapidly radiating genus *Heliconius*. A bifurcating tree provides a poor fit to the data, and we therefore explore a reticulate phylogeny for *Heliconius*. We probe the genomic architecture of gene flow, and develop a new method to distinguish incomplete lineage sorting from introgression. We find that most loci with non-canonical histories arose through introgression, and are strongly underrepresented in regions of low recombination and high gene density. This is expected if introgressed alleles are more likely to be purged in such regions due to tighter linkage with incompatibility loci. Finally, we identify a hitherto unrecognized inversion, and show it is a convergent structural rearrangement that captures a known color pattern switch locus within the genus. Our multi-genome assembly approach enables an improved understanding of adaptive radiation.

---

Adaptive radiations are responsible for most of today’s biodiversity. Initiated by key innovations and ecological opportunity, radiation is fueled by niche competition that promotes rapid diversification of species^1–3^. Reticulate evolution may aid radiation by introducing variation that enables rapidly emerging populations to take advantage of ecological opportunities^4,5^. Diverging from its sister genus ~12 million years ago, *Heliconius* butterflies radiated in a burst of speciation over the past five million years^6^. Their rapid diversification is likely due to their ability to feed on pollen, improved behavioral flexibility, tight coevolution with host plants, and Müllerian mimicry^7,8^, but we still know little about the genetic basis of these key innovations^8,9^. Introgression is well known in *Heliconius*, and particularly close attention has been paid to color pattern loci which have been passed between recently diverged species on several occasions^10–13^. Widespread reticulate evolution occurs across the genus^14^, but, aside from two pairs of closely related species^15,16^, we do not understand how introgression varies across the genome. Here, we use multiple *de novo* whole genome assemblies to study adaptive innovation, genome architecture, and introgression in *Heliconius*. We develop a new method to distinguish incomplete lineage sorting from introgression, and show that a majority of genomic loci that have evolutionary histories discordant with the canonical species tree arose due to introgression. We demonstrate that the distribution of introgressed loci is strongly influenced by local recombination rate, chromosome size, and coding sequence density. High-recombination and gene sparse regions of the genome are much more likely to harbor loci of hybrid origin, which implies an important role for linked selection in the fate of introgressing alleles. Genomic rearrangements are rare in *Heliconius*^17^, but we nonetheless found a convergent inversion around a known color pattern supergene.

We used an innovative short-read assembly method (*w2rap*^18^, an extension of DISCOVAR *de novo*^19,20^) to generate 20 new reference genome assemblies for species sampled from both major *Heliconius* sub-clades and three additional genera of Heliconiini (Supplementary Information Section 1). This strategy relies on high fidelity PCR-free Illumina sequencing, and is particularly powerful for low-complexity regions^21^. Our contigs were able to patch 30% of gaps between contigs of the more traditionally assembled *H. melpomene* Hmel2.5 genome^17,22^ (Supplementary Information Section 2). Our assemblies averaged an N50 of 47,976 bp, each composed of an average of 35,404 contigs (Supplementary Information Section 1). The assemblies had high ortholog scores, with an average of 98.9 percent of the BUSCO arthropod dataset recovered (93.3% complete and single copy, 3.5% complete and duplicated, 2.0% fragmented)^23^.

To investigate the *Heliconius* radiation, we aligned our highest quality *de novo* assemblies with representative Lepidoptera reference genomes^17,24–31^ (Supplementary Information Section 3). Previous work in *Heliconius* inferred a high level of phylogenetic discordance among genes, arguably a result of rampant introgression^6^, though this has been disputed^32^. We attempted to reconstruct a bifurcating species tree by estimating relationships using three different subsets of our alignment: protein-coding genes, conserved coding regions, and conserved non-coding regions. We generated species trees using coalescent-based and concatenation approaches, and using both the full Lepidoptera alignment and a restricted, Heliconiini-only sub-alignment. These results were largely congruent, but topologies differed in poorly supported parts of the tree (Figure 1A; all trees are given in Supplementary Information Section 4). For example, relative placements of the silvaniform species - *H. besckei, H. pardalinus*, and *H. numata* - were extremely unstable. In some datasets, *H. pardalinus* was recovered as sister to the *H. melpomene-cydno-timareta* subclade, while in other cases it formed a clade with other silvaniforms. Meanwhile, *H. besckei* was sometimes recovered branching from the root of the entire *melpomene-silvaniform* clade, and sometimes as sister to *H. numata*. Furthermore, *Heliconius telesiphe* showed higher affinity with the *H. sara-demeter* subclade in some datasets, while in others it was recovered as sister to *H. hecalesia* or branching from the root of a clade containing *H. hecalesia, H. erato*, and *H. himera*. Therefore, even with whole genome assemblies, we were unable to resolve the phylogeny of *Heliconius* as a simple bifurcating tree (Figure 1A, Supplementary Information Section 4).

**Figure 1:**
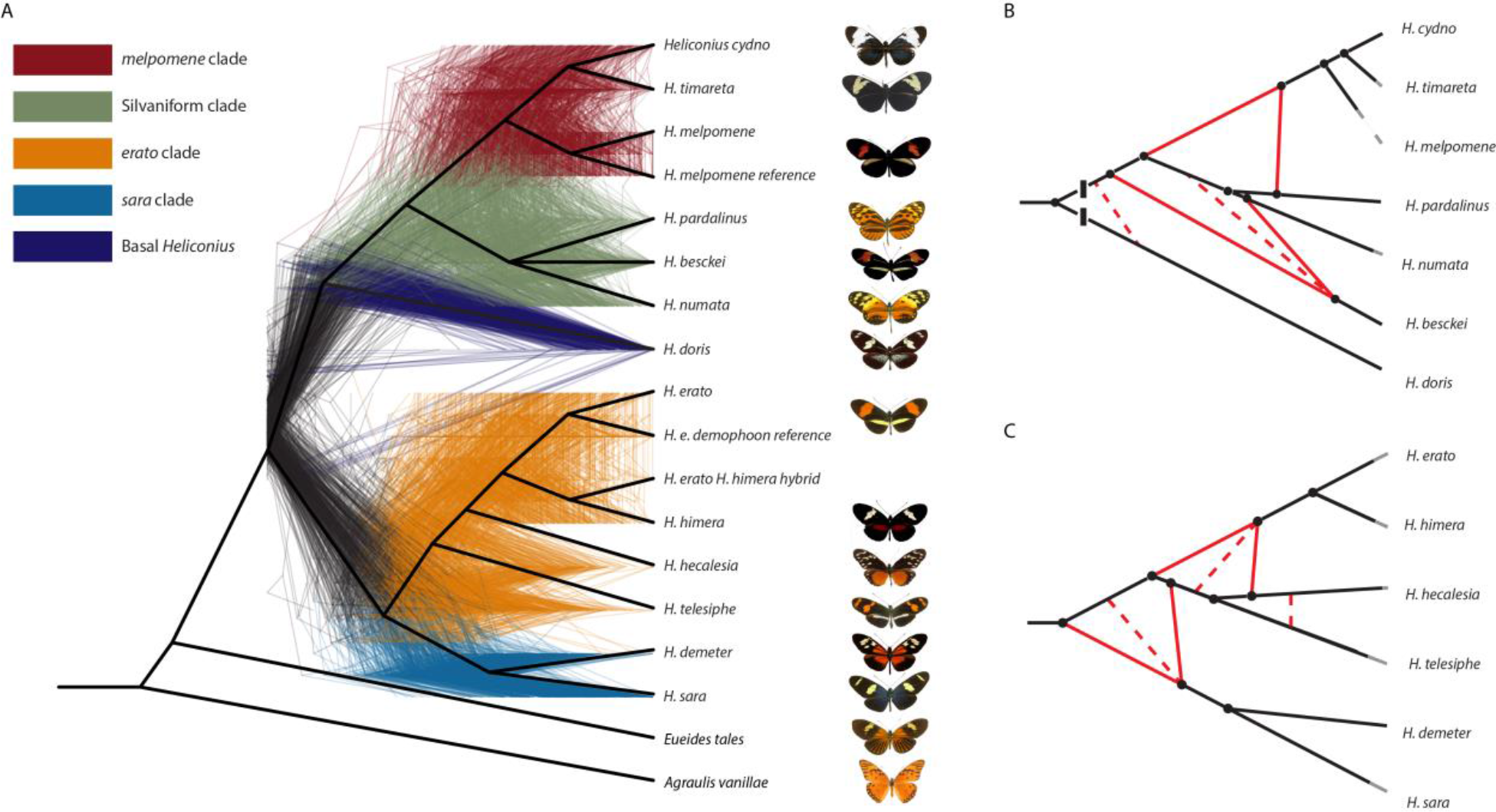
Phylogeny and phylogenetic networks of *Heliconius* show lack of support for bifurcating tree. **A. Consensus cladogram of *Heliconiini*.** Species trees were constructed with genes, fully-aligned coding blocks, and fully-aligned non-coding blocks. All nodes resolved in a majority of these species trees are shown in this cladogram (heavy black lines), while less well-resolved nodes are collapsed as a polytomy (see silvaniform species). The 500 colored trees were sampled from 32,000 10KB non-overlapping windows. Their discordance demonstrates the heterogeneity of evolutionary history across the genome. **B, C. Phylogenetic networks inferred with PhyloNet.** High-confidence tree structure (black) and introgression events (red) are shown as solid lines. Dashed red lines indicate weakly supported introgression events. The *erato-sara* clade is shown in **B**, the *melpomene-silvaniform* clade in **C.** Lengths of solid black lines are proportional to genetic distance along the branches. Breaks at the base in **B** indicate that the branch leading to *H. doris* has been shortened for display. See Supplementary Information Section 6 for full details.

To test whether this species tree uncertainty is due to incomplete lineage sorting (ILS) or introgression, we calculated Patterson’s *D*-statistics for every triplet of *Heliconius* species, holding *Eueides tales* as a constant outgroup^33,34^. In 201 of 364 triplets, we observed values significantly different from zero, implying significant evidence for introgression (Bonferroni-corrected *p*<.05, Supplementary Information Section 5). Although these significant *D* values suggest a history of admixture in the genus, this test alone provides little information on the number of admixture events or the fraction of genetic information transferred. We therefore used a different method, phyloNet^35,36^, to infer reticulate phylogenetic networks of the evolutionary history of these species (Figure 1B-C). Specifically, we randomly sampled sets of one hundred 10 KB windows across our alignment. For each such subset of the data, we co-estimated each regional gene tree and the overall species network in parallel^36^. We included chromosome-level reference genomes of *H. melpomene* and *H. erato* in our multi-species alignment; thus we analyzed the *melpomene-silvaniform* group with respect to the *H. melpomene* Hmel2.5 assembly^17^ and the *erato-sara* group with respect to the *H. erato demophoon* v1 assembly^24^. We repeated this analysis with 100 subsets of our data for the melpomene-silvaniform clade and 150 subsets for the *erato-sara* clade to evaluate support for each network (Supplementary Information Section 6). These networks show that introgression is pervasive across *Heliconius*. Nearly every species participated in an admixture event at some point in its history, and we were able to confirm extensive reticulation among silvaniform species in the *melpomene-silvaniform* clade, as well as discover major new gene flow events in the *erato-sara* clade. Based on these results, we propose the reticulate phylogenies in Figure 1B-C, in which most events occurred not among extant populations but among common ancestors of species we studied, and likely involved species not included here^14^.

To identify genetic changes that may have triggered the *Heliconius* radiation, we focused on genomic regions with significant shifts in evolutionary rate at the base of the genus. A total of 38,490 candidate loci, averaging 172 bp in length, were found specifically along the branch leading from the common ancestor of *Heliconius* + *Eueides* to *Heliconius* (Supplementary Information Section 7). Of these, 18,186 overlapped with 7,522 annotated genes (exonic or intronic). Gene Ontology (GO) term searches of these loci revealed an overrepresentation of accelerated genes involved in neuronal processes, specifically axon guidance and several terms related to ion channel structure and function (Table 1, Supplementary Information Section 7). *Heliconius* have pronounced behavioral complexity compared with other Lepidoptera, with learned home ranges and gregarious roosting behaviors as adults^37^. Concomitantly, mid-brain and mushroom bodies are unusually well developed in the genus, and the accelerated genes identified here provide candidates for these important phenotypes^38^.

**Table 1.**

| GO ID | GO Name | GO Category | Size | ES | NES | Nominal p-v | FDR q-val | FWER p-val | Rank at Max | Leading Edge |
| --- | --- | --- | --- | --- | --- | --- | --- | --- | --- | --- |
| GO:0007411 | axon guidance | BIOLOGICAL PROCESS | 22 | 0.71448416 | 2.2696512 | 0 | 0.00197327 | 0.007 | 1317 | tags=50%, list=15%, signal=59% |
| GO:0015299 | solute:proton antiporter activity | MOLECULAR FUNCTION | 10 | 0.81757474 | 2.076241 | 0 | 0.00824198 | 0.165 | 1311 | tags=40%, list=15%, signal=47% |
| GO:0005219 | ryanodine-sensitive calcium-release channel activity | MOLECULAR FUNCTION | 6 | 0.90566915 | 1.9730856 | 0 | 0.01828617 | 0.508 | 812 | tags=100%, list=9%, signal=110% |
| GO:0051209 | release of sequestered calcium ion into cytosol | BIOLOGICAL PROCESS | 6 | 0.90566915 | 1.9723397 | 0 | 0.01812592 | 0.514 | 812 | tags=100%, list=9%, signal=110% |
| GO:0009152 | purine ribonucleotide biosynthetic process | BIOLOGICAL PROCESS | 36 | 0.5674986 | 1.9656404 | 0 | 0.01924711 | 0.549 | 434 | tags=22%, list=5%, signal=23% |
| GO:0005891 | voltage-gated calcium channel complex | CELLULAR COMPONENT | 9 | 0.80593866 | 1.9653816 | 0.00132802 | 0.01888521 | 0.551 | 1285 | tags=89%, list=15%, signal=104% |
| GO:0009168 | purine ribonucleoside monophosphate biosynthetic process | BIOLOGICAL PROCESS | 25 | 0.6139621 | 1.959235 | 0 | 0.01879137 | 0.574 | 365 | tags=20%, list=4%, signal=21% |
| GO:0005245 | voltage-gated calcium channel activity | MOLECULAR FUNCTION | 10 | 0.76535124 | 1.9450445 | 0 | 0.01843062 | 0.646 | 1285 | tags=80%, list=15%, signal=94% |
| GO:0031984 | organelle subcompartment | CELLULAR COMPONENT | 53 | 0.510213 | 1.9105113 | 0 | 0.02531921 | 0.804 | 2296 | tags=55%, list=27%, signal=74% |
| GO:0044433 | cytoplasmic vesicle part | CELLULAR COMPONENT | 21 | 0.6063042 | 1.9100221 | 0.00119617 | 0.02513916 | 0.808 | 2569 | tags=71%, list=30%, signal=102% |
| GO:0035725 | sodium ion transmembrane transport | BIOLOGICAL PROCESS | 12 | 0.70204234 | 1.8933499 | 0.0025413 | 0.02913918 | 0.86 | 2553 | tags=92%, list=30%, signal=130% |
| GO:0042455 | ribonucleoside biosynthetic process | BIOLOGICAL PROCESS | 7 | 0.8435898 | 1.8900206 | 0.00135685 | 0.02996308 | 0.867 | 340 | tags=43%, list=4%, signal=45% |
| GO:0005887 | integral component of plasma membrane | CELLULAR COMPONENT | 54 | 0.49249074 | 1.8530432 | 0.00110254 | 0.04020768 | 0.952 | 2553 | tags=63%, list=30%, signal=89% |
| GO:0009206 | purine ribonucleoside triphosphate biosynthetic process | BIOLOGICAL PROCESS | 18 | 0.6313657 | 1.8367563 | 0.00370828 | 0.04228776 | 0.974 | 170 | tags=11%, list=2%, signal=11% |
| GO:0019318 | hexose metabolic process | BIOLOGICAL PROCESS | 15 | 0.63988847 | 1.832312 | 0.00251572 | 0.04348195 | 0.977 | 509 | tags=33%, list=6%, signal=35% |

Abbreviations:
ES: Enrichment Score, NES: Normalized Enrichment Score, FDR: False Discovery Rate, FWER: Family-Wise Error Rate

We next analyzed the distribution of ancestry across the genome by constructing trees in 50 KB sliding windows (25 KB overlap), again analyzing each major clade separately and using its respective reference genome. This larger window size allows us to generate a more contiguous map (i.e. each window is more likely to have sufficient data to generate a well-supported tree), and the relative abundance of each tree is qualitatively consistent regardless of block size (Supplementary Information Section 8). The *melpomene*-silvaniform group displayed a striking lack of consensus, unsurprisingly since introgression, especially of key mimicry loci, is well known from this clade^10,12–15^. The most common topology was found in only 4.3% of windows, with an additional 14 topologies appearing in 1-3.4% of windows (see Supplementary Data on Dryad). By contrast, in the *erato-sara* group, two topologies dominated the distribution, one of which (Tree 2, Figure 2B) matched our bifurcating consensus tree (Figure 1A) and the most recently published tree^6^. All eight of the most common trees in the *erato-sara* group, together comprising over 98% of all topologies, recovered monophyly of *H. demeter* + *H. sara* and *H. erato* + *H. himera*; only the placements of *H. telesiphe* and *H. hecalesia* were variable (Figure 2A-B, Supplementary Information Section 8). These results confirm the prevailing notion that introgression is most widespread in the melpomene-silvaniform clade, but also strengthens evidence of hybridization in the less studied *erato-sara* clade^14^.

**Figure 2:**
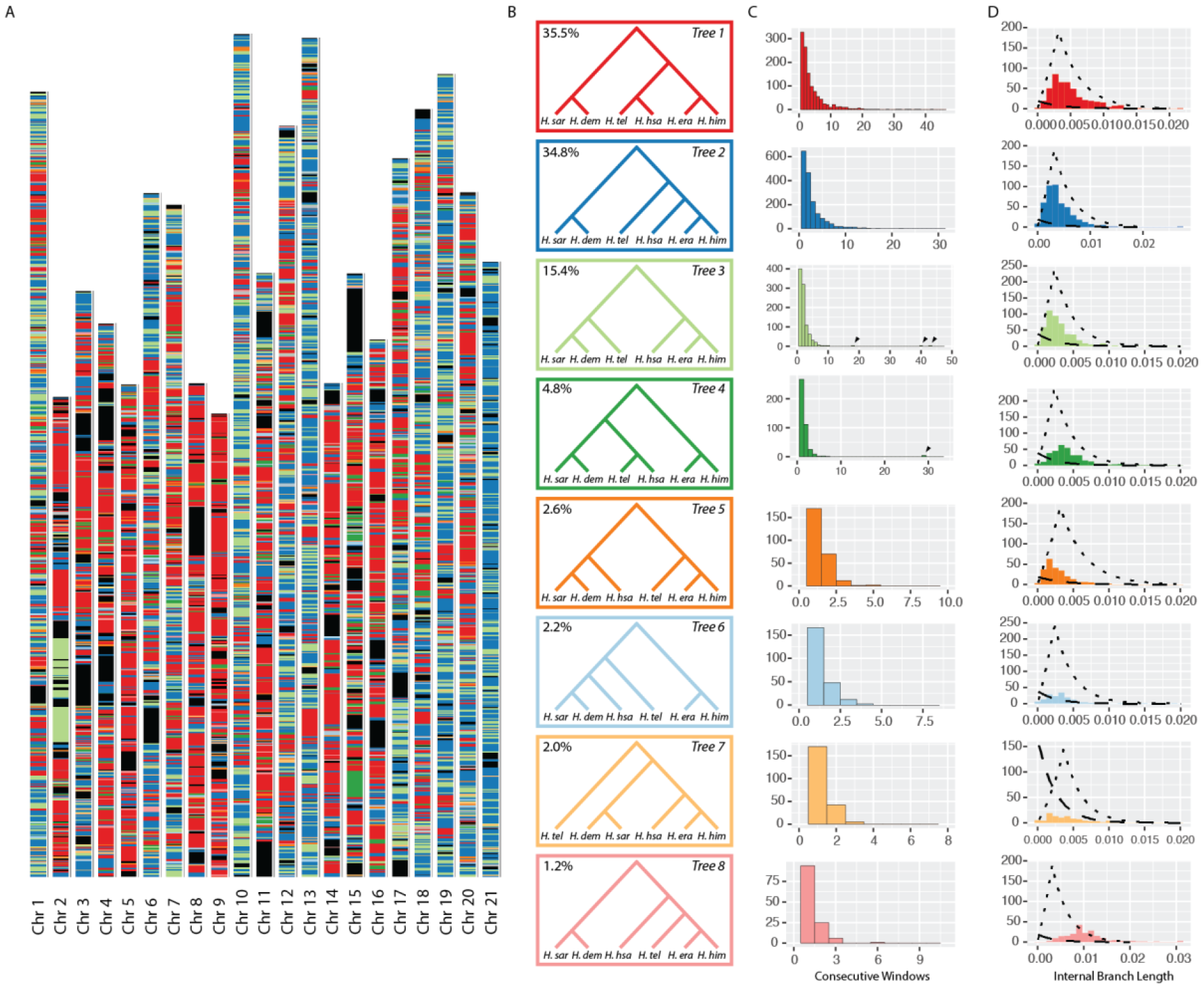
Local evolutionary history in the *erato-sara* clade is heterogeneous across the genome. **A. Tree topologies across the genome.** For each 50KB window, the colored bar represents the topology recovered from that region. Colors correspond to topologies in **B.** Note that while trees 1 and 2 (blue and red) dominate the genome, there is a large block of tree 3 on Chromosome 2, and a large block of tree 4 on Chromosome 15. Also note the relative homogeneity of Chromosome 21, the Z chromosome. Coordinates are in terms of *H. erato demophoon* v1^24^. Black regions show missing data. **B. Common topologies.** The eight most common trees are shown. The value in the top left corner is the percentage of all 50KB windows that recovers that topology. **C. Tree block length distribution.** Each histogram corresponds to the topology of the same color in B, and shows the distribution of the number of consecutive 50KB windows with that topology. Arrows indicate blocks in inversions. **D. Discriminating introgression from incomplete lineage sorting (ILS).** Plots show the distribution of internal branch lengths in the triplet *H. demeter, H. telesiphe, H. erato* for 10KB windows of each full gene tree type. Dashed lines show the exponential distribution of branch lengths generated by ILS, while dotted lines show the peaked distribution of branch lengths generated by speciation or introgression. Both distributions are conditioned on the triplet topology of the corresponding full gene tree, and therefore vary depending on whether *H. demeter, H. telesiphe*, or *H. erato* is recovered as the outgroup.

Regions that show a local topology discordant with the species tree could have arisen through introgression or ILS, and we were interested in distinguishing between these processes. We here develop a triplet gene tree test that compares the likelihoods that a given gene tree results from ILS or introgression. Our test is based on the distribution of internal branch lengths among windows for a given three-taxon subtree, conditional on its topology. For example, in the absence of introgression, the internal branches of triplet topologies that are discordant with the species tree (due to ILS) should be exponentially distributed. However, if introgression has occurred, their distribution should have a non-zero mode which corresponds to the time from the introgression event back to the most recent common ancestor of all three species (Supplementary Information Section 9). Using this model, we can discriminate between discordant gene tree topologies that are consistent with ILS and its expected exponential distribution (e.g. Tree 7, Figure 2D), and those that are not (e.g. Tree 4, Figure 2D). Once we have identified nodes that have likely experienced introgression, we can then examine each window and calculate the probability that it was generated through ILS (See Supplementary Information Section 9). With this method, we found that 87 percent of windows that recovered full trees discordant with our inferred species tree (Tree 2, Figure 2B) contain at least one triplet relationship that most likely (probability >= 0.9) arose due to non-ILS processes such as introgression.

Given that introgression appears variable across the genome, we next investigated correlates of this variation. In hybrid populations, individuals may have genomic regions that originate from other species and are incompatible with the recipient genome or with their environment^39,40^. As hybrid individuals backcross into a parental species, incompatible loci experience negative selection in accordance with their fitness costs, and are expected to be purged. Linked selection causes harmless or even beneficial introgressed loci to be removed along with these deleterious loci if they are tightly linked; this effect will depend on the strength of selection against deleterious loci and the local recombination rate^41,42^. We expect introgressed loci to be enriched in regions where selection is likely to be weak, such as gene deserts, or in regions of high recombination, where harmless introgressed loci more readily recombine away from linked incompatibility loci.

In *Heliconius* even distant species like *H. erato* and *H. melpomene* have the same number of chromosomes (21), and they are almost entirely collinear^17^, facilitating comparisons among species. Furthermore, the map lengths of each of the chromosomes is close to 50 centiMorgans (cM), corresponding to one crossover per chromosome per meiosis in males (there is no crossing over in female *Heliconius*)^22,24^. Chromosomes vary in length, and larger chromosomes therefore experience a lower per-base recombination rate than shorter chromosomes^16,17^ (Figure 2). For this analysis, we generated trees in 10 KB windows, and found a striking correlation between the fraction of windows in each chromosome that show a given topology and physical chromosome length (Figure 3A). Such relationships exist for all 8 trees in Figure 2B (Supplementary Information Section 8), but here we focus on the two most common trees: while trees 1 and 2 are found in an almost identical fraction of the genome, Tree 1 has a strongly negative correlation with chromosome size while Tree 2 (concordant with our inferred species tree) has a positive correlation.

**Figure 3:**
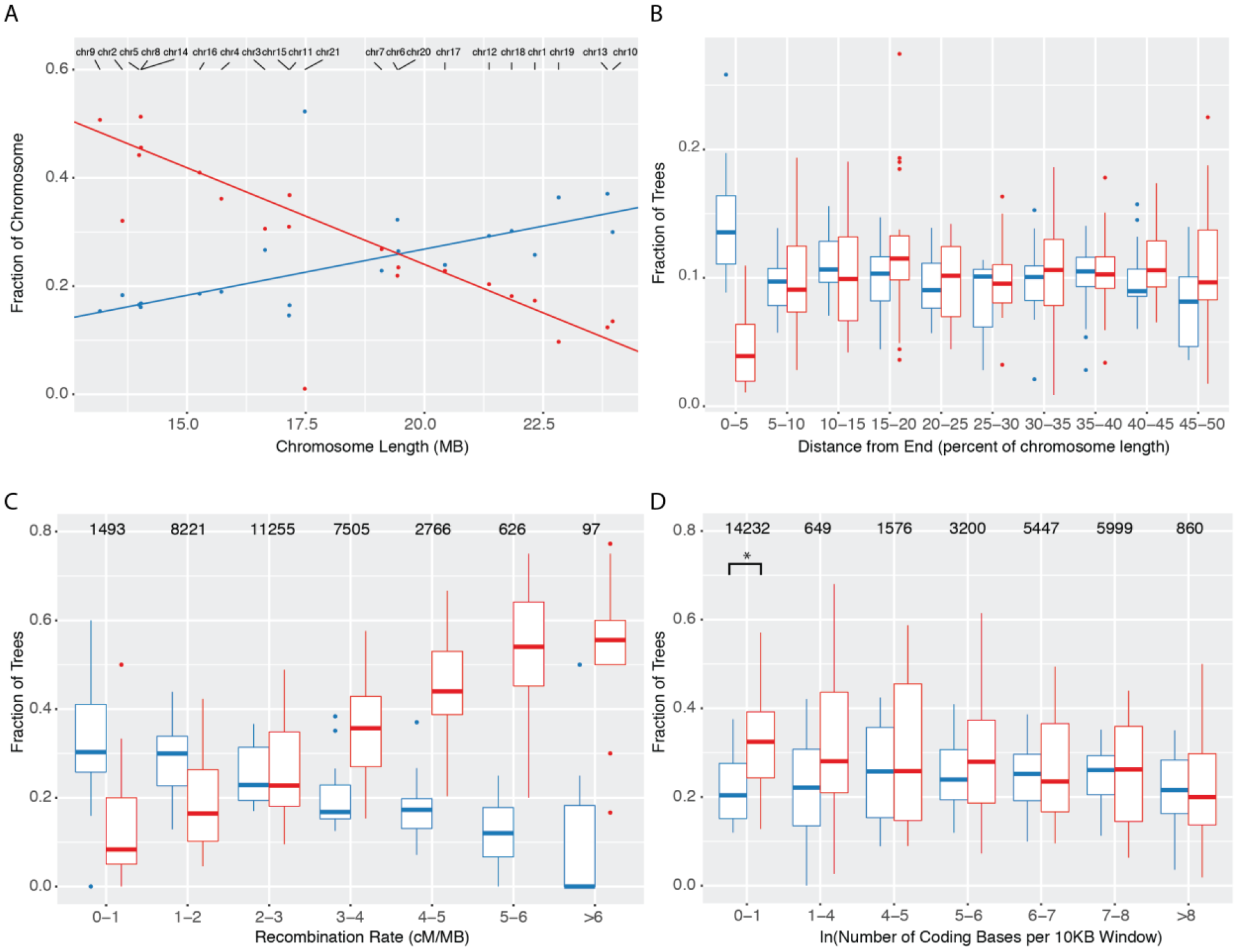
Chromosomal architecture is strongly correlated with local topologies. In all panels, Tree 1 is shown in red, and Tree 2 is shown in blue, as in Figure 2. **A. Topology correlates with chromosome size**. Tree 1 shows a negative relationship with chromosome size, while Tree2 shows a positive relationship. Lines are linear regressions with chromosome 21 excluded (Tree 1 p<.0001, r^2^=.883, Tree 2 p<.0001, r^2^=.726). **B. Topology is altered at the ends of chromosomes.** Each chromosome was divided into 10 equally sized bins, and the occupancy of each topology in each bin was calculated as the number of windows that recovered the topology in the bin divided by the number of windows that recovered the topology in the chromosome. **C. Topology correlates with local recombination rate.** We used the LD-based map of *H. erato demophoon*^24^ to assign a local recombination rate to each window. We then binned windows by their recombination rate, and calculated the fraction of each tree in each bin for each chromosome separately. Numbers above bars are the number of windows in each bin. **D. Density of coding sequence correlates slightly with topology.** Numbers of coding bases per window are distributed exponentially, so the data for this analysis are log-transformed (Supplementary Information Section 8). Asterisk denotes significance at 5% level (paired t-test, p=.025). In all boxplots, central line is median, box edges are first and third quartile, and whiskers extend to the largest value no further than 1.5*(inter-quartile range) ^47^

The most drastic exception is the Z, chromosome 21, which is the most strongly enriched for Tree 2. Because Tree 2 is the inferred species tree in the *erato-sara* clade, this result is consistent with the Z chromosome harboring or expressing a higher proportion of incompatibility loci than autosomes. Interspecific hybrid females in *Heliconius* are often sterile, thereby conforming to Haldane’s Rule, and sex chromosomes have been implicated as particularly important in generating this incompatibility^40,43–46^.

To confirm that the pattern we see among chromosomes is truly due to differences in recombination, we next investigated the relationship between recombination rate and tree topology within chromosomes. Recombination rate declines at the ends of chromosomes (Supplementary Information Section 8), and we find enrichment for the proposed species tree (Tree 2) in those regions (Figure 3B). In addition, even with the low-density recombination map of *H. erato*, when windows are grouped by local recombination rate calculated from population genetic data, we see a strong relationship with the recovered topology (Figure 3C). Finally, we observe a minor enrichment of Tree 1 in regions of very low gene density, but this effect is weak compared to that of recombination rate (Figure 3D, Supplementary Information Section 8). Taken together, these results imply an important role for linked selection in determining the hybrid makeup of *erato-sara* clade genomes.

While local recombination has a strong effect on the location of different topologies, the topology block size distribution in the *erato* clade (the number of consecutive sliding windows that correspond to a single topology) generally decayed exponentially (figure 2C). However, two unusually long blocks contained minor topologies, one on chromosome 2 (Tree 3, composed of three sub-blocks) and the other on chromosome 15 (Tree 4). Our study of the ~3 Mb topology block on chromosome 2 confirms the earlier suggestion of an inversion in *H. erato*^17^, and we show here that its rare topology can be explained by ILS including a long period of ancestral polymorphism (Supplementary Information Section 10).

The topology block on chromosome 15 is of particular interest, as it spans a mimicry switch locus containing the gene *cortex* (known as *Cr* in *H. erato, Yb* in *H. melpomene*, and *P*_1_ in *H. numata*) ^48,49^. The *cortex* topology block in the *erato* group is ~500 KB in length and recovers *H. telesiphe* and *H. hecalesia* as a monophyletic subclade, which together are sister to the *sara* clade (Figure 2B, Tree 4). We hypothesized that this block could be an inversion, as in *H. numata*, where introgression of the *P*_1_ inversion triggered the evolution of a ‘supergene’ polymorphism that enabled switching among mimicry morphs^12^. Our *de novo* assemblies enabled an effective search for contigs that mapped across topology transitions. Taking *H. melpomene* as the standard arrangement, we find clear inversion breakpoints in *H. telesiphe, H. hecalesia, H. sara*, and *H. demeter*. Conversely, *H. erato, H. himera*, and *E. tales* all contain contigs that map in their entirety across the breakpoints (Figure 4A), implying the standard *H. melpomene* arrangement is ancestral.

**Figure 4:**
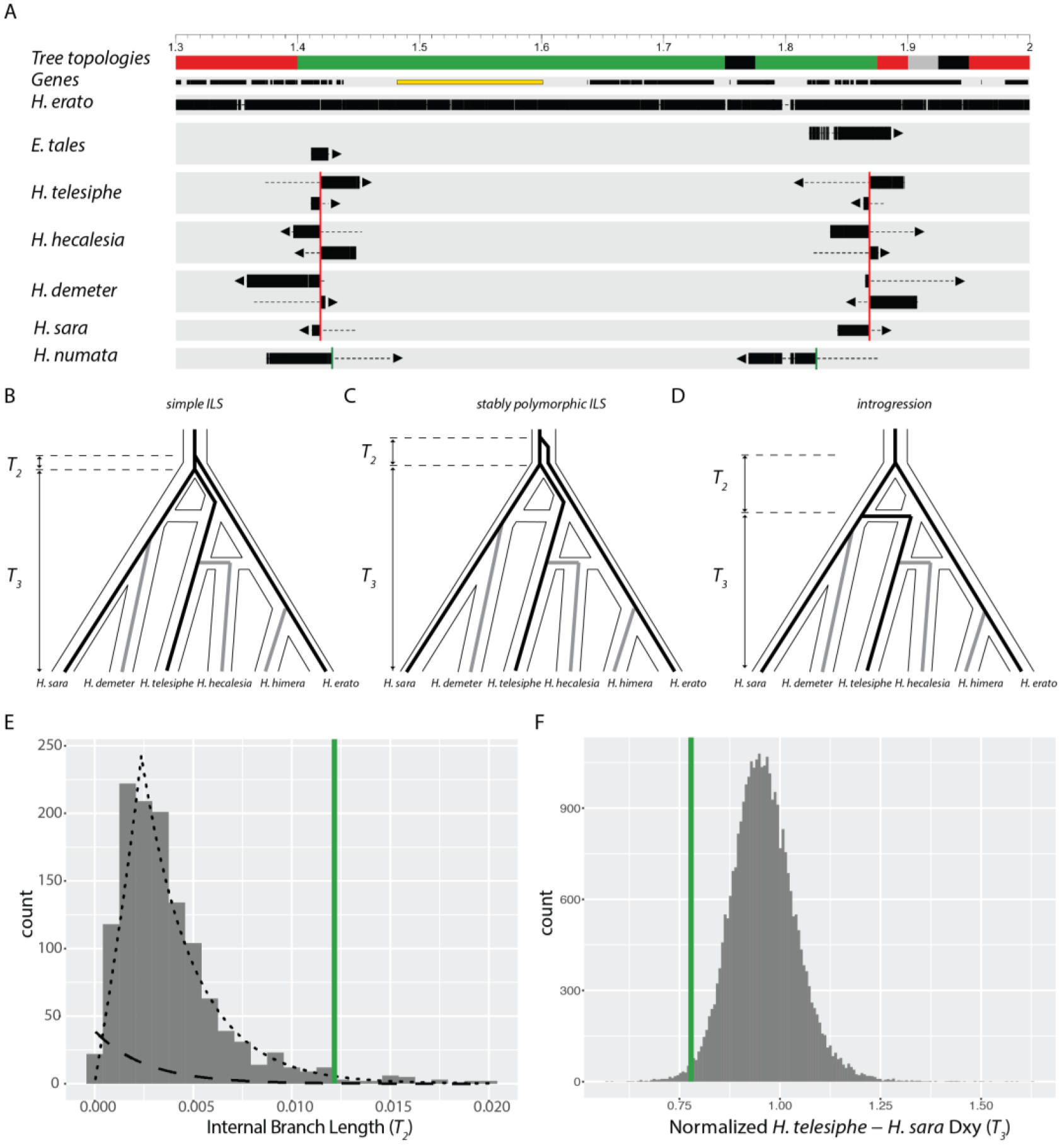
Parallel evolution of a major inversion at the *cortex* supergene locus. **A. Mapping the inversion breakpoints.** Map of 1.7Mb region on Chromosome 15. Coordinates are in terms of the Hmel 2.5 reference order, and ticks are in MB. Tree topology colors correspond to those in figure 2. Genes are shown as black rectangles; cortex is highlighted in yellow. Each line shows the mapping of a single contig. Aligned sections of each contig are shown as thick bars, while unaligned sections are shown as dotted lines. Arrows indicate the strand of the alignment. The *H. erato* group breakpoints are shown with red vertical lines, while the independent *H. numata* breakpoints are shown with green vertical lines. **B-D. Hypothetical evolutionary histories.** The histories of the three species used in the triplet gene tree method - *H. erato, H. telesiphe*, and *H. sara* - are shown as black lines, while lineages not included are shown as grey lines. **B** shows the scenario expected in the case of simple ILS. **C** shows another scenario of ILS, but in this case the inversion is polymorphic for some time in the common ancestor. **D** shows the case of introgression from the *H. sara+H. demeter* ancestor into the *H. telesiphe+H. hecalesia* ancestor. **E. Distribution of internal branch lengths.** Histogram is of internal branch lengths (*T*_*2*_) in the *H. erato*,(*H. telesiphe, H. sara*) topology. The inferred ILS distribution is shown as a dashed line, and the inferred introgression distribution is shown as a dotted line. The average internal branch length in the inversion is shown as a green vertical line. **F. Normalized *H. telesiphe-H. sara* D_XY_.** Normalized D_XY_ (*T*_*3*_) is calculated as *H. telesiphe-H. sara* D_XY_ divided by the mean pairwise D_XY_ among all species in each region. Mean normalized D_XY_ in the inversion is shown as a green vertical line.

Thus, the chromosome 15 topology block appears to have been generated by an inversion at almost exactly the same position as the 400 KB *P*_1_ inversion in *H. numata*^12,49,50^, but it must have evolved in parallel: our *de novo* contigs from the *P*_1_ inversion in *H. numata* show that the breakpoints of *P*_1_ are close but not identical to those of the inversion in the *erato* clade (Figure 4A). Furthermore, in topologies of *H. numata*, *H. telesiphe*, *H. erato*, and *E. tales* in 50 KB sliding windows across chromosome 15, not a single window recovered *H. numata* and *H. telesiphe* as a monophyletic subclade, as would be expected if the *erato* group inversion was homologous to *P*_1_ in *H. numata*.

Using our triplet gene tree test, we examined the triplet (*H. erato* +*H. telesiphe* + *H. sara*) to elucidate the evolutionary history of the inversion. With these species, the topology of interest is (*H. erato*,(*H. telesiphe, H. sara*)), where a small internal branch would suggest ILS while a large internal branch would be more consistent with introgression (Figure 4B,D). We find that the average internal branch length in the inversion has only a 6.1% percent chance of being generated due to simple ILS (Figure 4E). However, if the inversion was polymorphic in the ancestral population for some time, we should also recover a long internal branch (Figure 4C). We distinguish between this longer-term polymorphic scenario and introgression by comparing the genetic distance (*D*_*XY*_) between *H. telesiphe* and *H. sara*. Normalized *D*_*XY*_ within the inversion is considerably lower than in the rest of the genome, a result that is only predicted in the case of introgression (Figure 4F).

## Discussion

In the *Heliconius* radiation, the long-term fate of introgressed loci is tightly dependent on local genomic context. Whether measured at the local or chromosomal scale, genomic regions with high recombination are much more likely to have evolutionary histories consistent with introgression. This pattern is expected if incompatibility loci are common and distributed across the genome, as found in recent studies^51–53^, and if they generate linked selection against such loci over a longer range in low recombination regions. In extant hybrid populations, linked selection has been proposed as the mechanism by which hybrid ancestry is positively associated with local recombination rate^37,45^. If robust, this relationship may be useful in resolving complex phylogenetic relationships in groups that have undergone introgression. In our case, two tree topologies were recovered in approximately equal proportions across the genome, but only the tree consistent with the published species tree shows a negative relationship with recombination rate (Fig. 3). Without prior knowledge, this relationship could arguably be used as evidence for the true species tree.

*Heliconius* experienced ten chromosomal fusions along its basal branch^10,22,25^. Therefore, *Heliconius* has fewer, and on average longer, chromosomes compared most other Heliconiini or Nymphalidae. Genome architecture plays a critical role in adaptation and divergence. Between species, chromosomal rearrangements have been argued to prevent recombination and to allow genomic regions to diverge in the face of gene flow^54^, though it is unclear if they are causal in speciation^55^. *Heliconius* chromosomal fusions may have aided speciation for similar reasons. Compared with *Melitaea* (with the ancestral Lepidopteran count of n=31 chromosomes), *H. melpomene* and *H. erato* have 34% and 42% less recombination in the genome, respectively, based on total cM of each map^10,15,16^. The overall reduction in recombination caused by these fusions likely led to conditions more favorable to speciation with gene flow, whereby introgressed alleles are more easily purged from longer chromosomes due to their reduced recombination rate, and co-evolved sets of genes are more likely to remain linked^22^.

Inversion polymorphisms can reduce local recombination and preserve linkage of co-evolved genes, allowing populations to occupy multiple peaks on adaptive landscapes^49,56^. *Heliconius* boasts one of the clearest examples of a “supergene” locus: a system of tandem inversions in *H. numata* that completely determines an individual’s color pattern in that polymorphic species^49,50^. This locus is all the more striking because *Heliconius* generally lack major structural rearrangements, either between sister species or between species that diverged at the earliest branch point in the genus^17^. A key inversion, the P_1_ inversion, was inherited *via* introgression from a related species, *H. pardalinus*^12^. We were therefore surprised to find in the current study that several *erato-sara* clade species have an inversion that evolved in parallel in almost precisely the same location, and which likewise shows a strong signal of introgression. Our assemblies are based on only a single individual per species, so we cannot determine whether this inversion is polymorphic in *erato-sara* clade species. The *cortex* gene is trapped within the inversion and has been implicated in color patterning across *Heliconius* and other Lepidoptera, but there is no clear phenotype that unites *erato-sara* clade species having this inversion. Nonetheless, due to the known effects of *cortex* and the fact that convergent structural rearrangements have been found rarely and in important contexts such as mating loci and meiotic drive loci in fungi^57,58^, and as a basis for social organization in ants^59^, this locus should be the object of further study.

Technical advances in long-read, single-molecule sequencing strategies are revolutionizing *de novo* genome assembly of non-model organisms^60–62^. However long-read technologies remain error-prone, expensive, and require high coverage for successful assembly. In contrast, here we pioneer a low-cost approach that employs the extremely low error rate inherent in PCR-free Illumina short-read technology to create high fidelity genome assemblies. We predict that ours will be the first of many studies to employ fully-aligned *de novo* genome assemblies for comparative analysis of adaptive radiation in non-model species. These new genomes with the help of two previous chromosome-level reference assemblies allow us to probe the history of the *Heliconius* radiation in unprecedented detail. We have discovered parallel evolution of an inversion around a supergene switch locus, a pattern of sustained hybridization and introgression throughout the genome, and candidate genes and genome architectures that help to understand the causes for the greater diversification rate in *Heliconius* compared with the rest of the Heliconiini.

## Methods

### Samples

Individuals for sequencing were collected mainly in the wild, while others were from partially inbred stocks (Table S1.1). For *Heliconius melpomene*, we used a sibling of the strain of the original reference individual for the Hmel1 and Hmel2.5 assemblies^10^. We also sequenced parents of a cross between *H. erato* and *H. himera*, and one of their F1 hybrid offspring - the *H. erato* mother was a sibling from the inbred strain used in the *H. erato demophoon* v1 genome^24^.

### Genome assembly with DISCOVAR de novo and w2rap

DNA was extracted from each specimen (Table S1.1). DNA samples were fragmented to ~450 bp and sequenced to at least 60X coverage using paired-end, 250 bp reads on the Illumina Hi-Seq 2500 according to DISCOVAR *de novo* protocol^20^. The w2rap-contigger^18^ was then used to assemble contigs (Table S1.2). Heterozygous genomes such as these yield complex assembly graphs where loci fail to collapse into a single representation but are expanded into two alternative alleles (Figure S1.1, left side). To overcome this problem, we filtered initial contigs to create a collapsed mosaic of haplotypes where each locus is represented by only one of the alternative alleles, ignoring phase, using the following procedure:

1. Homozygous content was selected using frequency of kmer spectra^63^; this is content shared by both haplotypes and in a mosaic collapsed scenario should be included only once in the solution.
2. Initial contigs were sorted by size and filtered one at a time starting from the longest.
3. A tally was kept for all included content and resulting contigs were filtered by comparing the content that they had from the selected unique set and not already included in the final set. If a contig contained mostly kmers already included in the final set this contig was marked as complementary and saved in a separate file.

This method will for the most part result in inclusion of only one alternative allele at each locus in the final set of contigs. This filtered set of contigs with a putative single representation for each locus was then further scaffolded, using the same paired-end sequence data, with SOAPdenovo2 scaffolder^64^ (Figure S1.1, right side)

### Assembly quality assessment

We used a custom python script employing standard formulae to extract the number of contigs, their lengths, and N50 scores. In order to estimate heterozygosity, we used the k-mer based method GenomeScope^65^. This online tool uses the output of jellyfish^66^, which we ran with the commands

jellyfish count -m 15 -o <count output> -s 2000M -t 16 -F 2 -C <(zcat <reads1.fastq>) <(zcat <reads2.fastq>)

jellyfish histo -t 16 <count output> > <hist output>

We characterized the repeat content with RepeatMasker v4.05^67^. For each genome, we used the command

RepeatMasker -pa 4 -species hexapoda -xsmall -nocut <genome.fa>.

We include full results on Dryad.

We used BUSCO_v2^23^ with the arthropoda_odb9 database to compute the percentage of complete, partial, duplicated, and missing genes in our genome assemblies. Specifically, we used the command:

BUSCO.py -i <genome fasta> -c 32 -m geno -l arthropoda_odb9-sp heliconius_melpomene 1 -o <output file>-r

We summarized the data with BUSCO_plot.py, using R to display the results.

### Alignment

w2rap Genomes with fewer than 50,000 scaffolds were aligned with previously published, representative genomes from across the lepidopteran phylogeny using progressiveCactus^68^. This tool generates pairwise alignments in cactus graph format between each pair of sister species, generating inferred ancestral sequences. It iteratively continues this process for each node of an input phylogeny, generating blocks of aligned homologous sequence among all input genomes. This process allows alignment blocks to be projected onto the coordinates of any genome assembly and produces little bias towards segments highly similar to any one reference genome. All configuration files are available on Dryad, and additional information can be found in Supplementary Information Section 3.

### Phylogeny

We extracted nine different datasets to compute phylogenies. These can be categorized into three main groups: conserved coding regions, conserved non-coding regions, and genes. All three types were generated for the full lepidoptera dataset and for the subset of species in *Heliconiini*. In addition, minimum alignment length filters of both 100 and 150 were used when extracting conserved coding and non-coding regions. Datasets consisting of conserved coding or noncoding regions were extracted by first converting the progressiveCactus hal-format multigenome alignment into Multiple Alignment Format (MAF), with *H. melpomene* Hmel2.5 as the reference. Each block in the MAF alignment was processed and retained if it contained exactly one sequence from each species. Genomic coordinates in BED format were extracted from the single-copy, fully-aligned MAF blocks and intersected with CDS entries in the Hmel2 annotation file to separate coding from non-coding blocks. Finally, each MAF block was converted to fasta using msa_view.

The genes were extracted by first extracting gene models from the Hmel2.5 annotation file with UCSC Kent binaries^69^, then using the resulting gene model bed file to extract hal alignments to MAF, and finally converting MAF to fasta with a custom python script.

For each of the nine datasets, we first estimated individual locus maximum likelihood trees (for gene-based trees, each locus consisted of all coding sequence within each gene, while the block trees were single contiguous stretches of fully-aligned sequence). We selected the best model for each locus using ModelFinder as implemented into IQtree^70,71^. Following model selection, we estimated the best ML tree and 1000 ultra-fast bootstraps using IQtree^72^. Using the resulting bestML trees from each locus as input, we generated species trees in ASTRAL^73^. For each dataset, we also generated concatenated supermatrices using FASconCAT v.1.1^74^. We selected models for each supermatrix, first by searching for the best partitioning scheme using the relaxed clustering algorithm implemented in IQtree, using the individual loci as the starting data blocks, and second, by estimating substitution models for each partitioning scheme subset using ModelFinder. We then used the best fit model partitioning scheme to estimate maximum likelihood trees on the supermatrix. For each data set we performed 10 ML searches, five with parsimony starting trees and five with random starting trees in IQtree and chose the topology with the best maximum likelihood score.

### Evolutionary rate

We extracted MAF alignments of each scaffold from the HAL-formatted whole genome alignment using the hal2maf command from the haltools suite^75^, with the *H. melpomene* genome as reference. We then converted MAF to SS with msa_view from the phast suite^76^. We created a null model of evolutionary rate with 4-fold degenerate sites for each scaffold separately with PhyloFit, and then averaged them together according to the PhastCons best practices^76^. Finally, we ran dless on each scaffold separately using the averages neutral model file and default parameters. We defined genic regions using bedtools based on ‘gene’ designations in the *Heliconius melpomene* Hmel2.5 gff3 file^17^.

GO-term enrichment analysis was carried out using Blast2GO^77^. The annotation file was generated by running blastp with all Hmel2.5 proteins against the NCBI refseq_protein database. GO mapping and annotation of the resulting output files was performed with the Blast2GO GUI using default parameters. A ranked list of dless hits was generated for coding and non-coding regions separately. LOD scores of accelerated regions remained positive values, while LOD scores of conserved regions were assigned negative values. We then performed a gene set enrichment analysis in the Blast2GO GUI, using the weighted method which searches for over-representation at both the top and bottom of the ranked list.

### Introgression analysis

#### ADMIXTOOLS

We first extracted MAF alignments of each scaffold from the HAL-formatted whole genome alignment using the hal2maf command from the haltools suite^75^, with the *H. melpomene* genome as reference. We then converted from MAF to FASTA using the msa_view tool from the PHAST toolkit^76^ FASTA to VCF using snp-sites^78^, and finally VCF to EIGENSTRAT using the vcf2eigenstrat tool from the gdc suite. Scaffold positions were converted to chromosomal positions using a custom python script. Sites were included if they were present in all species (i.e. no missing data or gaps) as a single copy, bi-allelic SNPs. This resulted in a set of 6,671,421 SNPs.

We computed all possible D statistics using the qpDstat command with default parameters, and considered all triplets of *Heliconius* species, holding *E. tales* constant as the outgroup. We assessed significance through a block-jackknifing approach as implemented in Admixtools^34^, and applied a Bonferroni correction to assign significance at the 95% confidence level to a p-value of 1.37 x 10^−4^, which corresponds to a Z-score of 3.81.

### Sliding window tree building

We defined 50KB windows with start sites 25KB apart on the *H. erato demophoon*^24^ chromosomal scale using the *makewindows* tool in bedtools v2.1^79^. We then translated the chromosomal coordinates to scaffold coordinates using *bedtools intersect*, and extracted alignments from each window using *hal2maf* from the haltools suite^75^. We filtered the maf alignments for alignment blocks that were present as a single copy in every species of interest with a custom python script. We further filtered for windows that contained at least 5KB of aligned sites, converted the maf files to fasta with *msa_view*, converted fasta to phylip with PGD Spider v2.1.1.3^80^ and constructed maximum likelihood trees with PhyML, using default parameters ^81^. After obtaining a single best tree for each qualifying window, we categorized them by topology with a custom python script and visualized the results in R using the package *karyoploteR*^82^.

### PhyloNet

Due to computational limits, we were unable to analyze the whole *Heliconius* data set at once. We therefore divided our analysis into the *erato* and *melpomene* clades. When analyzing the *erato* clade, we used the *H. erato demophoon* genome^24^ as our reference, and when analyzing the *H. melpomene* clade, we used Hmel2 as our reference^22^. We defined 10KB windows, spaced 50KB apart using the *makewindows* tool in bedtools v2.1. We then extracted MAF alignments of each region from the whole-genome alignment using the tool *hal2maf*. We considered only single-copy regions, and filtered for windows with at least 1000 bp aligned among all species. This resulted in 5445 usable *erato* clade windows and 3208 *melpomene* clade windows. Because of computational constraints, we were only able to include a subset of the alignments in a single run of the PhyloNet v3.6.7 program MCMC_Seq. We therefore ran 100 iterations of the program, each time sampling 100 random loci without replacement. We used the default parameters, running a 10,000,000-iteration chain with a 2,000,000-iteration burn-in, sampling every 5,000 iterations. We also included two hot-chains of scale 2.0 and 3.0. We generated consensus networks from these results in a method fully outlined in Supplementary Information Section 6.

### Topology distribution

Trees were inferred as described in “*Sliding window tree building*”, except that window size was reduced to 10KB, and windows were non-overlapping. To determine the fraction of each tree topology on each chromosome, we divided the number of windows that recovered the given topology by the total number of windows that successfully resolved any topology on that chromosome. For the distance to end of chromosome, we used the center of each window as the reference point and calculated the number of base pairs between that reference and the nearest edge. We then divided each chromosome into 10 equally-sized bins and calculated the fraction of each bin that recovered each topology in each chromosome independently. We assigned each of our 10KB windows a local recombination rate^24^ by intersecting the window positions with the recombination map using bedtools ^79^. We then split the windows into bins based on their local recombination rate, and calculated the fraction of each bin that recovered each topology, again evaluating each chromosome separately. Finally, we used the gene annotations from *H. erato demophoon* v1^24^ to determine the number of coding bases in each 10KB window. We found that the number of coding bases per window was distributed exponentially, so we log-transformed the data before dividing it into bins. We then determined the topology distributions per bin per chromosome as above.

### Recombination rate estimates

We estimated fine-scale variation in recombination rate along the *H. erato demophoon* chromosomes from linkage-disequilibrium in population genetic data using LDhelmet v1.7^83^. Genotypes were called from whole genome resequencing data of ten *H. erato demophoon* individuals from Panama obtained from Van Belleghem et al. 2017^24^ using the Genome Analysis Tool Kit Haplotypecaller^84^ with default parameters. Individuals’ genotypes were subsequently phased along each *H. erato demophoon* reference genome scaffold using Beagle v4.1 ^85^ with default parameters. For further analysis, phased genotypes were retained only if they had a minimum depth (DP) ≥ 10, maximum depth (DP) ≤ 100 (to avoid false SNPs due to mapping in repetitive regions), and for variant calls, a minimum genotype quality (GQ) ≥ 30. Next, from the phased genotypes, fasta sequences were generated for 50KB windows. These 50KB windows were transformed to haplotype configuration files with the recommended window size of 50 SNPs used by LDhelmet to estimate composite likelihoods of the recombination rate. From the haplotype configuration files, lookup tables for two-locus pairwise recombination likelihoods and Padé coefficients were generated within the recommended value range. Transition matrices were calculated for each chromosome separately by comparing genotypes obtained from *H. erato demophoon* to the outgroup species *H. ricini*, *H. sara*, *H. telesiphe*, *H. clysonymous*, *H. hortense*, *H. hermathena* and *H. charithonia*. The likelihood lookup tables, Padé coefficients and transition matrices were used in the rjMCMC procedure of LDhelmet to estimate the recombination map. In this latter step, 1,000,000 Markov chain iterations were run with a burn-in of 100,000 iterations, a window size of 50 SNPs and block penalty of 50. For convenience, recombination rate estimates (ρ) were converted to cM/Mb by scaling values for each chromosome according to the map length of each chromosome as obtained from pedigree data.

### Triplet Gene Tree Test

We first calculated gene trees in 10KB windows as described above. To ensure each tree was independent, we only considered one window every 50KB. We then split each full gene tree into its component triplet topologies, determined the outgroup for each triplet, and calculated the internal branch length. We next grouped each set of branch lengths by triplet and by outgroup, and for each subset determined the likelihood that the branch lengths were best described by a simple exponential distribution as expected under ILS or a mixture of ILS and either introgression or speciation processes. The full method is detailed in Supplementary Information Section 9.

### Mapping inversions

We first identified regions of interest (ROI) based on the coordinates of the first and last windows that displayed the altered tree topology. We then used the command *halLiftover* from the haltools suite to identify the contigs of each species that mapped to the ROI. We used a custom python script to extract the relative positions of each alignment block on each contig and filtered for those that had segments aligned in both directions in the region of interest. We inspected these candidate contigs manually with IGV^86^ and identified those in which the beginning of the contig mapped in one direction to one side of the ROI, and the end mapped in the opposite direction to the other side of the ROI. We used the coordinates where the contig alignments terminated as candidate breakpoints, and manually inspected all contig alignments for those that mapped across the breakpoints in a single direction.

## Supporting information

## Data Availability

On publication, reads generated in this study will have been deposited in the SRA. IDs TBD. Genome assemblies generated in this study are available on LepBase (http://lepbase.org). Upon publication, all other data will be available on Dryad.

## Code Availability

Upon publication, all code used to analyze data will be available on the github repositories github.com/nbedelman/HeliconiusComparativeGenomics (general analyses), github.com/nbedelman/ScaffoldingWithDiscovar (SWD pipeline) github.com/michaelmiyagi (triplet gene tree test).

## Author Contributions

N.B.E., N.P., D.E.N., R.C., S.K., M.B., R.D.R., K.K.D., M.K., M.J., C.D.J, W.O.M, D.J., and J.M. conceived the project; N.B.E., P.B.F., B.C., D.J., and J.M. designed experiments; J.D., G.M., C.S., M.C., R.D.R., K.K.D., M.J., C.D.J., W.O.M., and J.M. supplied specimens and extracted DNA; D.J., B.C., G.G.A., and F.D.P. assembled genomes, P.B.F. inferred species trees; N.B.E. and R.D. performed evolutionary rate analysis; N.B.E performed reference gap filling, phylogenetic network, local gene tree, and genome structure analyses; J.D. and S.V.B. generated linkage maps, M.M., J.W., and A.B. developed gene tree test; N.B.E. and J.M. wrote and edited the manuscript with input from all authors.

## Acknowledgments

This project was funded by a SPARC Grant from the Broad Institute of Harvard and MIT and startup and studentship funds from Harvard University. We thank the Harvard FAS Research Computing team for their support. We also thank Jonathan Edelman for assistance with the Scaffolding With Discovar pipeline and Jiafan Zhu and Luay Nakhleh for their guidance with the PhyloNet toolkit. We also thank Eadaoin Harney for assistance with qpGraph, as well as Neil Rosser, Tianzhu Xiong, and Simon Martin for illuminating discussions. LepBase development was supported by BBSRC grant BB/K020161/1.

## Works Cited

1. Schluter, D. The Ecology of Adaptive Radiation. (OUPO Oxford, 2000).

2. Gillespie, R. Community assembly through adaptive radiation in Hawaiian spiders. Science 303, 356–359 (2004).

3. Moore, W. & Robertson, J. A. Explosive adaptive radiation and extreme phenotypic diversity within ant-nest beetles. Curr. Biol. 24, 2435–2439 (2014).

4. Jønsson, K. A. et al. Ecological and evolutionary determinants for the adaptive radiation of the Madagascan vangas. Proc. Biol. Sci. 109, 6620–6625 (2012).

5. Lamichhaney, S. et al. Evolution of Darwin’s finches and their beaks revealed by genome sequencing. Nature 518, 371–375 (2015).

6. Kozak, K. M. et al. Multilocus species trees show the recent adaptive radiation of the mimetic Heliconius butterflies. Syst. Biol. 64, 505–524 (2015).

7. Gilbert, L. E. in Coevolution of Animals and Plants (ed. Gilbert, L. E.) 210–240 (1975).

8. Brown, K. S. The biology of Heliconius and related genera. Annu. Rev. Entomol. 26, (1981).

9. Jiggins, C. D. The Ecology and Evolution of Heliconius Butterflies. (Oxford University Press, 2017).

10. The Heliconius Genome Consortium. Butterfly genome reveals promiscuous exchange of mimicry adaptations among species. Nature 487, 94–98 (2012).

11. Pardo-Diaz, C. et al. Adaptive Introgression across Species Boundaries in Heliconius Butterflies. PLoS Genet. 8, e1002752–13 (2012).

12. Jay, P. et al. Supergene evolution triggered by the introgression of a chromosomal inversion. Curr. Biol. 28, 1839–1845.e3 (2018).

13. Zhang, W., Dasmahapatra, K. K., Mallet, J., Moreira, G. R. P. & Kronforst, M. R. Genome-wide introgression among distantly related Heliconius butterfly species. Genome Biol. 17, 25 (2016).

14. Kozak, K. M., McMillan, O., Joron, M. & Jiggins, C. D. Genome-wide admixture is common across the Heliconius radiation. bioRxiv 414201 (2018). doi: 10.1101/414201

15. Martin, S. H. et al. Genome-wide evidence for speciation with gene flow in Heliconius butterflies. Genome Research 23, 1817–1828 (2013).

16. Martin, S. H., Davey, J., Salazar, C. & Jiggins, C. Recombination rate variation shapes barriers to introgression across butterfly genomes. bioRxiv 297531 (2018). doi: 10.1101/297531

17. Davey, J. W. et al. No evidence for maintenance of a sympatric Heliconius species barrier by chromosomal inversions. Evolution Letters 1, 138–154 (2017).

18. Clavijo, B. et al. W2RAP: a pipeline for high quality, robust assemblies of large complex genomes from short read data. bioRxiv 110999 (2017). doi: 10.1101/110999

19. Weisenfeld, N. I. et al. Comprehensive variation discovery in single human genomes. Nat. Genet. 46, 1350–1355 (2014).

20. Broad Institute. DISCOVAR: Assemble genomes, find variants. https://www.broadinstitute.org/software/discovar/blog (2015). Available at: (Accessed: 29 April 2016)

21. Love, R. R., Weisenfeld, N. I., Jaffe, D. B., Besansky, N. J. & Neafsey, D. E. Evaluation of DISCOVAR de novo using a mosquito sample for cost-effective short-read genome assembly. BMC Genomics 17, 187–10 (2016).

22. Davey, J. W. et al. Major improvements to the Heliconius melpomene genome assembly used to confirm 10 chromosome fusion events in 6 million years of butterfly evolution. G3 6, 695–708 (2016).

23. Simão, F. A., Waterhouse, R. M., Ioannidis, P., Kriventseva, E. V. & Zdobnov, E. M. BUSCO: assessing genome assembly and annotation completeness with single-copy orthologs. Bioinformatics 31, 3210–3212 (2015).

24. Van Belleghem, S. M. et al. Complex modular architecture around a simple toolkit of wing pattern genes. Nat. Ecol. Evol. 1, 52 (2017).

25. Ahola, V. et al. The Glanville fritillary genome retains an ancient karyotype and reveals selective chromosomal fusions in Lepidoptera. Nat. Commun. 5, 4737 (2014).

26. Zhan, S. et al. The genetics of monarch butterfly migration and warning colouration. Nature 514, 317–321 (2014).

27. Nowell, R. W. et al. A high-coverage draft genome of the mycalesine butterfly Bicyclus anynana. Gigascience 6, 1–7 (2017).

28. Nishikawa, H. et al. A genetic mechanism for female-limited Batesian mimicry in Papilio butterfly. Nat. Genet. 47, 405–409 (2015).

29. Cong, Q., Borek, D., Otwinowski, Z. & Grishin, N. V. Skipper genome sheds light on unique phenotypic traits and phylogeny. BMC Genomics 16, 639 (2015).

30. The International Silkworm Genome Consortium. The genome of a lepidopteran model insect, the silkworm Bombyx mori. Insect Biochem. Mol. Biol. 38, 1036–1045 (2008).

31. You, M. et al. A heterozygous moth genome provides insights into herbivory and detoxification. Nat. Genet. 45, 220–225 (2013).

32. Brower, A. V. Z. & Orduña, I. J. G. Missing data, clade support and ‘reticulation’: the molecular systematics of Heliconius and related genera (Lepidoptera: Nymphalidae) reexamined. Cladistics 34, 151–166 (2018).

33. Green, R. E. et al. A Draft Sequence of the Neandertal Genome. Science 328, 710–722 (2010).

34. Patterson, N. et al. Ancient admixture in human history. Genetics 192, 1065–1093 (2012).

35. Than, C., Ruths, D. & Nakhleh, L. PhyloNet: a software package for analyzing and reconstructing reticulate evolutionary relationships. BMC Bioinformatics 9, 322 (2008).

36. Wen, D., Yu, Y. & Nakhleh, L. Bayesian inference of reticulate phylogenies under the multispecies network coalescent. PLoS Genet. (2016). doi:10.1371/journal.pgen.1006006

37. Mallet, J. Gregarious roosting and home range in Heliconius butterflies. Natl. Geogr. Res. 2, 198–215 (1986).

38. Montgomery, S. H., Merrill, R. M. & Ott, S. R. Brain composition in Heliconius butterflies, posteclosion growth and experience-dependent neuropil plasticity. J. Comp. Neurol. 524, 1747–1769 (2016).

39. Coyne, J. A. & Orr, H. A. Speciation. (Sinauer Associates Incorporated, 2004).

40. Dobzhansky, T. Genetics and the Origin of Species. (Columbia University Press, 1937).

41. Begun, D. J. & Aquadro, C. F. Levels of naturally occurring DNA polymorphism correlate with recombination rates in D. melanogaster. Nature 356, 519–520 (1992).

42. Schumer, M. et al. Natural selection interacts with recombination to shape the evolution of hybrid genomes. Science 360, eaar3684–660 (2018).

43. Orr, H. A. & Turelli, M. Dominance and Haldane’s rule. Genetics 143, 613–616 (1996).

44. Jiggins, C. D. et al. Sex-linked hybrid sterility in a butterfly. Evolution 55, 1631–1638 (2001).

45. Naisbit, R. E., Jiggins, C. D., Linares, M., Salazar, C. & Mallet, J. Hybrid sterility, Haldane’s rule and speciation in Heliconius cydno and H. melpomene. Genetics 161, 1517–1526 (2002).

46. Van Belleghem, S. M. et al. Patterns of Z chromosome divergence among Heliconius species highlight the importance of historical demography. Mol. Ecol. 27, 3852–3872 (2018).

47. Wickham, H. ggplot2: elegant graphics for data analysis. (Springer, 2016).

48. Nadeau, N. J. et al. The gene cortex controls mimicry and crypsis in butterflies and moths. Nature 534, 106–110 (2016).

49. Joron, M. et al. A conserved supergene locus controls colour pattern diversity in Heliconius butterflies. PLoS Biol. 4, e303–10 (2006).

50. Joron, M. et al. Chromosomal rearrangements maintain a polymorphic supergene controlling butterfly mimicry. Nature 477, 203–206 (2011).

51. Corbett-Detig, R. B., Zhou, J., Clark, A. G., Hartl, D. L. & Ayroles, J. F. Genetic incompatibilities are widespread within species. Nature 504, 135–137 (2013).

52. Presgraves, D. C. A fine-scale genetic analysis of hybrid incompatibilities in Drosophila. Genetics 163, 955–972 (2003).

53. Schumer, M., Cui, R., Powell, D. L., Elife, R. D. 2014. High-resolution mapping reveals hundreds of genetic incompatibilities in hybridizing fish species. eLife

54. Machado, C. A., Haselkorn, T. S. & Noor, M. A. F. Evaluation of the genomic extent of effects of fixed inversion differences on intraspecific variation and interspecific gene flow in Drosophila pseudoobscura and D. persimilis. Genetics 175, 1289–1306 (2007).

55. Fuller, Z. L., Leonard, C. J., Young, R. E., Schaeffer, S. W. & Phadnis, N. Ancestral polymorphisms explain the role of chromosomal inversions in speciation. PLoS Genet. 14, e1007526 (2018).

56. Lamichhaney, S. et al. Structural genomic changes underlie alternative reproductive strategies in the ruff (Philomachus pugnax). Nat. Genet. 48, 84–88 (2016).

57. Svedberg, J. et al. Convergent evolution of complex genomic rearrangements in two fungal meiotic drive elements. Nat. Commun. 9, 4242 (2018).

58. Badouin, H. et al. Chaos of rearrangements in the mating-type chromosomes of the anther-smut fungus Microbotryum lychnidis-dioicae. Genetics 200, 1275–1284 (2015).

59. Purcell, J., Brelsford, A., Wurm, Y., Perrin, N. & Chapuisat, M. Convergent genetic architecture underlies social organization in ants. Curr. Biol. 24, 2728–2732 (2014).

60. Lu, H., Giordano, F. & Ning, Z. Oxford nanopore MinION sequencing and genome assembly. Genomics, Proteomics & Bioinformatics 14, 265–279 (2016).

61. Berlin, K. et al. Assembling large genomes with single-molecule sequencing and locality-sensitive hashing. Nat. Biotechnol. 33, 623–630 (2015).

62. Coombe, L. et al. Assembly of the complete sitka spruce chloroplast genome using 10X genomics’ GemCode sequencing data. PLoS ONE 11, e0163059 (2016).

63. Mapleson, D. et al. KAT: a K-mer analysis toolkit to quality control NGS datasets and genome assemblies. Bioinformatics 33, 574–576 (2017).

64. Luo, R. et al. SOAPdenovo2: an empirically improved memory-efficient short-read de novo assembler. Gigascience 1, 18 (2012).

65. Vurture, G. W. et al. GenomeScope: fast reference-free genome profiling from short reads. Bioinformatics 33, 2202–2204 (2017).

66. Marçais, G. & Kingsford, C. A fast, lock-free approach for efficient parallel counting of occurrences of k-mers. Bioinformatics 27, 764–770 (2011).

67. Smit, A., Hubley, R. & Green, P. 2013-2015. RepeatMasker Open-4.0. (2013).

68. Paten, B. et al. Cactus: Algorithms for genome multiple sequence alignment. Genome Research 21, 1512–1528 (2011).

69. Kent, W. J. et al. The human genome browser at UCSC. Genome Research 12, 996–1006 (2002).

70. Kalyaanamoorthy, S., Minh, B. Q., Wong, T. K. F., Haeseler, von, A. & Jermiin, L. S. ModelFinder: fast model selection for accurate phylogenetic estimates. Nat. Methods 14, 587–589 (2017).

71. Nguyen, L.-T., Schmidt, H. A., Haeseler, von, A. & Minh, B. Q. IQ-TREE: a fast and effective stochastic algorithm for estimating maximum-likelihood phylogenies. Mol. Biol. Evol. 32, 268–274 (2015).

72. Hoang, D. T., Chernomor, O., Haeseler, von, A., Minh, B. Q. & Vinh, L. S. UFBoot2: Improving the ultrafast bootstrap approximation. Mol. Biol. Evol. 35, 518–522 (2018).

73. Mirarab, S. et al. ASTRAL: genome-scale coalescent-based species tree estimation. Bioinformatics 30, i541–8 (2014).

74. Kück, P. & Meusemann, K. FASconCAT: Convenient handling of data matrices. Molecular Phylogenetics and Evolution 56, 1115–1118 (2010).

75. Hickey, G., Paten, B., Earl, D., Zerbino, D. & Haussler, D. HAL: a hierarchical format for storing and analyzing multiple genome alignments. Bioinformatics 29, 1341–1342 (2013).

76. Hubisz, M. J., Pollard, K. S. & Siepel, A. PHAST and RPHAST: phylogenetic analysis with space/time models. Brief. Bioinformatics 12, 41–51 (2011).

77. Conesa, A. et al. Blast2GO: a universal tool for annotation, visualization and analysis in functional genomics research. Bioinformatics 21, 3674–3676 (2005).

78. Page, A. J. et al. SNP-sites: rapid efficient extraction of SNPs from multi-FASTA alignments. Microb. Genom. 2, e000056 (2016).

79. Quinlan, A. R. BEDTools: The Swiss-army tool for genome feature analysis. Curr. Protoc. Bioinformatics 47, 11.12.1–34 (2014).

80. Lischer, H. E. L. & Excoffier, L. PGDSpider: an automated data conversion tool for connecting population genetics and genomics programs. Bioinformatics 28, 298–299 (2012).

81. Guindon, S. et al. New algorithms and methods to estimate maximum-likelihood phylogenies: assessing the performance of PhyML 3.0. Syst. Biol. 59, 307–321 (2010).

82. Gel, B. & Serra, E. karyoploteR: an R/Bioconductor package to plot customizable genomes displaying arbitrary data. Bioinformatics 33, 3088–3090 (2017).

83. Chan, A. H., Jenkins, P. A. & Song, Y. S. Genome-wide fine-scale recombination rate variation in Drosophila melanogaster. PLoS Genet. 8, e1003090 (2012).

84. Van der Auwera, G. A. et al. From FastQ data to high-confidence variant calls: the genome analysis toolkit best practices pipeline. Curr. Protoc. Bioinformatics 43, 11.10.1–10.33 (2013).

85. Browning, B. L. & Browning, S. R. Genotype imputation with millions of reference samples. Am. J. Hum. Genet. 98, 116–126 (2016).

86. Thorvaldsdottir, H., Robinson, J. T. & Mesirov, J. P. Integrative genomics viewer (IGV): high-performance genomics data visualization and exploration. Brief. Bioinformatics 14, 178–192 (2013).

