## Supplementary material for "Genomic architecture and introgression shape a butterfly radiation"

#### Supplementary Information

**Section 1. *DISCOVAR de novo*/w2rap Assembly and Quality**

As stated in the main text, our initial assemblies contained many uncollapsed haplotypes, so that single-copy genomic loci actually appeared multiple times in k-mer plots (Figure S1.1, left side). We used the protocol detailed in the main text methods to collapse those haplotypes in the final assembly (Figure S1.1, right side).

DAS_09-93 - *Heliconius burneyi*

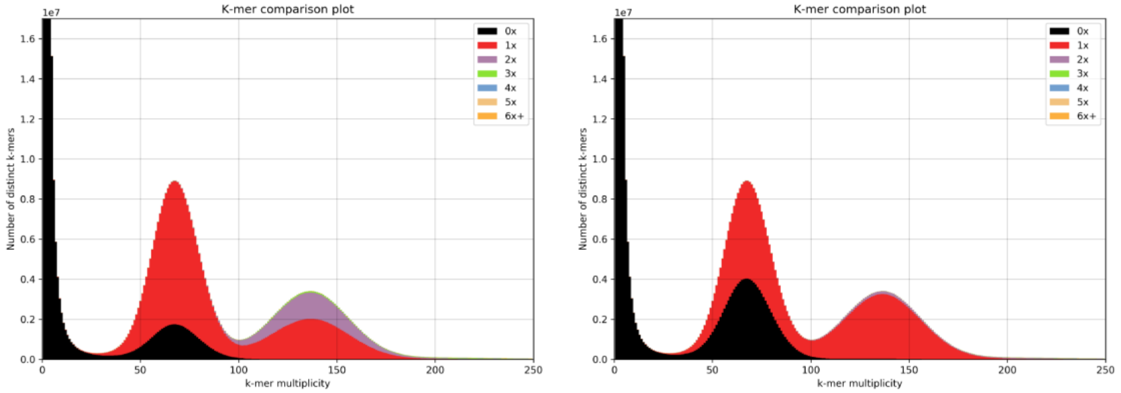

DAS_09-132 - *Eueides tales*

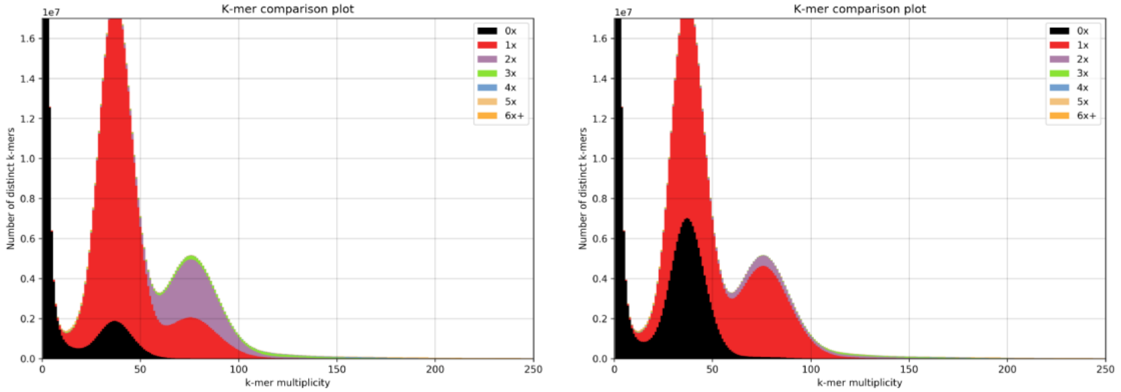

DAS_09-229 - *Heliconius telesiphe* (contaminated)

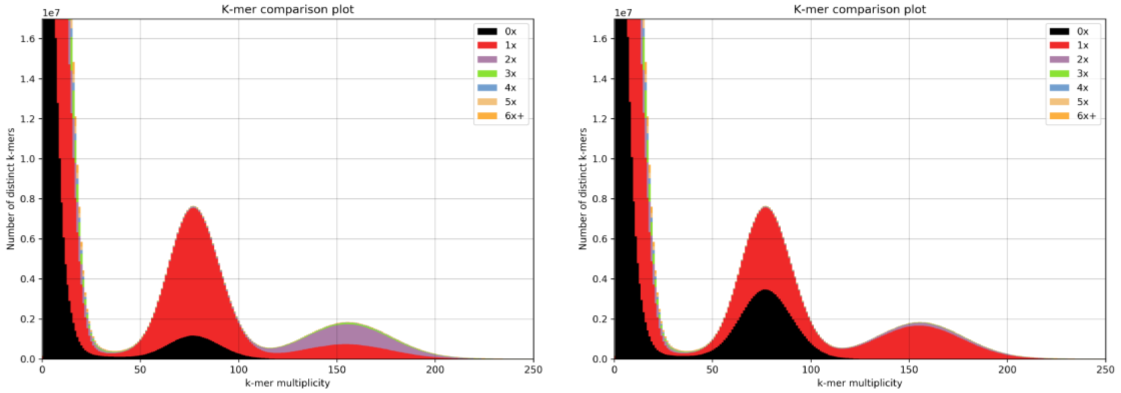

#### DAS_11-919 - *Heliconius telesiphe*

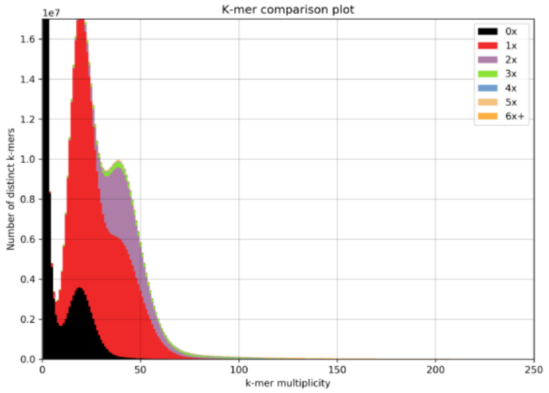

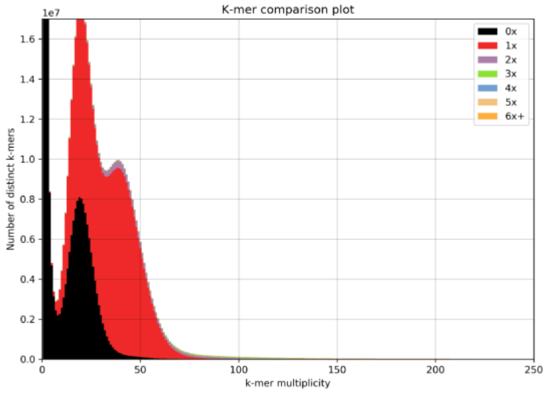

#### CAM-008802 - *Heliconius sara*

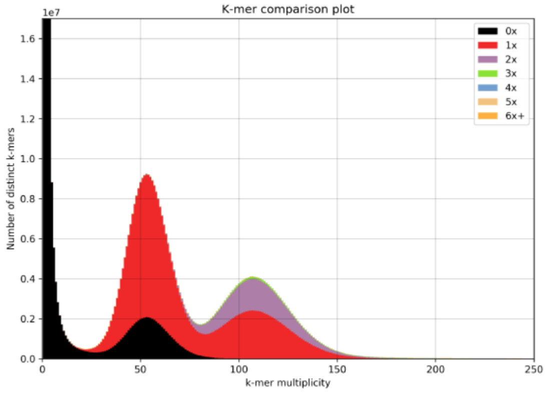

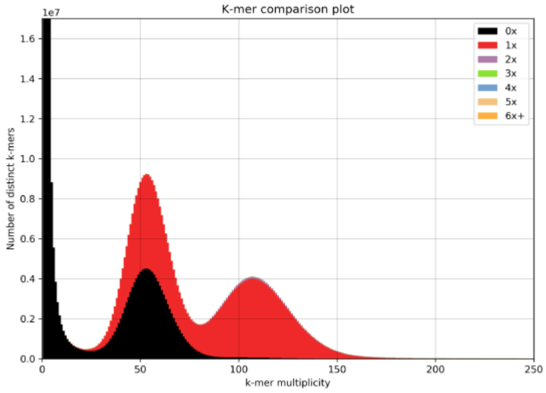

#### DAV_3 - *Heliconius hecale* (replaced)

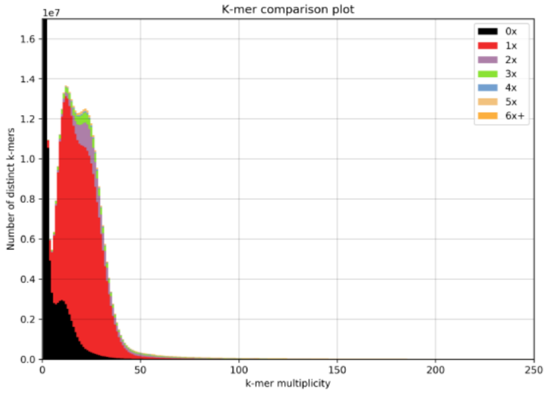

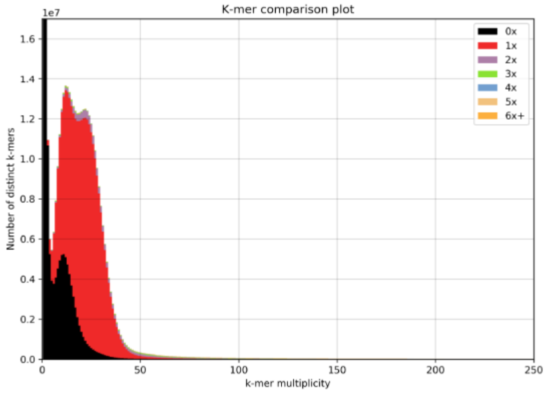

MCM_6897 - *Heliconius doris*

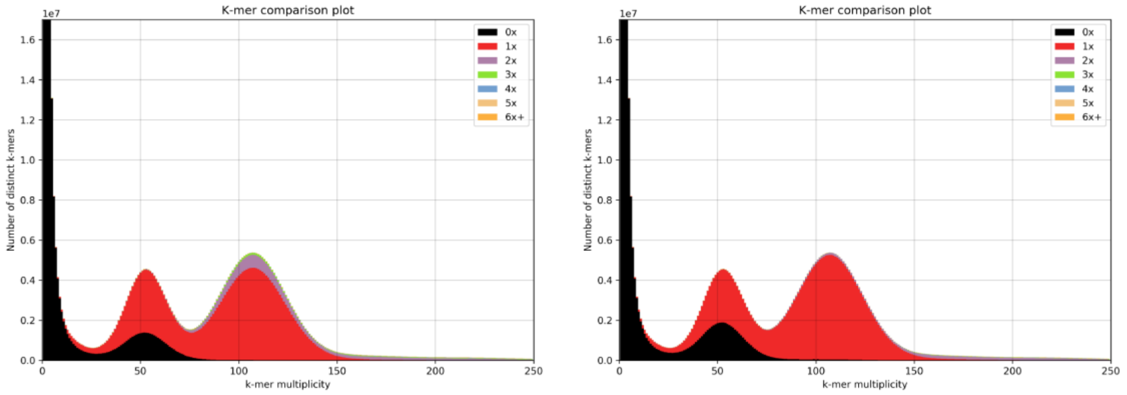

CAM-002492 - *Heliconius hecalesia*

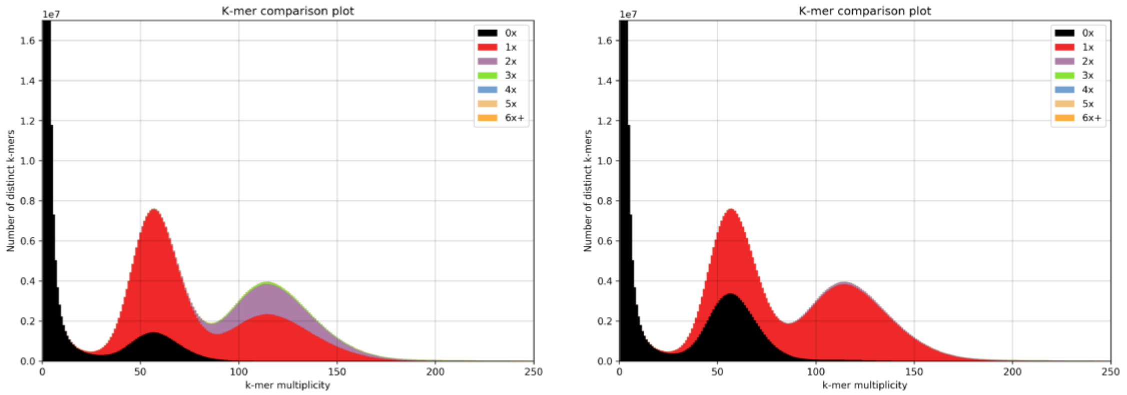

DAS_11-759 *Dryas iulia*

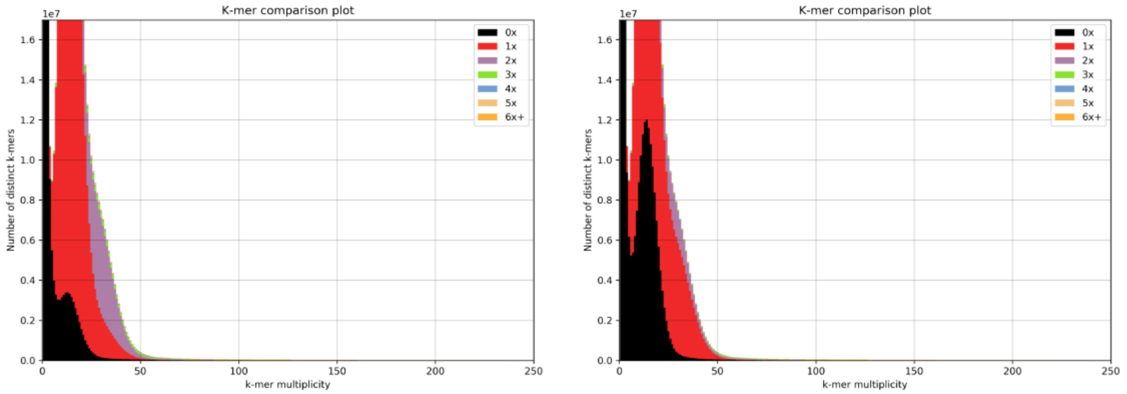

MCM_6900 - *Heliconius himera*

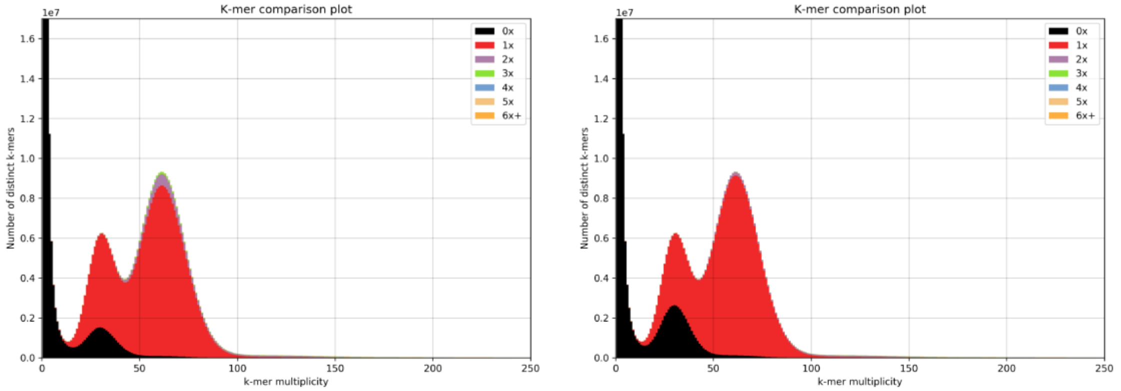

MCM_7203 - *Heliconius hecale*

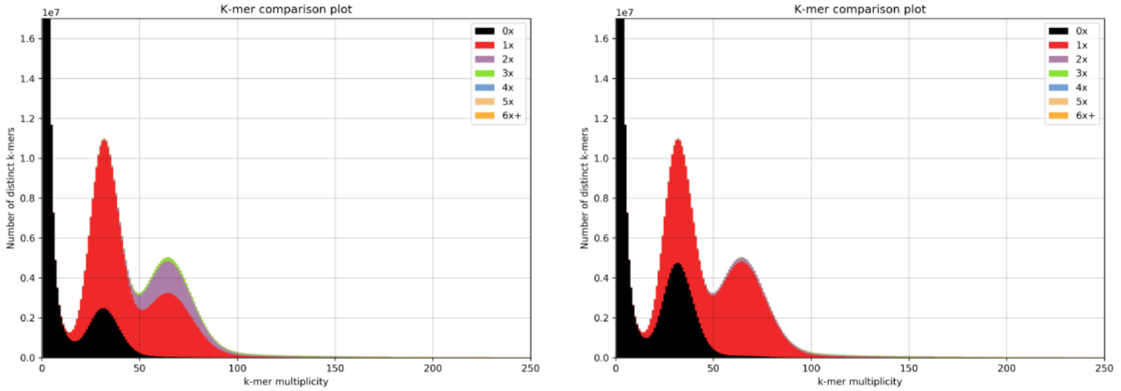

MCM_6700 - *Heliconius erato* x *himera* F1

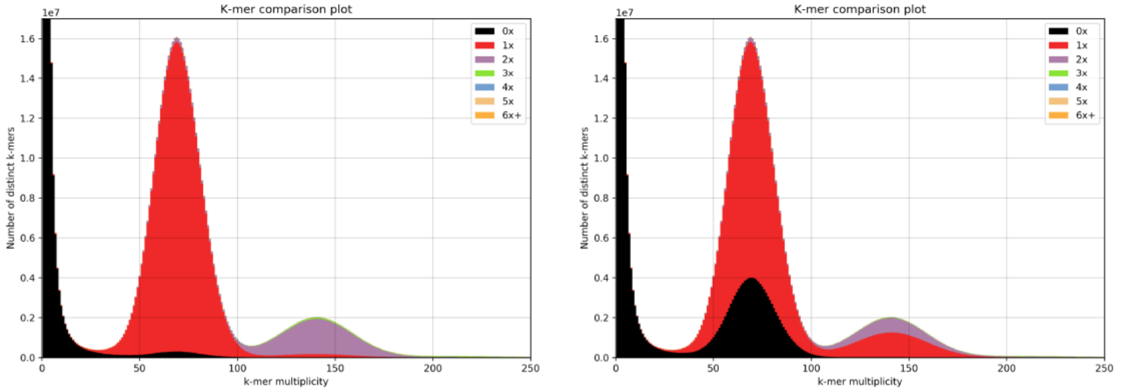

REE *Agraulis vanillae*

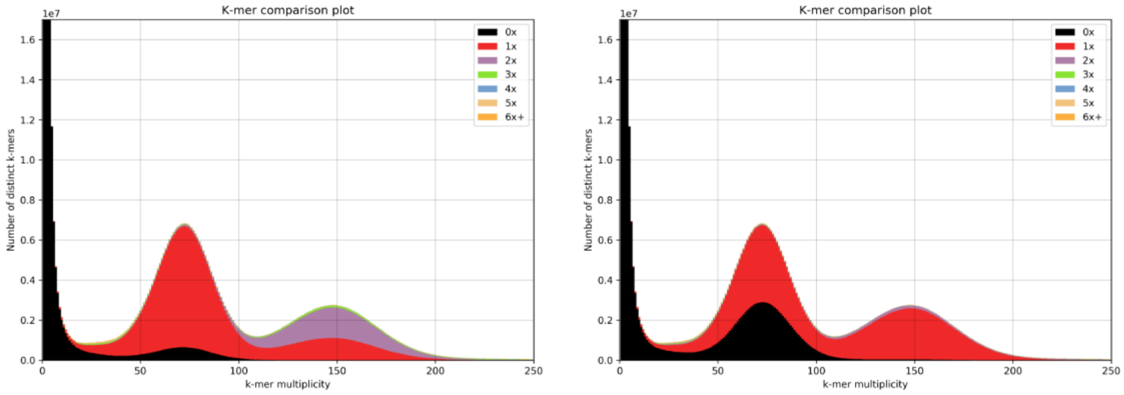

MCM_6394 - *Heliconius erato* mother

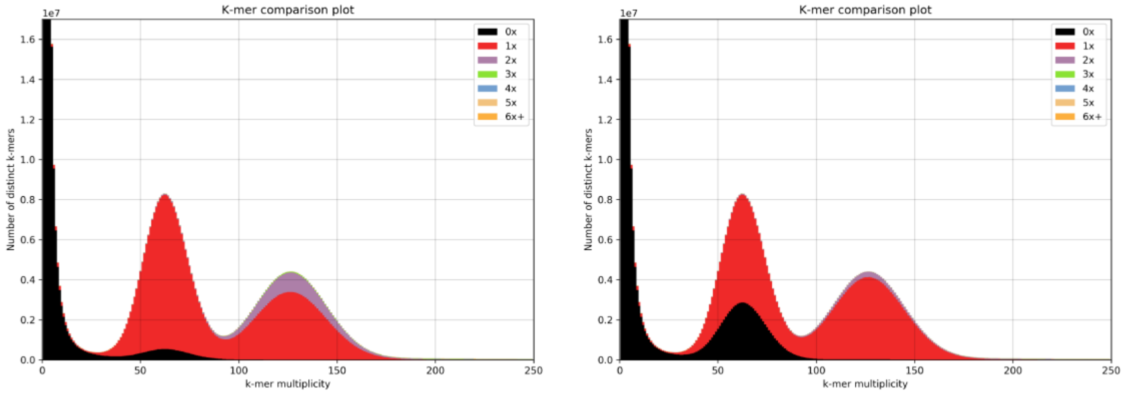

MCM_6363 - *Heliconius himera* father

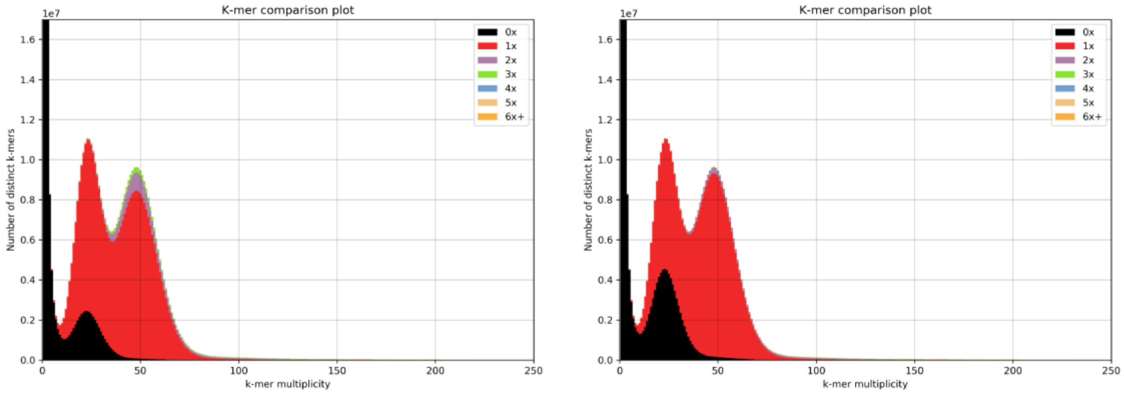

#### DAV_2 - *Heliconius cydno*

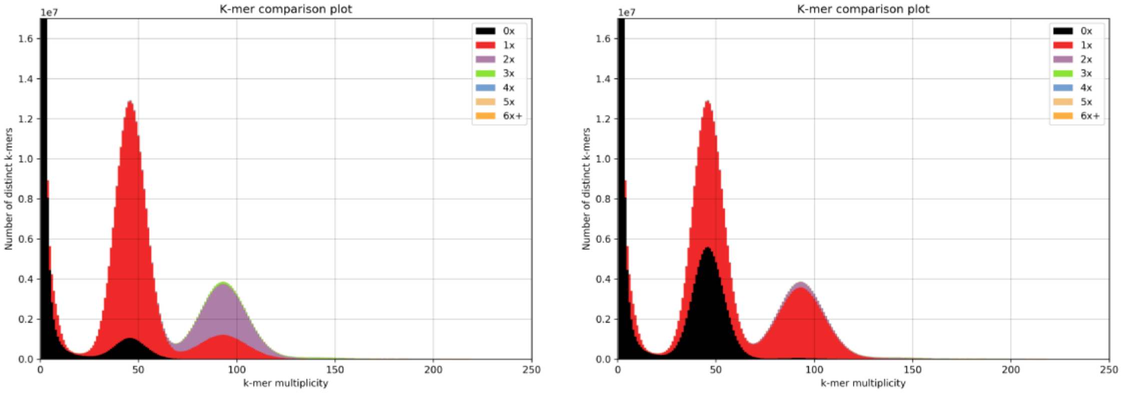

#### DAV_4 - *Heliconius numata*

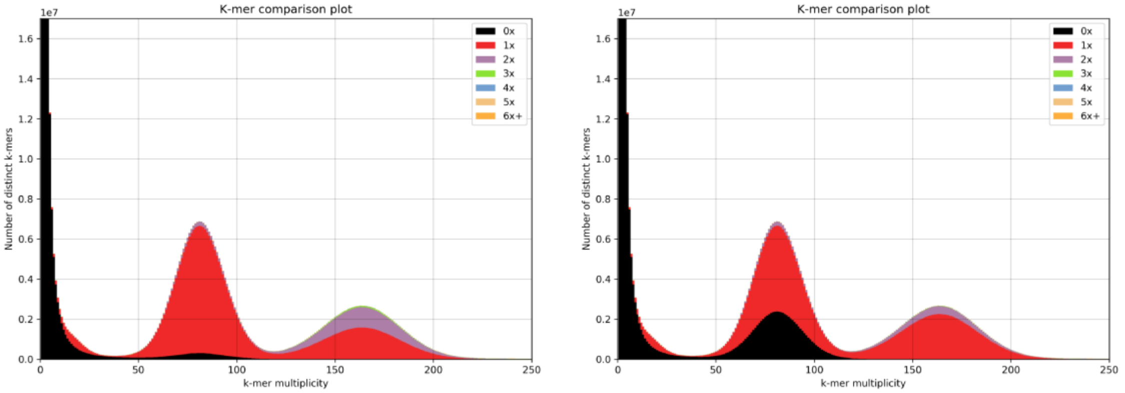

#### DAV_5 - *Heliconius melpomene*

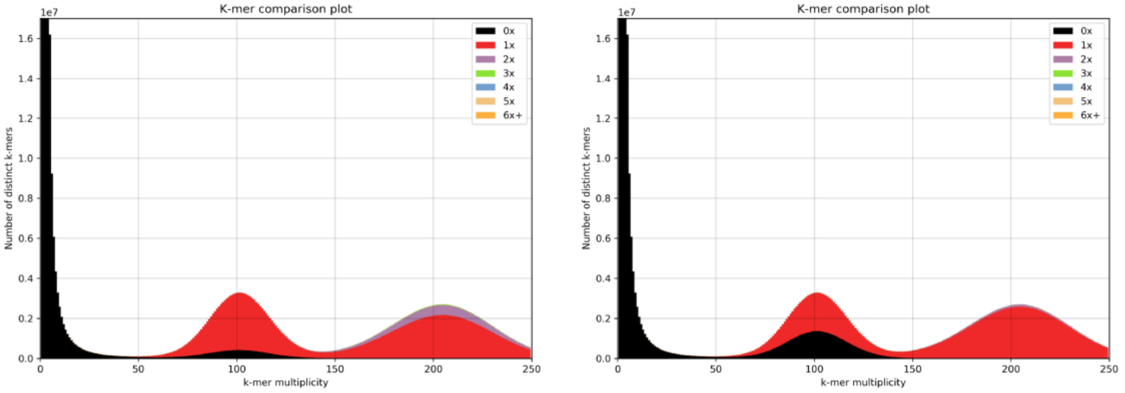

DAS_GELE - *Heliconius elevatus*

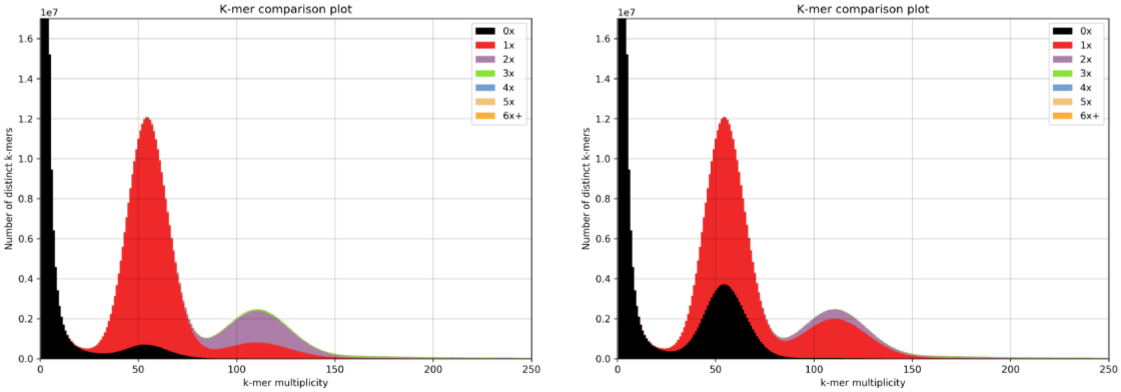

DAS_REL13_139 - *Heliconius pardalinus*

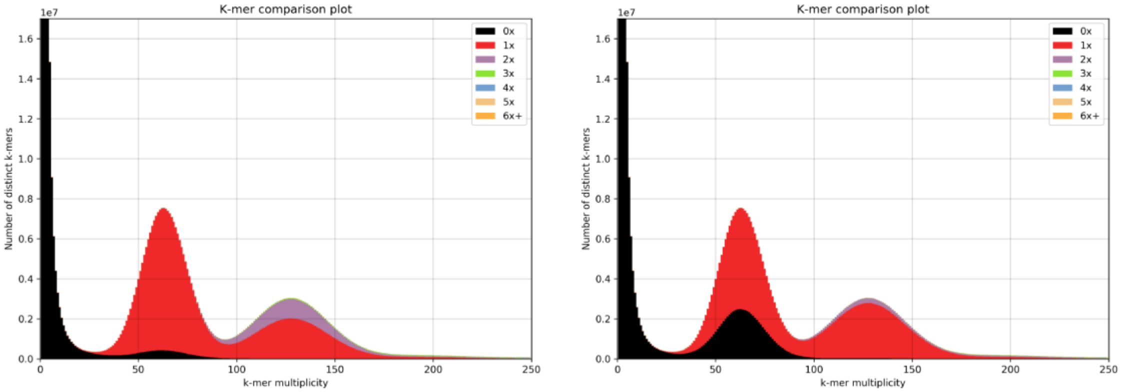

DAV_1 - *Heliconius timareta*

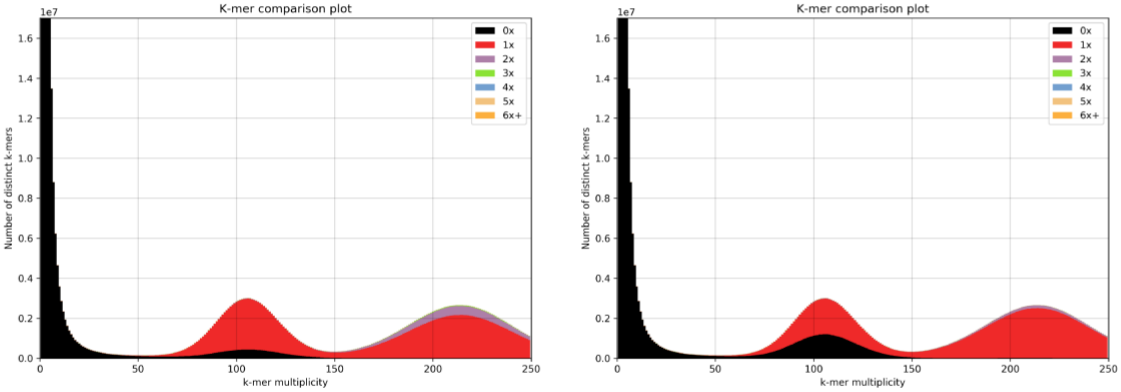

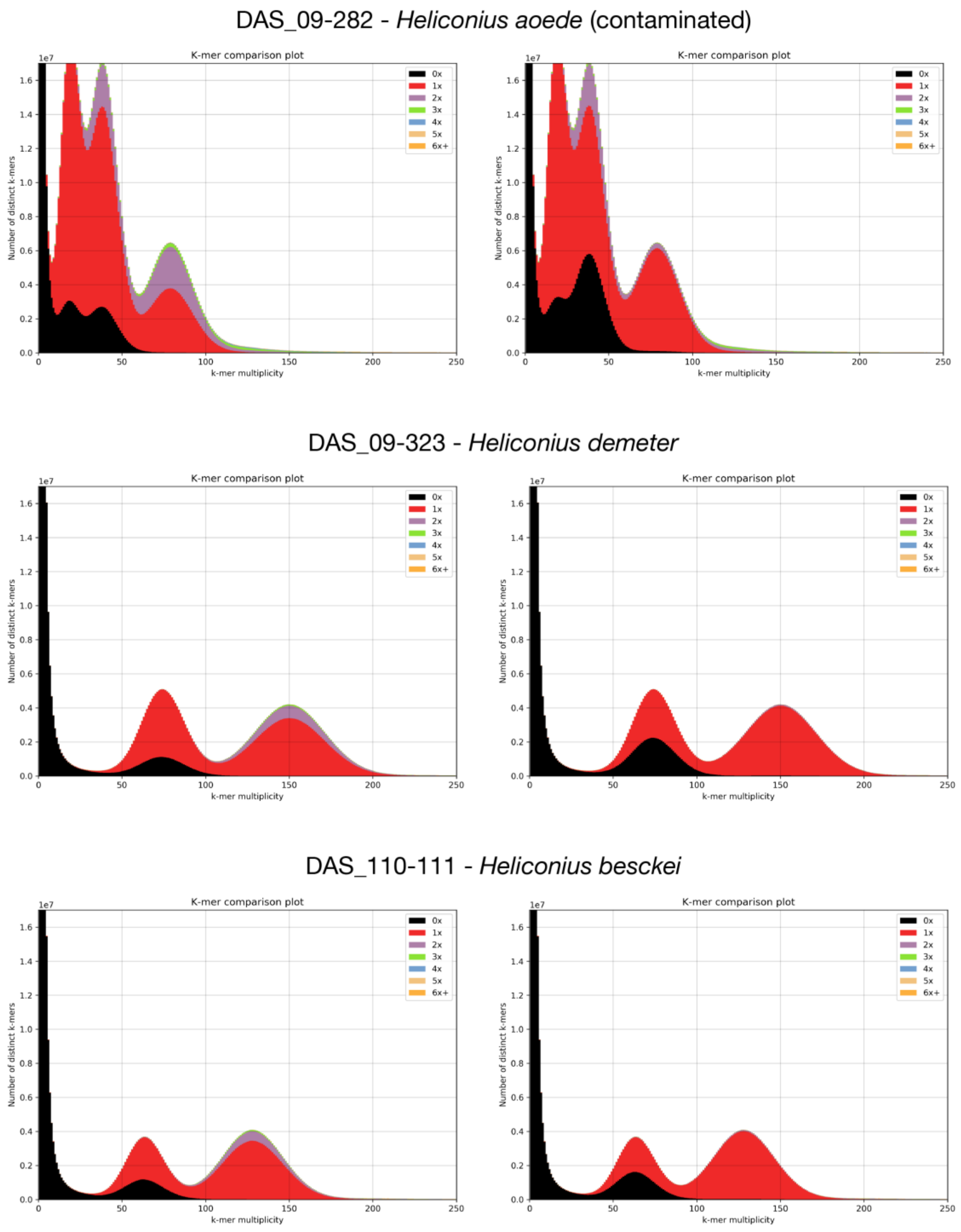

**Figure S1.1. K-mer spectra for all de novo assemblies.** *The x-axis of each plot corresponds to the multiplicity of distinct k-mers in the raw reads. The first peak, near zero, corresponds to contaminants, the second peak corresponds to heterozygous content, and the third to homozygous content. The colors represent the number of times those k-mers are present in the genome assembly. Ideally, none of the contaminants would be assembled, half of the heterozygous content would be present as a single copy, and all of the homozygous content would be present as a single copy. For each species, the plot on the left shows the initial assembly, and the plot on the right shows the final assembly.*

We assessed the quality of our *de novo* assembled genomes by calculating standard metrics including N50, number of contigs, repeat content, and gene content. These values can be found in Table S1.2. N50 values reported here are relatively low compared to published *Heliconius* genomes due to the fact that they were assembled solely using short-read Illumina sequencing, and the number of contigs is relatively large for the same reason. Both of these metrics were quite variable, with N50 values ranging from 21,413 to 106,325 and number of contigs ranging from 17,678 to 62,414. We investigated whether the variation in contiguity could be explained by repeat content by evaluating each *de novo* genome with RepeatMasker[1]. The distribution of repeat content appeared bimodal, with most *melpomene* clade species containing roughly 15% masked bases and most *erato* clade species containing 18-22% masked bases. However, there was no correlation between repeat content and N50 (Figure S1.2).

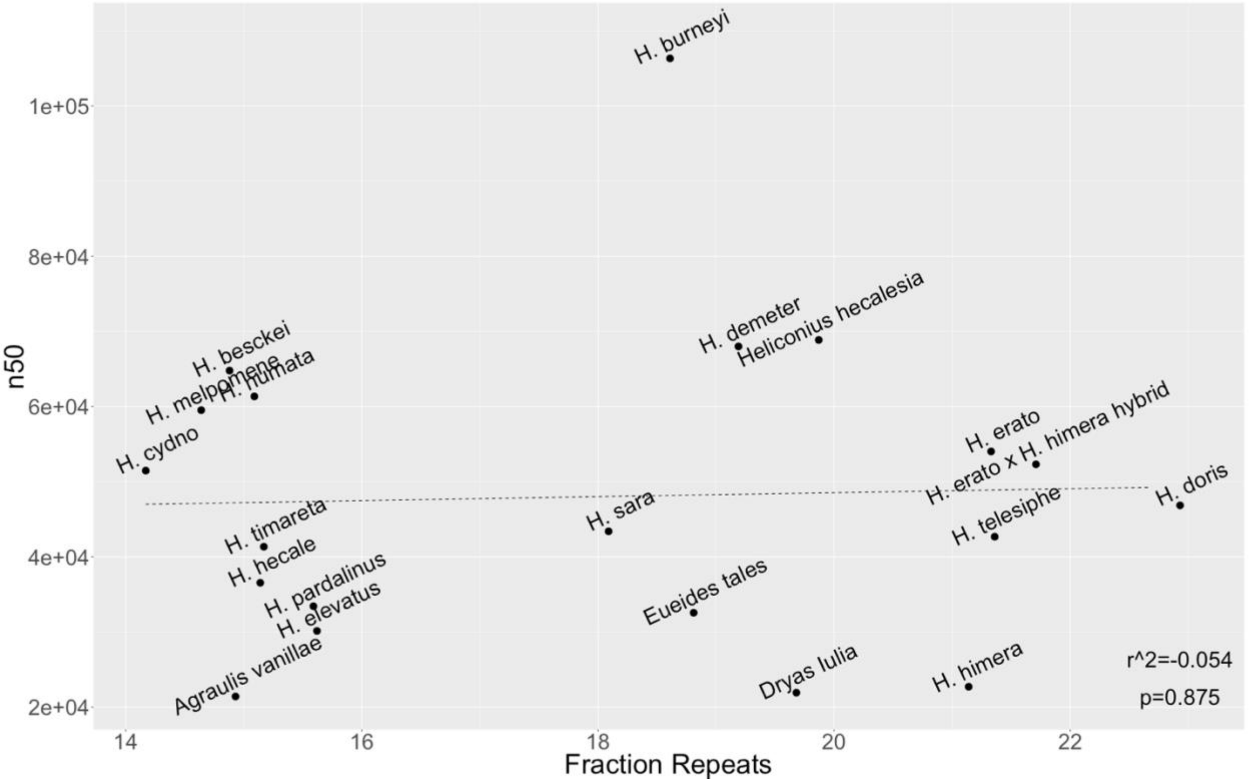

**Figure S1.2: Relationship of N50 to repeat content**

Another possible factor contributing to variation in contiguity is heterozygosity. Heterozygosity may have been especially important, as *Heliconius* species generally have 1-3% heterozygous sites, and most of our specimens were wild-caught rather than inbred. We estimated heterozygosity using the k-mer based GenomeScope method[2], but found no correlation between heterozygosity and N50 value (Figures S1.3). The only variable we tested that was correlated with N50 was contig number which, if genome sizes are comparable, is tautological (Figure S1.4).

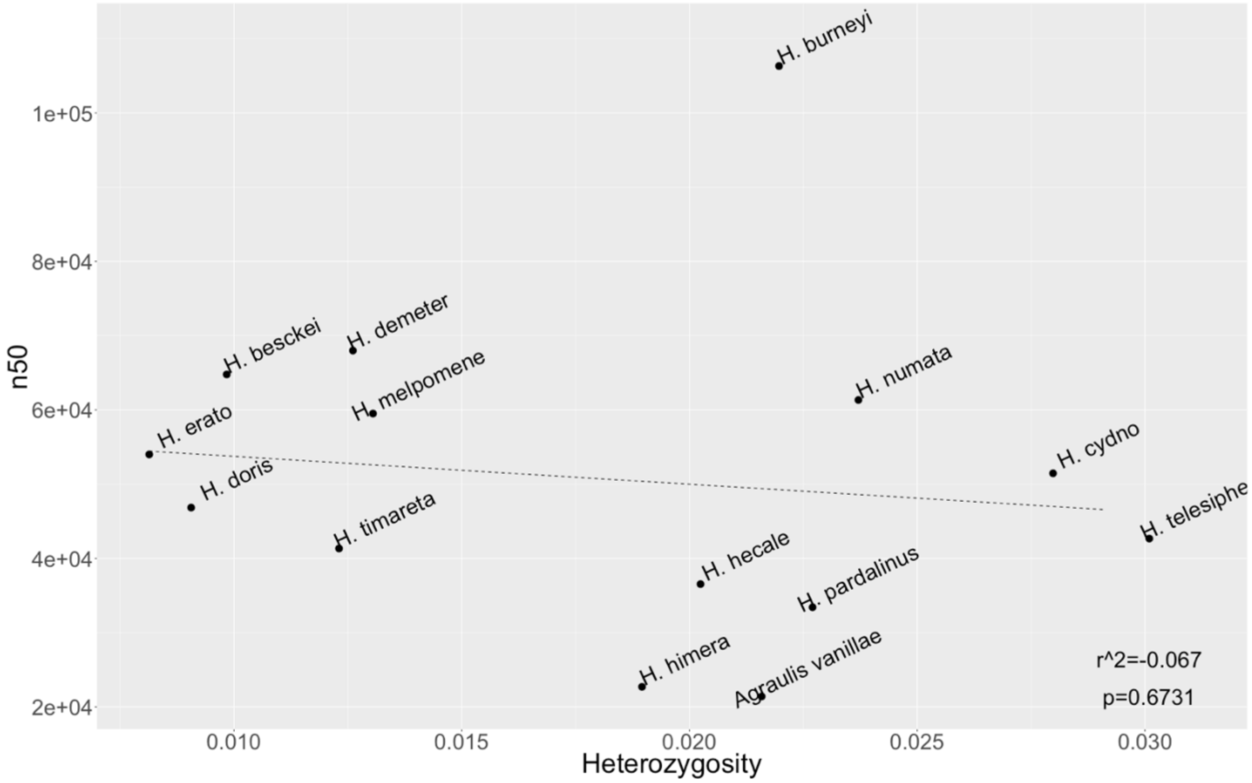

**Figure S1.3: Relationship of N50 to heterozygosity**

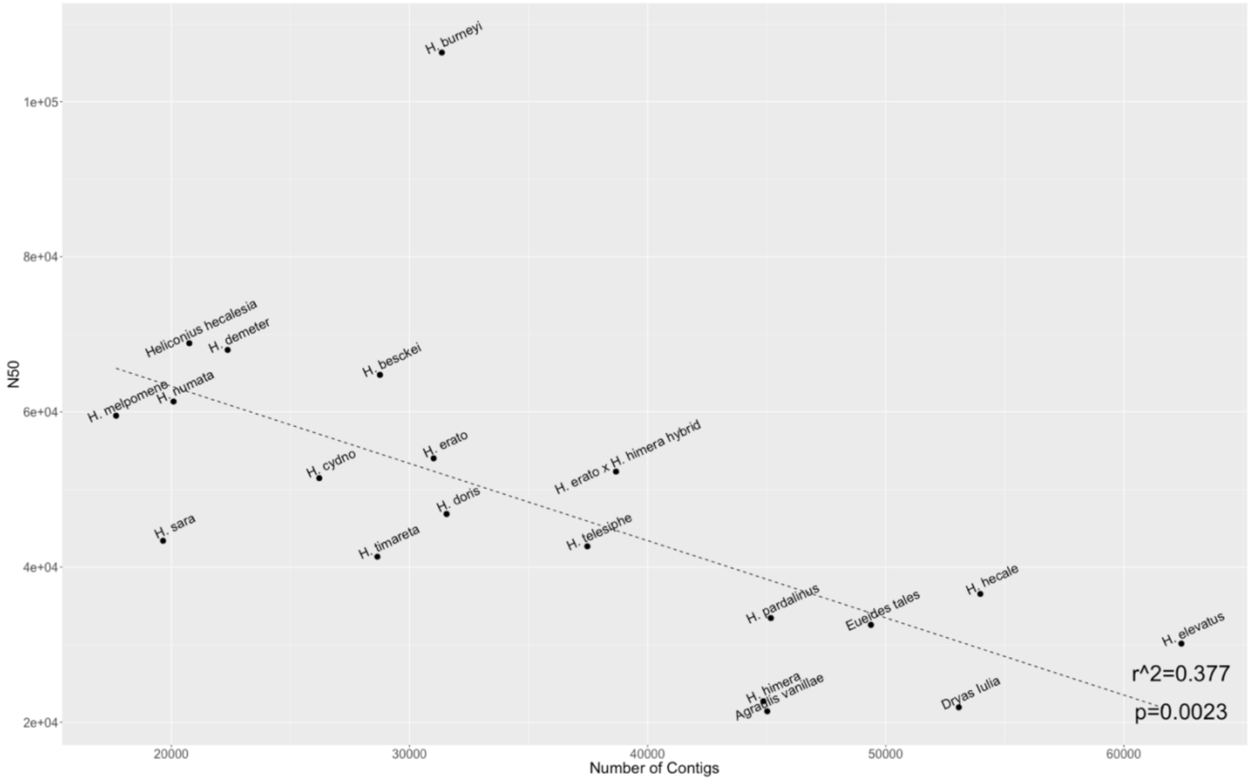

**Figure S1.4: Relationship of N50 to number of contigs**

We further assessed the quality of the genome assemblies by aligning the *H. melpomene* and *H. erato* w2rap genomes to their respective reference assemblies[3,4]. We examined the coverage of the w2rap assembly on the previously published reference and vice versa and found that on average the w2rap assemblies covered 94 percent of the reference genomes, while the reference genomes covered 91 percent of the w2rap genomes (Table S1.3). The discrepancy is likely due in large part to a difference in total assembly size. The *H. melpomene* w2rap assembly is 14MB (5.2%) larger than Hmel2, and the *H. erato* w2rap assembly is 58MB (15%) larger than the *H. erato demophoon* reference. This difference cannot be explained purely by overrepresentation of repetitive elements, as the overall proportion of the genome identified by RepeatMasker as repeats is almost identical between the conspecific assemblies[1]. It is more likely that the larger w2rap genome sizes are caused by uncollapsed haplotypes due to stringent thresholds employed in w2rap for collapsing heterozygous regions. This hypothesis is further supported by the observation that the F1 interspecific hybrid individual (*H. erato/H. himera* hybrid) has by far the largest total assembly size among *Heliconius* species and by BUSCO analysis showing slight increases in duplicated genes relative to high quality lepidopteran assemblies, as well as to other w2rap assemblies (Figure S1.5) [5].

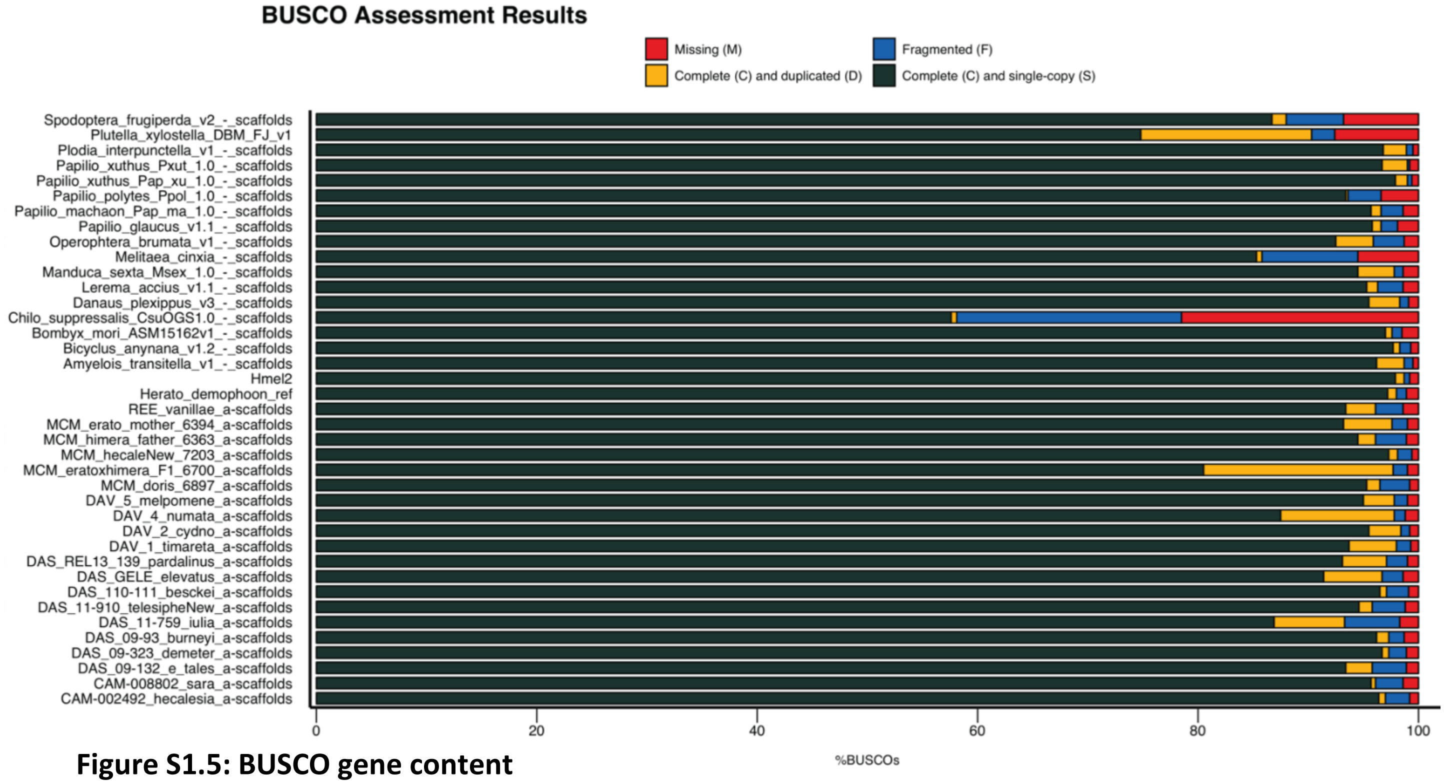

**Figure S1.5: BUSCO gene content**

*This plot displays BUSCO results from all de novo genomes assembled here, as well as other lepidopteran genomes obtained from lepbase.org. All bars below "*Herato_demophoon_ref*" resulted this study. Gene content is comparable between the w2rap assemblies and other lepidopteran assemblies, though the percentage of complete genes identified in w2rap genomes is slightly lower, and percent duplicated slightly higher, than the highest ranking Lepidoptera genomes.*

Apart from genomic coverage, we also queried whether w2rap contigs contained assembly errors by mapping all scaffolds greater than 1000 bp of the *H. melpomene de novo* genome onto its conspecific reference. In order to assess alignment quality, we scored each contig according to the following scale. Colors in parentheses correspond to those in figures S1.6-S1.8:

*Unique Map* (dark green): The entire w2rap contig maps as a block to a single location in the genome.
*All Inside* (light green): The alignment of the contig may be broken into multiple blocks, but more than 95% of the contig maps to a single location defined by projecting the ends of the contig out from its highest quality alignment block.
*>75% inside* (yellow): More than 75%, but less than 95% of the contig maps to a single location as defined above.
*>50% inside* (orange): More than 50%, but less than 75% of the contig maps to a single location as defined above.
*>50% Scaffold* (pink): Fewer than 50% of the contig maps to a single location, but more than 50% of the contig maps to a single scaffold.
*<50% Scaffold* (red): There is no scaffold in which more than 50% of the contig maps.
*2X Duplicated* (sky blue): entire contig maps to exactly 2 locations in genome.
*<=5X Duplicated* (blue): entire contig maps to 3-5 locations in genome.
*>5x Duplicated* (dark blue): entire contig maps to at least 5 locations in genome.

We expect that each contig should map to a unique location in the reference, and we find that to be the case for 4,540 contigs (38% of contigs, 45% of genome size). We find that an additional 4,827 contigs (41% of contigs, 49% of genome size) map primarily to a single contig but not a single location (Figures S1.6-S1.8). These cases could indicate that the w2rap contigs contain repetitive elements, are chimeric in content, or were able to bridge a gap in the reference assembly. We developed a method to identify regions that were likely in the latter case and successfully used the w2rap assembly to patch gaps in the Hmel2 reference (Supplement Section 2). A similar successful approach had also been adopted using our first pass *DISCOVAR de novo H. erato* assembly by the team assembling the *H. erato demophoon* reference genome [4].

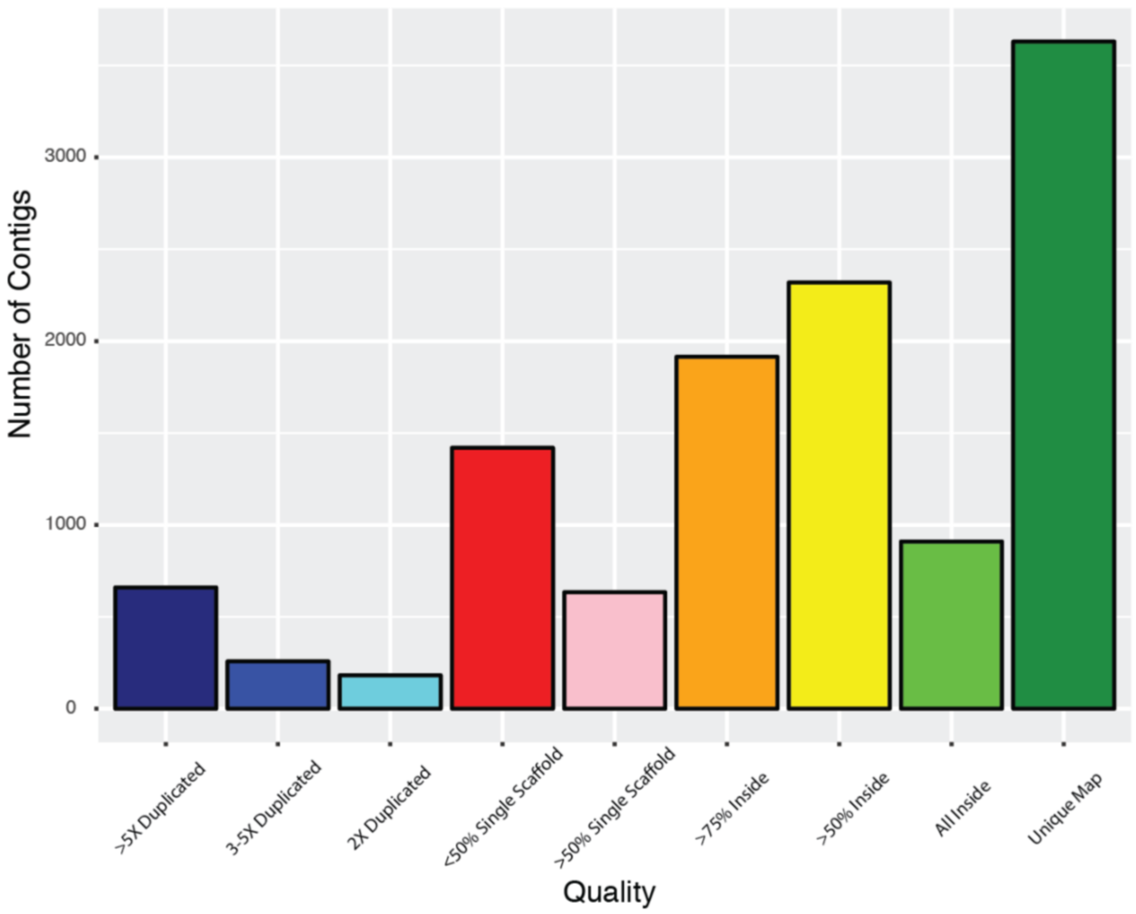

**Figure S1.6: *H. melpomene* w2rap to Hmel2 contig alignment quality**

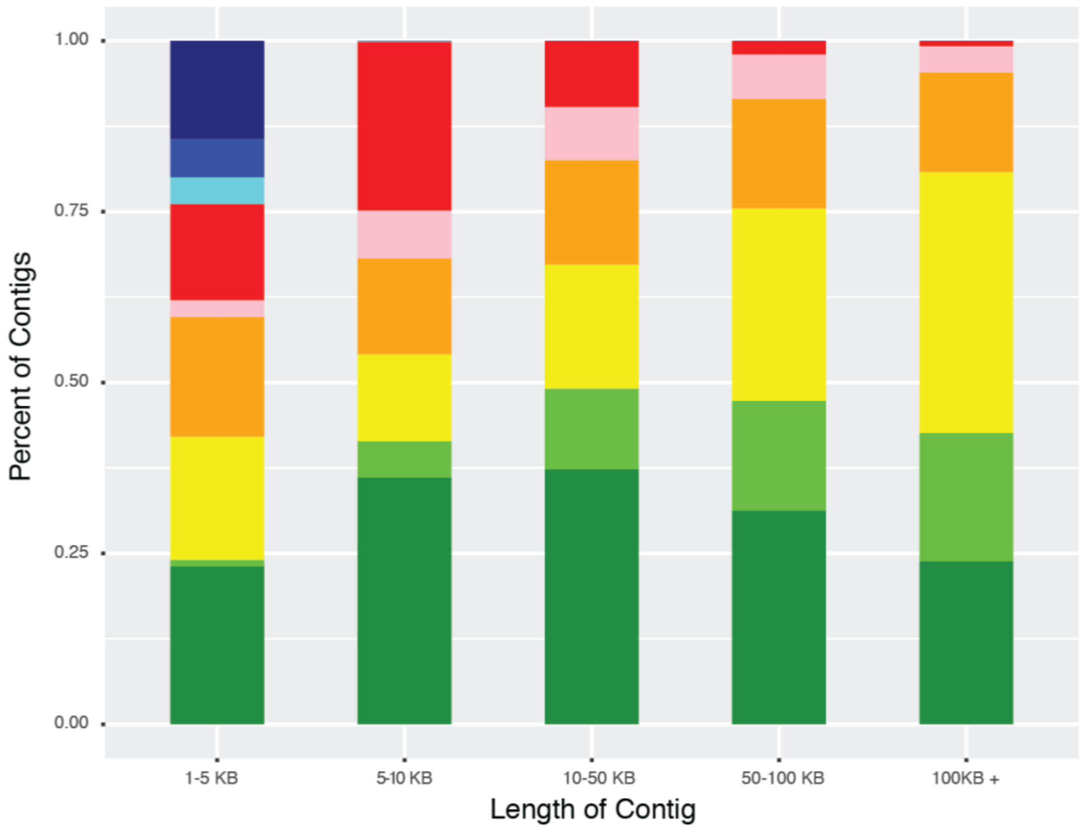

**Figure S1.7: Contig alignment quality by contig size**

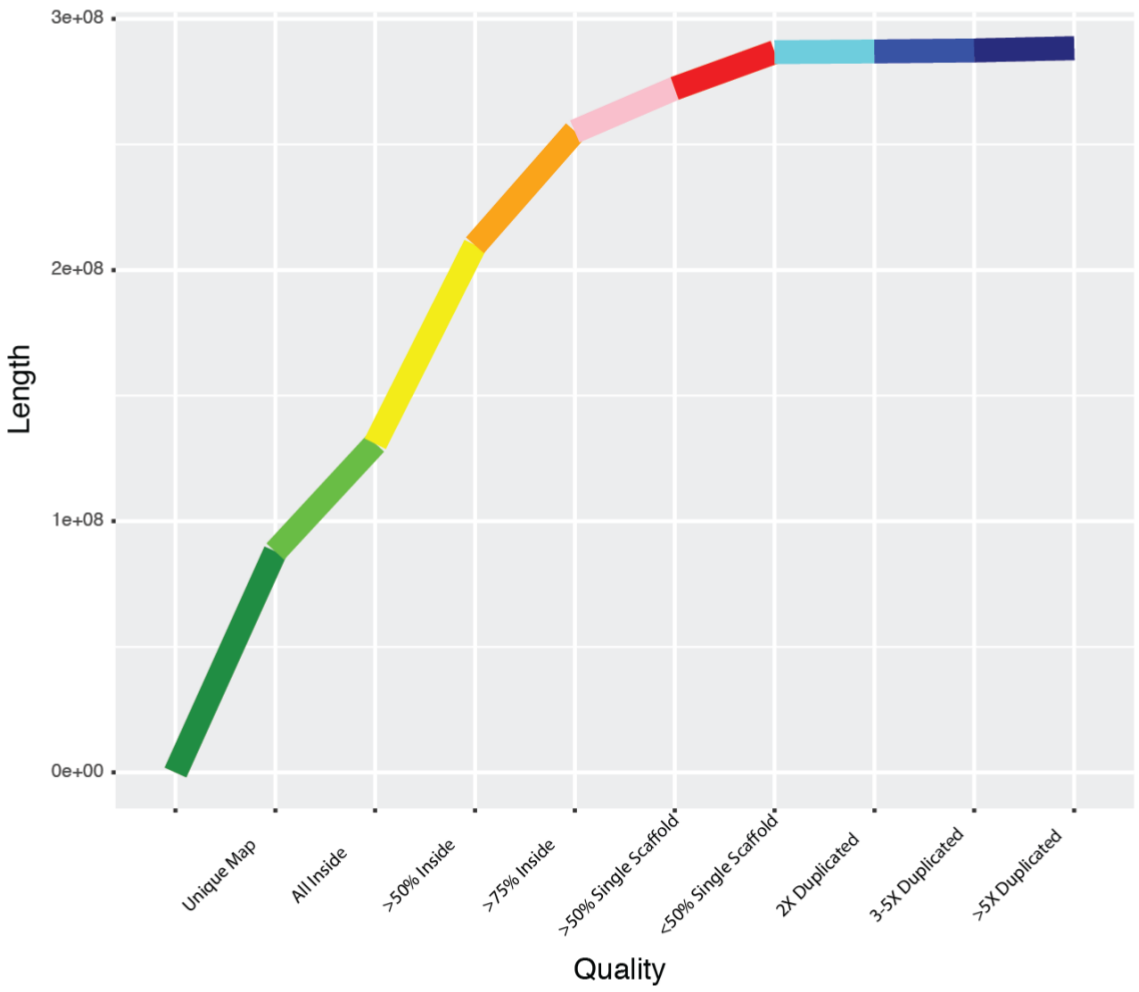

**Figure S1.8: Cumulative alignment length by quality**

**Section 2: Scaffolding With *DISCOVAR***

*return to top*

The assemblies in this study were generated with high coverage, PCR-free sequencing. One advantage of this strategy is that slightly imperfect repeats can be distinguished more readily than in traditional assemblies that must account for PCR errors and base call uncertainty. Generally, contigs tend to “break” when they encounter repetitive elements, and if w2rap is more efficient at assembling these areas, we should be able to link contigs within scaffolds with the *de novo* assemblies. In fact, we may be able to correct ordering and orientation errors as well, but here we focus minimally on gap-filling between contigs. We developed a pipeline "Scaffolding With *DISCOVAR*" (SWD) for this purpose, though it can be used with any pair of assemblies and is not limited to *DISCOVAR/w2rap*. Code is available at www.github.com/nbedelman/ScaffoldingWithDiscovar . SWD was designed with the assumption that one assembly has high-confidence scaffolding but some gaps (“reference genome”), and the other has high-confidence base-call quality but is not well scaffolded (“filler genome”). The flow chart for this process is shown in Figure S2.1 and is composed of 4 major steps. Below we give an explanation of the process assuming one is attempting to fill gaps between contigs within scaffolds. The tool will also be useful for ordering and orienting scaffolds, but these functionalities are still under development as of this writing. The commands below can be altered to fill gaps between scaffolds within chromosomes by simply replacing “chromosome” for “scaffold” and “scaffold” for “contig”.

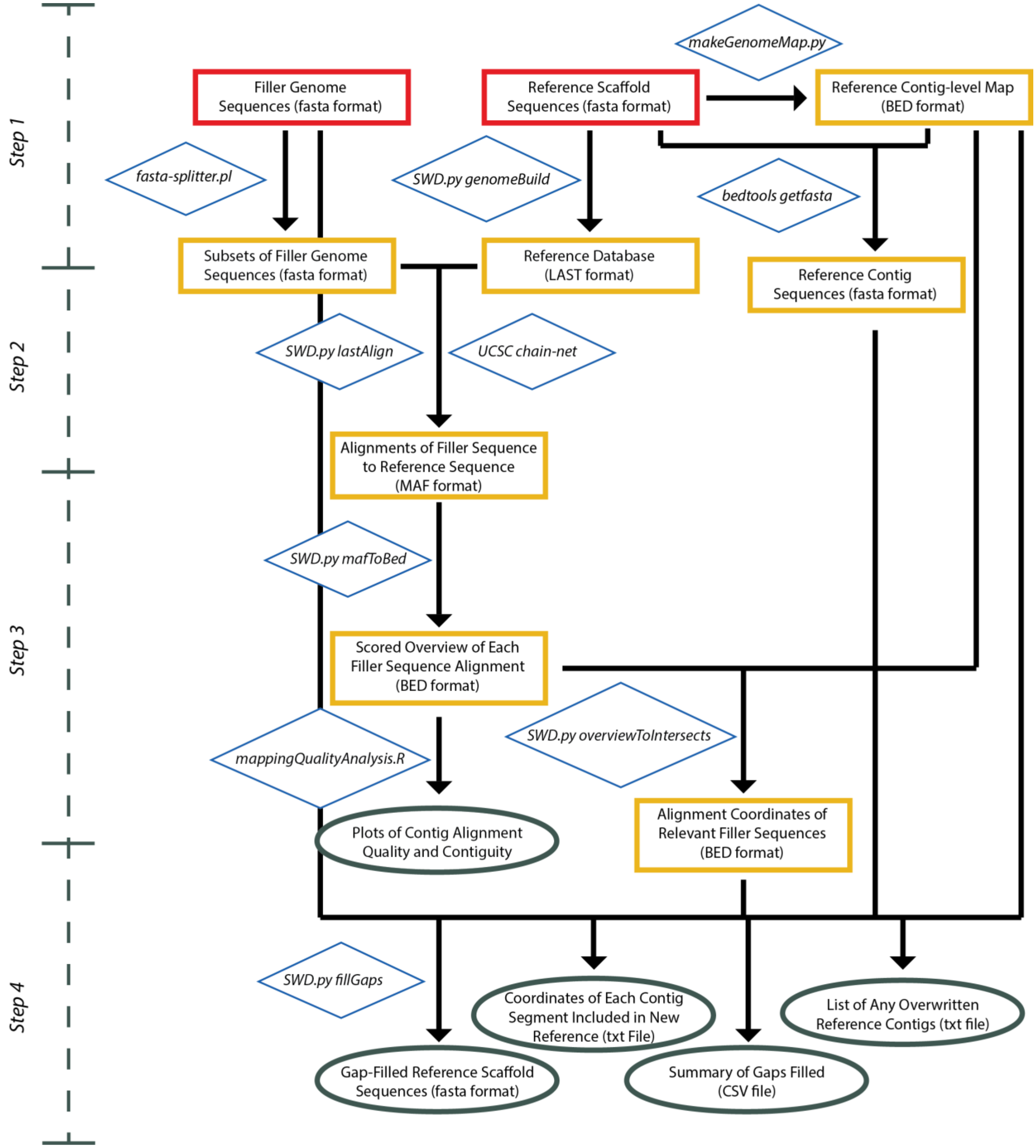

**Figure S2.1: Scaffolding with *DISCOVAR* flow chart**

*Required input files are outlined with red boxes, intermediate data files are outlined with gold boxes, and final outputs are outlined with green boxes. Blue diamonds adjacent to arrowed lines hold scripts used to convert the file(s) at the base of the arrow to the file(s) at the tips. Many of these rely on previously published tools (fasta-splitter.pl was written by K. Kryukov; many scripts rely on bedtools). Further details about each step are given in the text below.*

*Step 1: Preparing the data*
SWD takes:
1) a reference genome fasta with a sequence for every scaffold and containing a string of N's between each contig,
2) a reference genome fasta with a sequence for every contig,
3) a map in bed format indicating the location of each contig on its respective scaffold, and
4) a filler genome.

If a user has (1), the scripts "*makeGenomeMap.py*" (SWD script) and *bedtools getfasta* (from the bedtools suite) will convert such reference genome fastas into (2) and (3). *makeGenomeMap.py* is capable of dividing a genome either at regions of *at least* a given number of N's (default) or a specific number of N's (flag "exact").

*Step 2: Aligning the filler genome to the reference genome*
This pipeline was developed using LAST, but any aligner can be used so long as the output is in MAF format[6]. The SWD package has wrapper scripts for building a genome index and running the LAST aligner *(SWD.py genomeBuild* and *SWD.py lastAlign*), which are optional but hopefully useful. After the genome has been aligned, the pipeline follows the chain and net protocol developed by UCSC, which can be read in more detail here ([7]). This ensures that we are examining the location of best alignment for each filler genome contig on the reference genome.

*Step 3: Filtering for contigs that may be useful in gap-filling*
We convert the resulting MAF alignments into specially-formatted BED files. Specifically, each BED file is comprised of the alignment location of a single filler genome contig and contains information as to which part of the contig aligns to which reference region, and provides a score based on how contiguous the full contig alignment is. The scoring strategy begins by projecting the location of the end points of the contig out from the longest sub-alignment. For example, if a *DISCOVAR* contig is 10KB in length, and its longest sub-alignment begins at contig position 2000 and ends at contig position 5000, and maps to a reference scaffold from positions 102000 to 105000, the projected position would be from reference scaffold 100000 to 110000(Figures S2.2, S2.3). The contig would then be scored as follows:

10: The contig maps as a single alignment to a single location in the reference genome
9: The contig maps as multiple sub-alignments, but at least 95% maps within the projected region
8: At least 75% of the contig maps within the projected region
7: At least 50% of the contig maps within the projected region
6: At least 50% of the contig maps to the same scaffold as the longest sub-alignment
5: Less than 50% of the contig maps to the same scaffold as the longest sub-alignment
4: The entirety of the contig maps to exactly two locations in the reference genome
3: The entirety of the contig maps to 3-5 locations in the reference genome
2: The entirety of the contig maps to more than 5 locations in the reference genome

Contigs whose score ranges from 6 to 8 are retained for the next step.

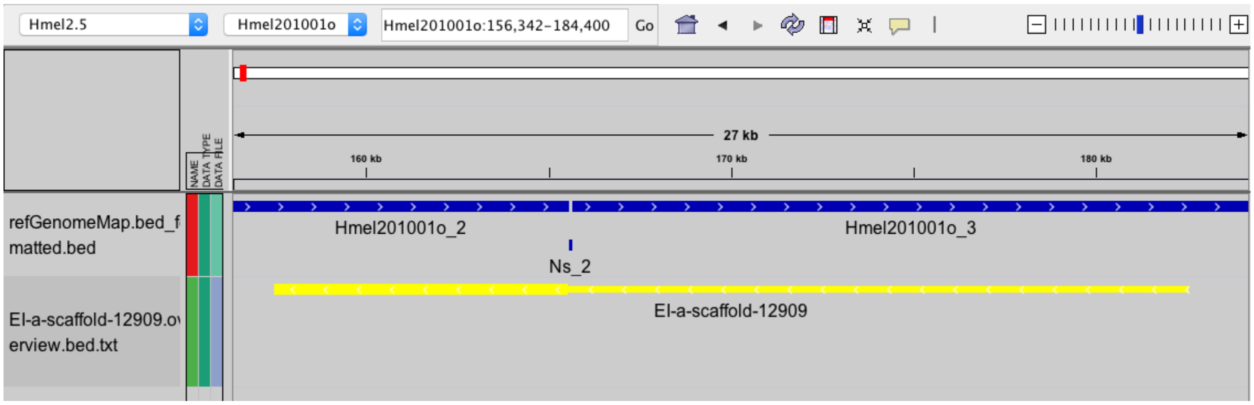

**Figure S2.2: Example overview alignment**
*This screenshot from IGV shows a 27KB region of Hmel2.5 chromosome 1. The blue track displays the location of reference contigs Hmel201001o_2 and Hmel201001o_3, as well as the stretch of Ns that separates them. The yellow track shows the longest sub-alignment of the de novo H. melpomene genome contig EI-a-scaffold-12909, which overlaps with Hmel201001o_2 (thick line). When we project the remainder of EI-a-scaffold-12909 based on the location of the sub-alignment (thin line), we see that it is expected to continue on to Hmel201001o_3. This overview is colored yellow, indicating that more than 75% of the sub-alignments of this contig are found within the projected area, as can be seen in Figure S2.3.*

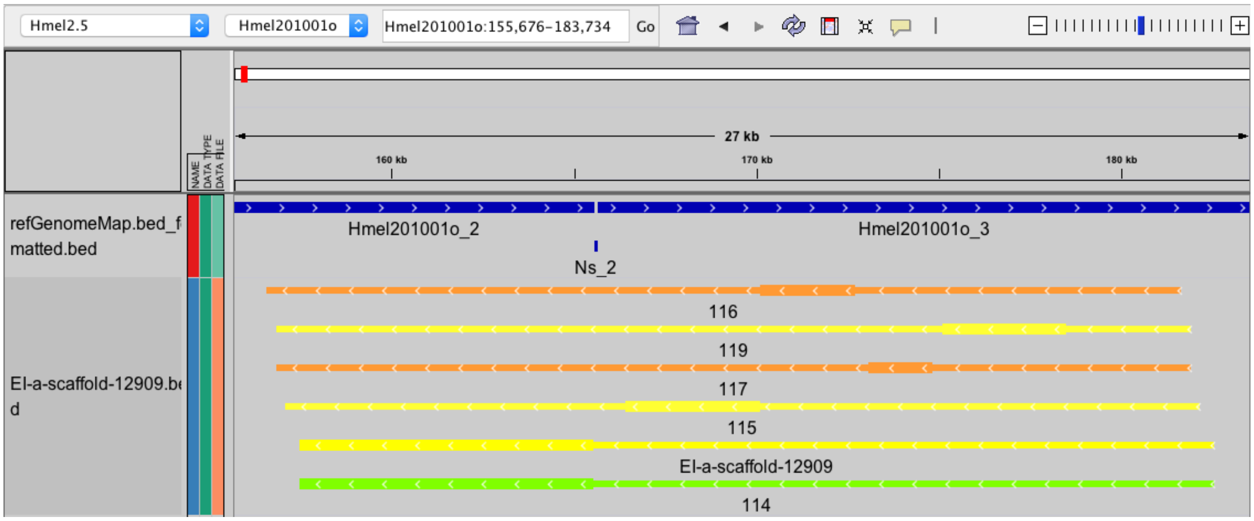

**Figure S2.3: Example full contig alignment**
*Here, the remaining sub-alignments of EI-a-scaffold-12909 are displayed in the same manner as the overview – the thick lines indicate the location of the sub-alignment, while the thin lines project the position of the contig based on the aligned region. One can see that the sub-alignments are located in positions expected if Hmel201001o_2 and Hmel201001o_3 are in fact contiguous. The alignments labeled 114-119 are the sub-alignments, and their colors are simply an indication of size relative to the full contig.*

*Step 4: Gap Filling*
This step takes as input the bed files of contigs that passed the filter in step 3, the fasta file of reference genome contigs, and the fasta file of the filler genome contigs. For gap filling, as was done here, the flag "consecutiveOnly" must be set to True. The procedure in this step is as follows:

First, we identify groups of reference contigs and filler contigs that are linked together. For example, if filler contig 1 overlaps with reference contigs A and B, and filler contig 2 overlaps with reference contigs B and C, reference contigs A, B, and C, and filler contigs 1 and 2 would form a group. Next, we consolidate sub-alignments that are collinear and neighbor one another on a single reference contig. Once the sub-alignments are consolidated, we form super-contigs by joining together all the reference contigs that can be linked by each filler contig. In order to approve a link, there must be at least 1000bp of aligned sites on each of the reference scaffolds, and the order and orientation must be correct. This means that for reference contigs A and B to be linked by filler contig 1, the beginning of filler contig 1 must overlap the end of reference contig A by at least 1000bp, and the end of filler contig 1 must overlap the beginning of reference contig B by at least 1000bp. Once the super-contigs are made with the information from each individual fill contig, the super-scaffolds within a group are linked together in a similar manner. In each case, the end of the reference contig being joined will be replaced by the filler contig that overlaps it. This ensures contiguity across the gap and reflects the heuristic that the per-base accuracy of the filler contig is more trustworthy in these areas than the reference contig (Figures S2.3-S2.4, Table S2.1).

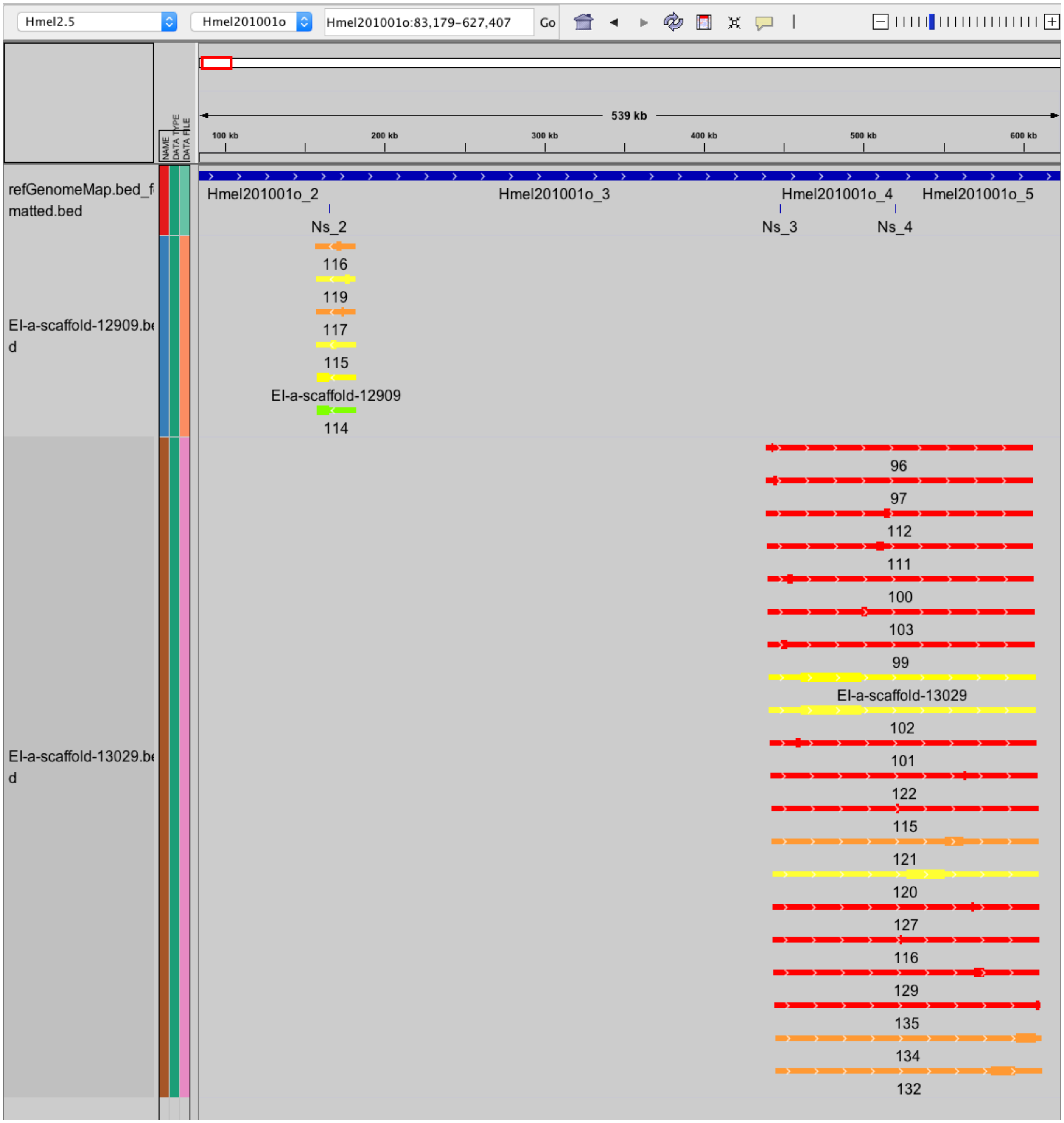

**Figure S2.4: Example full group alignment**

*Here, we see that EI-a-scaffold-13029 maps to reference contig Hmel201001o_3 as well, and also connects to Hmel201001o_4 and Hmel201001o_5. These regions are treated as a group, and are joined into a single super-scaffold, keeping as much of the reference contigs as possible but joining them together with the filler contigs (See Table S2.1). Three regions of the de novo filler genome (names beginning with EI) were used to connect the four reference contigs (names beginning with Hmel) in the group. Each of these regions is between 500 and 1300 base pairs, and replaces fewer than 1000 bp of any individual reference contig.*

In this study, we filled gaps solely within reference scaffolds. Taking a conservative approach, in which we assumed that the published order of contigs was correct, we were able to fill 30% of gaps (899/2990 stretches of >= 100 Ns) and increased the contig N50 by 39%. The overall genome size was reduced by 97KB, after a reduction of 248 KB of Ns (Table S2.2). A similar approach was previously employed to use DISCOVAR data to fill gaps in the *H. erato demophoon* v1 reference genome[4], resulting in a 12X improvement in N50. We examined the regions of the w2rap contigs that were inserted into the reference genome and found that they were more than twice as rich in repetitive elements as the full genome. This elevation was driven primarily by retroelements (15.69% of inserted sequences, 1.52% of whole genome) and small RNAs (9.41% of inserted sequences, 0.02% of whole genome). Interestingly, the abundance of DNA transposons decreased in inserted regions, comprising just 4.72% as opposed to 10% in the whole genome. In order to confirm that our scaffolding procedure was not collapsing tandem repeats into single elements, we calculated read coverage in 10KB windows and compared the mean coverage of gap-filled regions to that of other genomic loci. We found that the distribution of coverage depth was the same between the two groups, consistent with valid contig joins (Figures S2.5-S2.7).

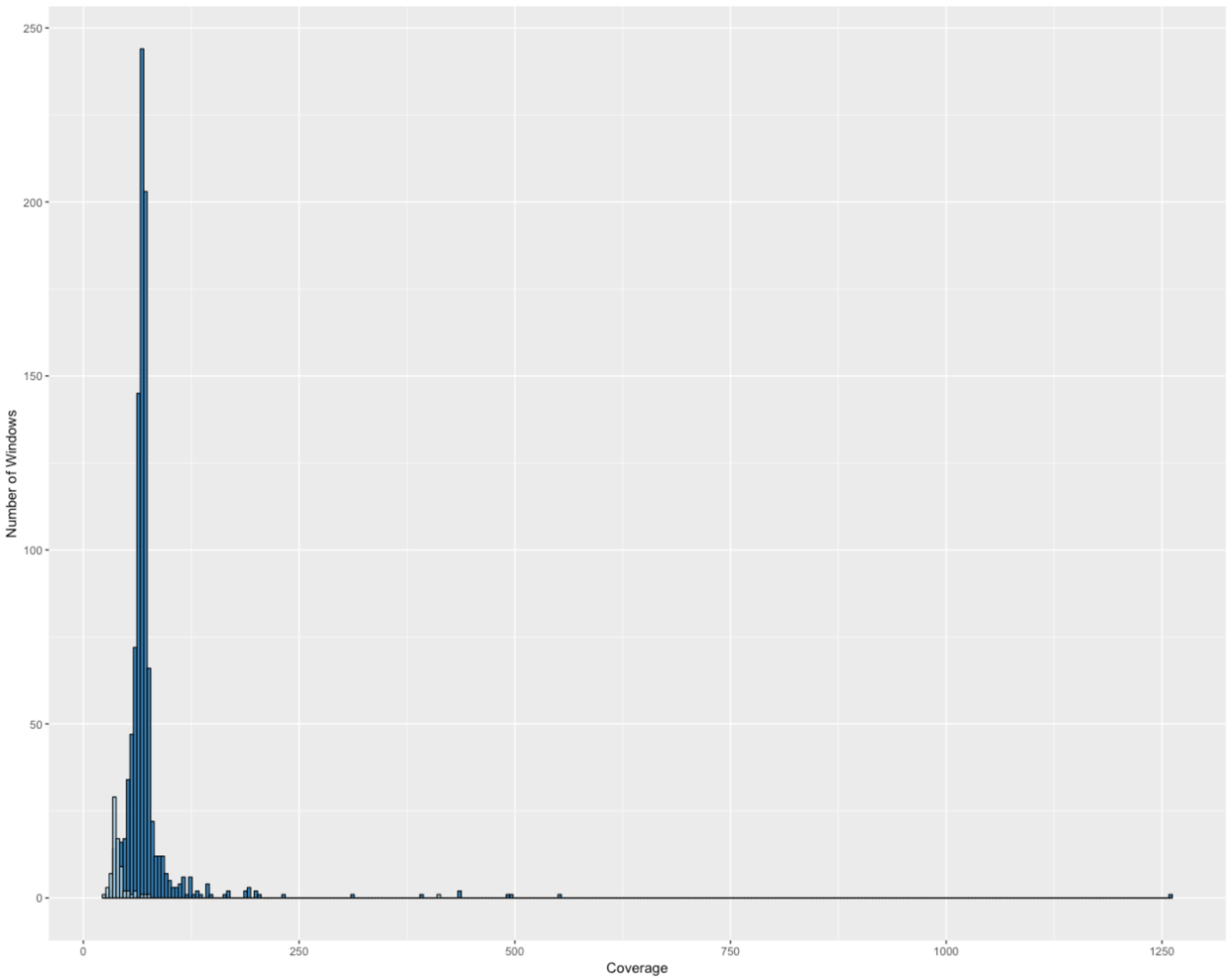

**Figure S2.5: Coverage of gap-filled regions**

*Histogram of mean coverage in 10KB windows overlapping gap-filled regions. Light blue is the Z-chromosome, and dark blue is autosomes.*

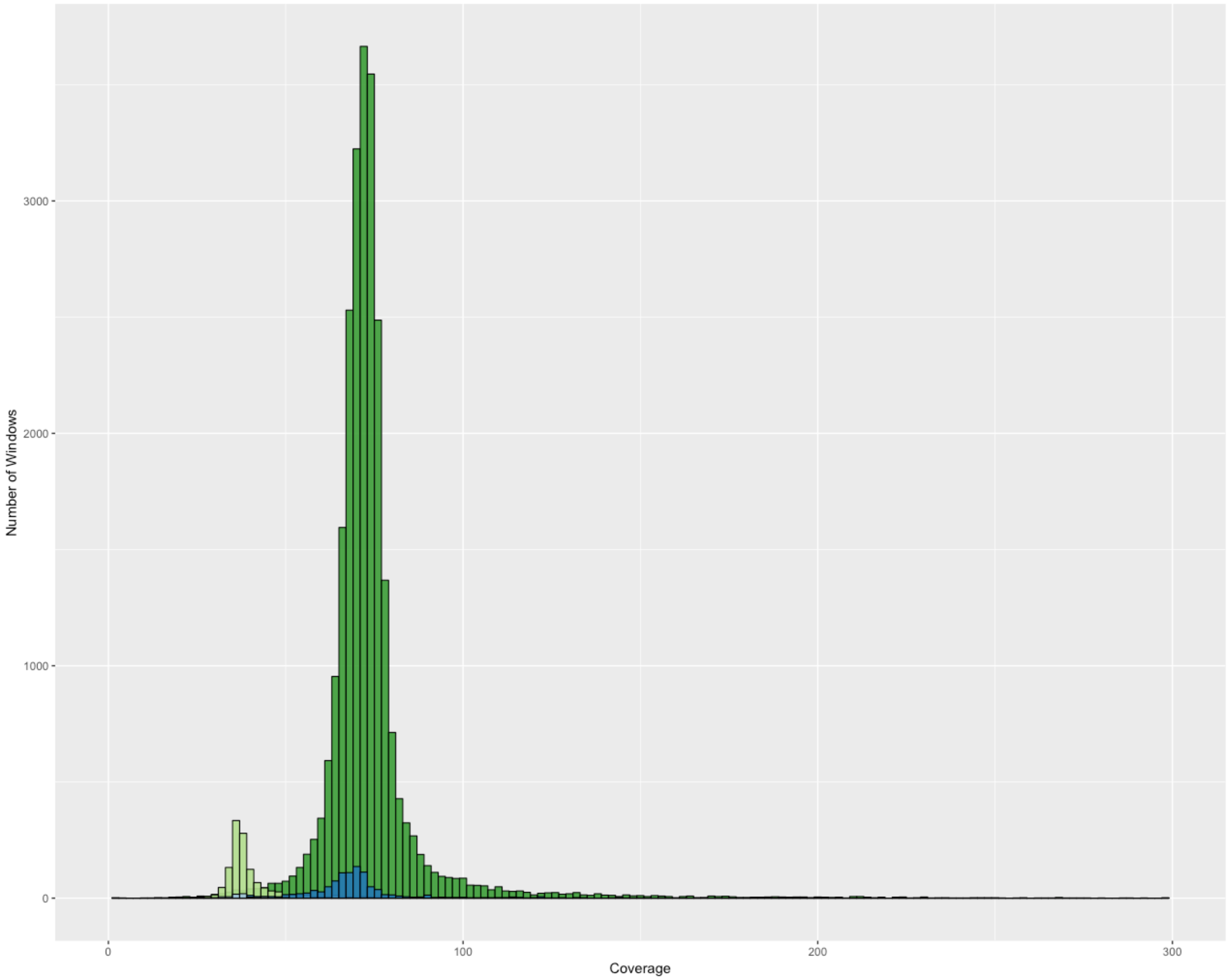

**Figure S2.6: Coverage of gap-filled and non gap-filled regions**
*Histogram of mean coverage in 10KB windows. Blues are the same data as Figure 2.5, light green is non gap-filled Z chromosome, and dark green is non gap-filled autosomes*

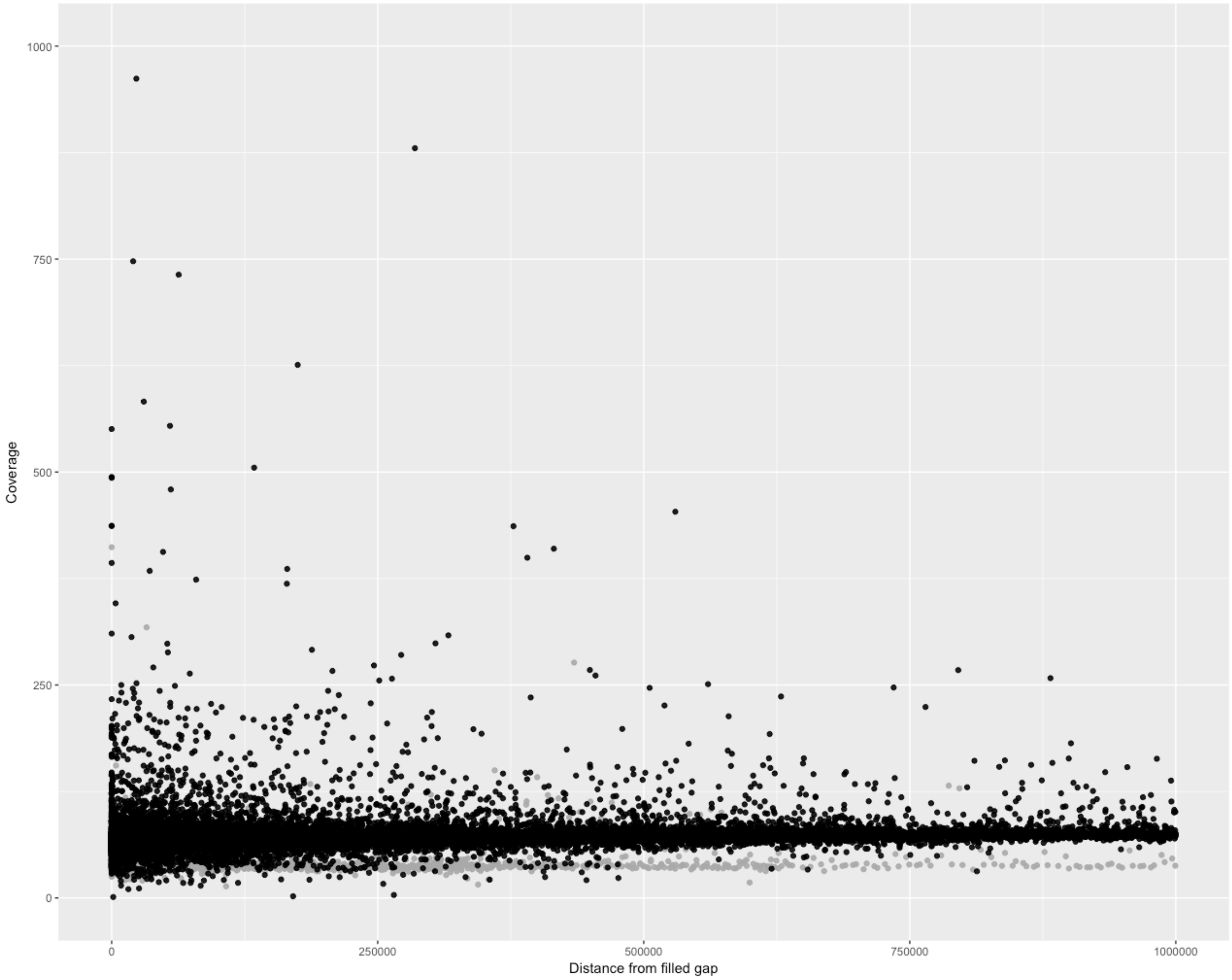

**Figure S2.7: Coverage with distance from filled gap**
*Histogram of mean coverage in 10KB windows with distance from the nearest filled gap on the x-axis. Gaps tend to be in repetitive regions, as seen by increased coverage near zero, but the increase in coverage begins well before the filled gap itself.*

**w2rap melpomene full genome**

```
==================================================
sequences:         12179
total length:  289529429 bp  (289477054 bp excl N/X-runs)
GC level:         32.74 %
bases masked:   42390139 bp ( 14.64 %)
==================================================
               number of      length   percentage
               elements*    occupied  of sequence
--------------------------------------------------
Retroelements        18032      4390549 bp    1.52 %
   SINEs:              684        34049 bp    0.01 %
   Penelope              2          155 bp    0.00 %
   LINEs:            12167      3257088 bp    1.12 %
    CRE/SLACS            0            0 bp    0.00 %
     L2/CR1/Rex       6128      1511859 bp    0.52 %
     R1/LOA/Jockey    2340       674824 bp    0.23 %
     R2/R4/NeSL       1459       468411 bp    0.16 %
     RTE/Bov-B        1740       483943 bp    0.17 %
     L1/CIN4            12          624 bp    0.00 %
   LTR elements:      5181      1099412 bp    0.38 %
     BEL/Pao           633       144481 bp    0.05 %
     Ty1/Copia         178        40374 bp    0.01 %
     Gypsy/DIRS1      3862       807954 bp    0.28 %
       Retroviral        0            0 bp    0.00 %

DNA transposons     231972     30820377 bp   10.64 %
   hobo-Activator     8196      1506936 bp    0.52 %
   Tc1-IS630-Pogo   123631     15828698 bp    5.47 %
   En-Spm                0            0 bp    0.00 %
   MuDR-IS905            0            0 bp    0.00 %
   PiggyBac             88        35700 bp    0.01 %
   Tourist/Harbinger    11         1563 bp    0.00 %
   Other (Mirage,      146         7447 bp    0.00 %
    P-element, Transib)

Rolling-circles          0            0 bp    0.00 %

Unclassified:         1997       282672 bp    0.10 %

Total interspersed repeats:    35493598 bp   12.26 %

Small RNA:             790        49738 bp    0.02 %

Satellites:            331        18247 bp    0.01 %
Simple repeats:     128725      5687070 bp    1.96 %
Low complexity:      25791      1246843 bp    0.43 %
```

**w2rap sequences inserted across gaps**

```
==================================================
sequences:           792
total length:    1909413 bp  (1908113 bp excl N/X-runs)
GC level:          32.90 %
bases masked:     600714 bp ( 31.46 %)
==================================================
               number of      length   percentage
               elements*    occupied  of sequence
--------------------------------------------------
Retroelements         1805       299637 bp   15.69 %
   SINEs:             1432       179734 bp    9.41 %
   Penelope             15         7175 bp    0.38 %
   LINEs:              274        91167 bp    4.77 %
    CRE/SLACS            0            0 bp    0.00 %
     L2/CR1/Rex         84        29195 bp    1.53 %
     R1/LOA/Jockey      23         5461 bp    0.29 %
     R2/R4/NeSL         19         7326 bp    0.38 %
     RTE/Bov-B          71        24710 bp    1.29 %
     L1/CIN4             0            0 bp    0.00 %
   LTR elements:        99        28736 bp    1.50 %
     BEL/Pao             9         2630 bp    0.14 %
     Ty1/Copia           5          322 bp    0.02 %
     Gypsy/DIRS1        45        10596 bp    0.55 %
       Retroviral        0            0 bp    0.00 %

DNA transposons        620        90142 bp    4.72 %
   hobo-Activator       41         8030 bp    0.42 %
   Tc1-IS630-Pogo      371        40697 bp    2.13 %
   En-Spm                0            0 bp    0.00 %
   MuDR-IS905            0            0 bp    0.00 %
   PiggyBac             51        10913 bp    0.57 %
   Tourist/Harbinger     0            0 bp    0.00 %
   Other (Mirage,        9          704 bp    0.04 %
    P-element, Transib)

Rolling-circles          0            0 bp    0.00 %

Unclassified:          652       174562 bp    9.14 %

Total interspersed repeats:      564341 bp   29.56 %

Small RNA:            1431       179655 bp    9.41 %

Satellites:              7          338 bp    0.02 %
Simple repeats:        695        30687 bp    1.61 %
Low complexity:        137         6194 bp    0.32 %
```

**Table S2.3: Comparison of general *H melpomene de novo* repeat content with repeat content of regions used to fill gaps in Hmel2.**

**Section 3: progressiveCactus Alignment**
*return to top*

We generated a multi-species whole genome alignment using progressiveCactus[8,9]. In order to create a high-quality alignment with power to detect evolutionary change at the root of the *Heliconius* clade, we filtered *de novo Heliconius* assemblies for those with fewer than 50,000 scaffolds, and further filtered the resulting assemblies for scaffolds larger than 1000 base pairs. We included previously published Lepidoptera assemblies including three additional species in the family *Nymphalidae* (*Melitaea cinxia* v1.0[10], *Bicyclus anynana* v1.2[11] and *Danaus plexippus* v3[12]), one additional butterfly (*Papilio polytes* v1.0[13]), one skipper (*Lerema accius* v1.1[14]), and two moths (*Bombyx mori* ASM15162v1[15] and *Plutella xylostella* DBM_GJ_v1.1[16]). Genome size filtering was performed with a custom python script.

In order to run progressiveCactus on the Harvard Odyssey compute cluster, modification of the program was carried out by Harvard Research Computing to integrate with the SLURM scheduler. The modified version we used can be found here: https://github.com/harvardinformatics/progressiveCactus .

The final alignment was run with the command

```
runProgressiveCactus.sh <sequence file> <work directory> <hal output file>
--batchSystem=slurm \
--slurm-partition=serial_requeue \
--slurm-jobname <job name> \
--slurm-scriptpath=<batch script path> \
--slurm-time=1440 --slurm-constraint='holyib' \
--defaultMemory=6000000000 --bigMemoryThreshold=5000000000 \
--retryCount=3 --bigBatchSystem=singleMachine \
--configFile=local_config.xml \
--maxThreads=32 \
--logLevel=DEBUG \
--stats
```

All config files are available on Dryad.

In order to assess the success of the progressiveCactus alignment, we calculated pairwise alignment coverage between all species and *H. melpomene* Hmel2.5[17] (Figure S3.1, Table S3.1). We also computed cumulative 'alignment depth' using *H. melpomene* and *B. mori* as reference sequences. The cumulative alignment depth is a measure, calculated for each number between 0 and the total species in the alignment, of the percent of the reference genome that is aligned among at least that number of species (Figures S3.2,S3.3).

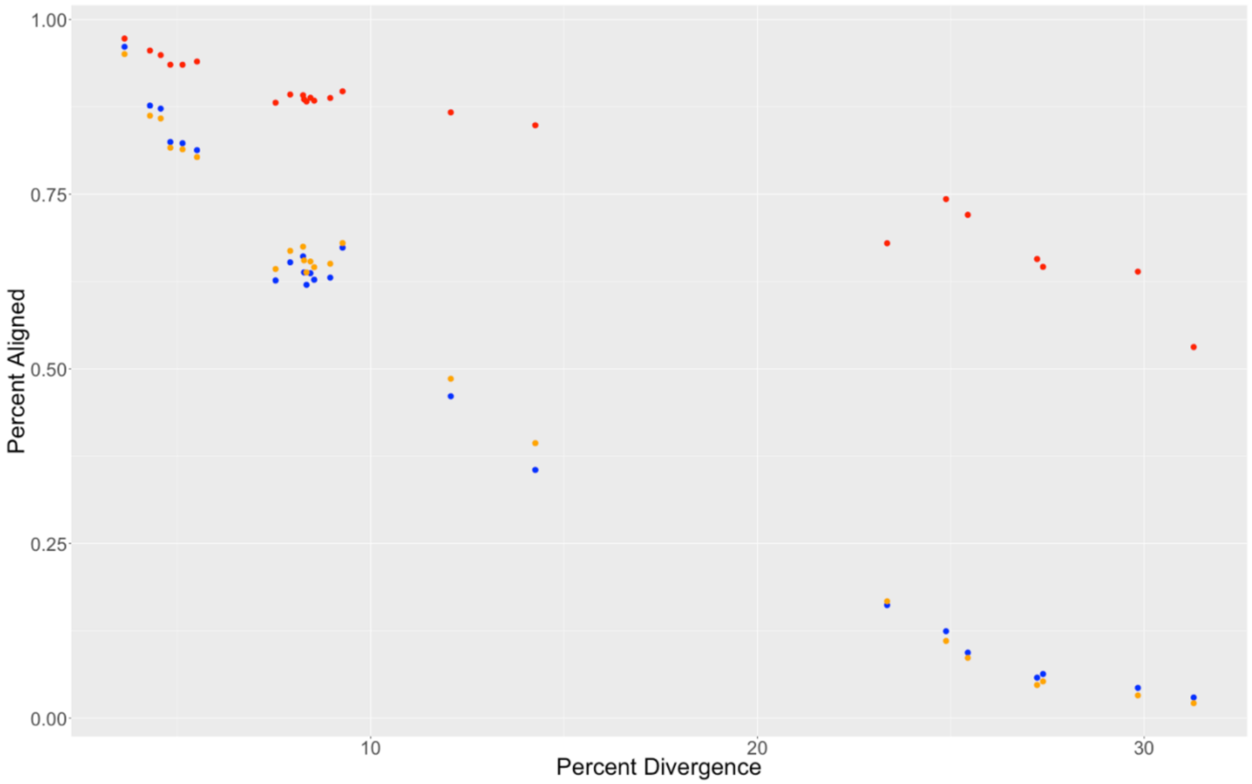

**Figure S3.1: Pairwise alignment depth**

*Figure S3.1 and Table S3.1 show the pairwise alignment depth between other Lepidoptera and Hmel2.5 in the progressiveCactus alignment. The "percent divergence" metric is the pairwise distance in concatenated non-coding regions aligned among all species. Exons are shown in red, introns in orange, and intergenic regions in blue.*

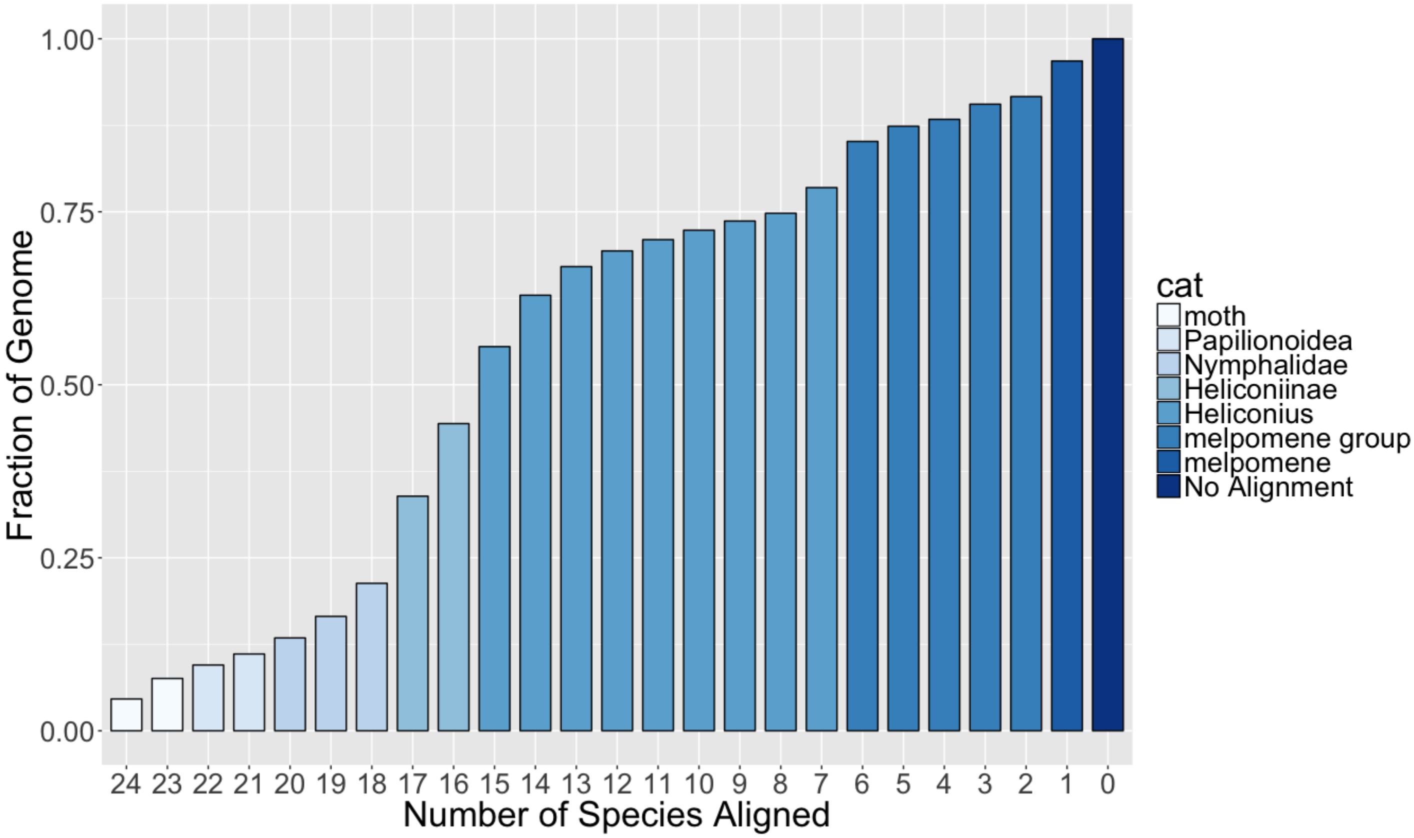

**Figure S3.2: Cumulative alignment depth to *H. melpomene* Hmel2.5**

*The Y-axis shows the percentage of Hmel2.5 sites that are aligned at least once among at least the number of species indicated on the X-axis. Colors represent the narrowest taxonomic category that corresponds to each number of species. For example, there are 15 Heliconius species not including Hmel2. Critically, however, the values of each bar correspond to number of species. The colors are therefore only a qualitative visual aid.*

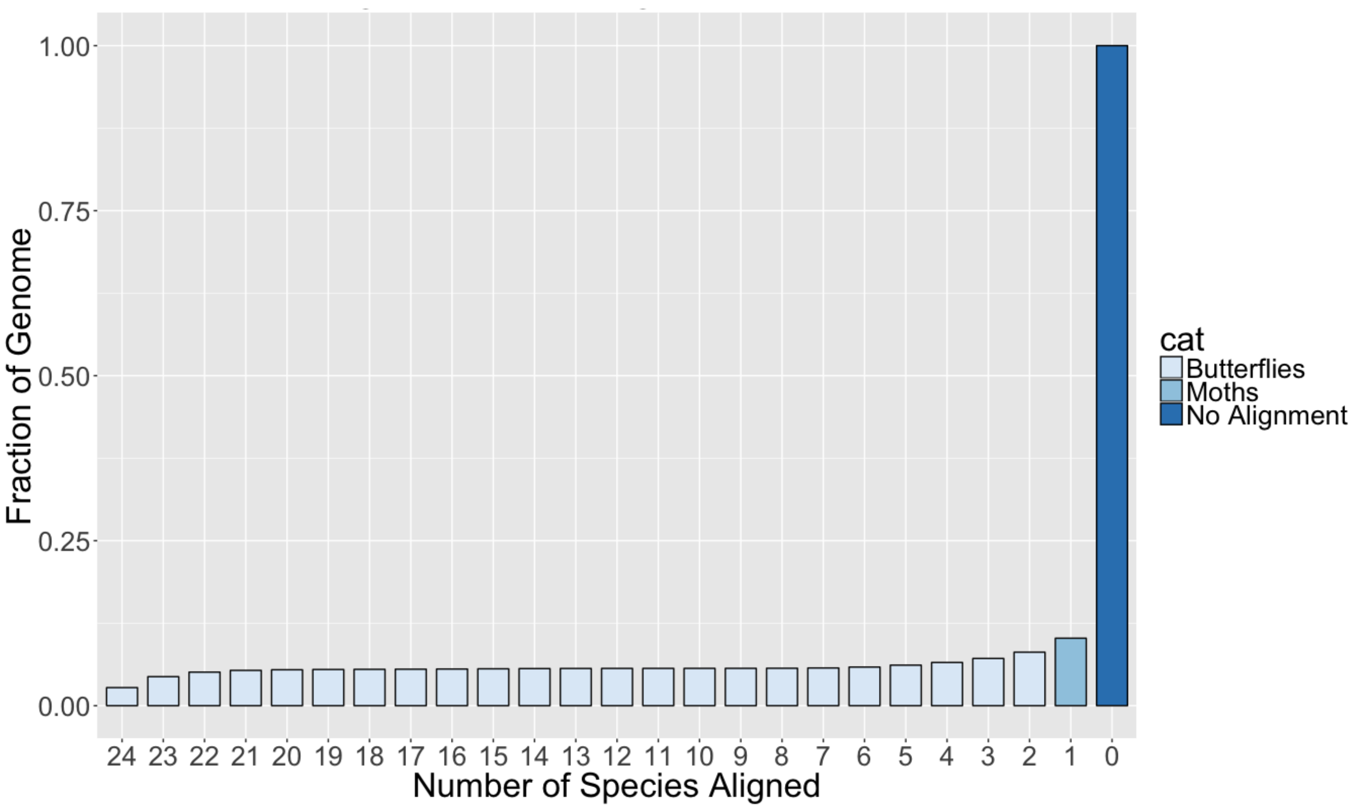

**Figure S3.3: Cumulative alignment depth to *B. mori***

*Same as figure S3.2, but using B. mori as the reference genome. Very little of the B. mori genome aligns to any other lepidopteran genome assemblies employed here.*

**Section 4: Phylogeny**
*return to top*

This data set and the resulting alignment resulted in a truly genome-wide dataset that we used to infer phylogenies on multiple subsets of the data. In general, we analyzed three different data sets: full genes, fully aligned sites (defined as sites that contain data for each species in the alignment) that are coding, and fully aligned sites that are non-coding. In addition, we generated the same data sets for only those taxa included in Heliconiini (*Heliconius* species plus *Agraulis* and *Eueides*) (Table S4.1). For the gene set, we removed all genes that had 3 or fewer taxa and greater than 30% missing data.

For each data set, we generated single locus trees, ASTRAL "species" trees (using the single locus trees as input), and concatenated trees. For each single locus tree, we used IQtree v. 1.6.1 to estimate the model (using the ModelFinder function in IQtree), the tree, and 1000 ultrafast bootstraps (command: iqtree -s <alignment> -m MFP -bb 1000 -safe) [18-20]. For each concatenated tree, we used IQtree v1.6.5 to:

(1) select an optimal partitioning scheme using the relaxed clustering algorithm (command: iqtree -s <alignment> -spp <partition_definition> -m TESTMERGEONLY -rclusterf 5 -rcluster-max 5000 -mset GTR -nt 48),

(2) select optimal models for each subset in the partitioning scheme selected from the previous run using ModelFinder, followed by a ML tree search and 1000 ultrafast bootstrap (command: iqtree -s <alignment> -spp <partition_definition>.best_scheme.nex -m MFP -nt 24 -safe -bb 1000), and

(3) conduct 15 additional ML searches for each concatenated data set using the relaxed clustering + ModelFinder partitioning scheme generated in step 2 (command for each search: iqtree -s <alignment> -spp <partition_definition>.best_scheme.nex -nt 24 -bb 1000 -safe). For each set of trees, we selected the tree with the best maximuim likelihood value.

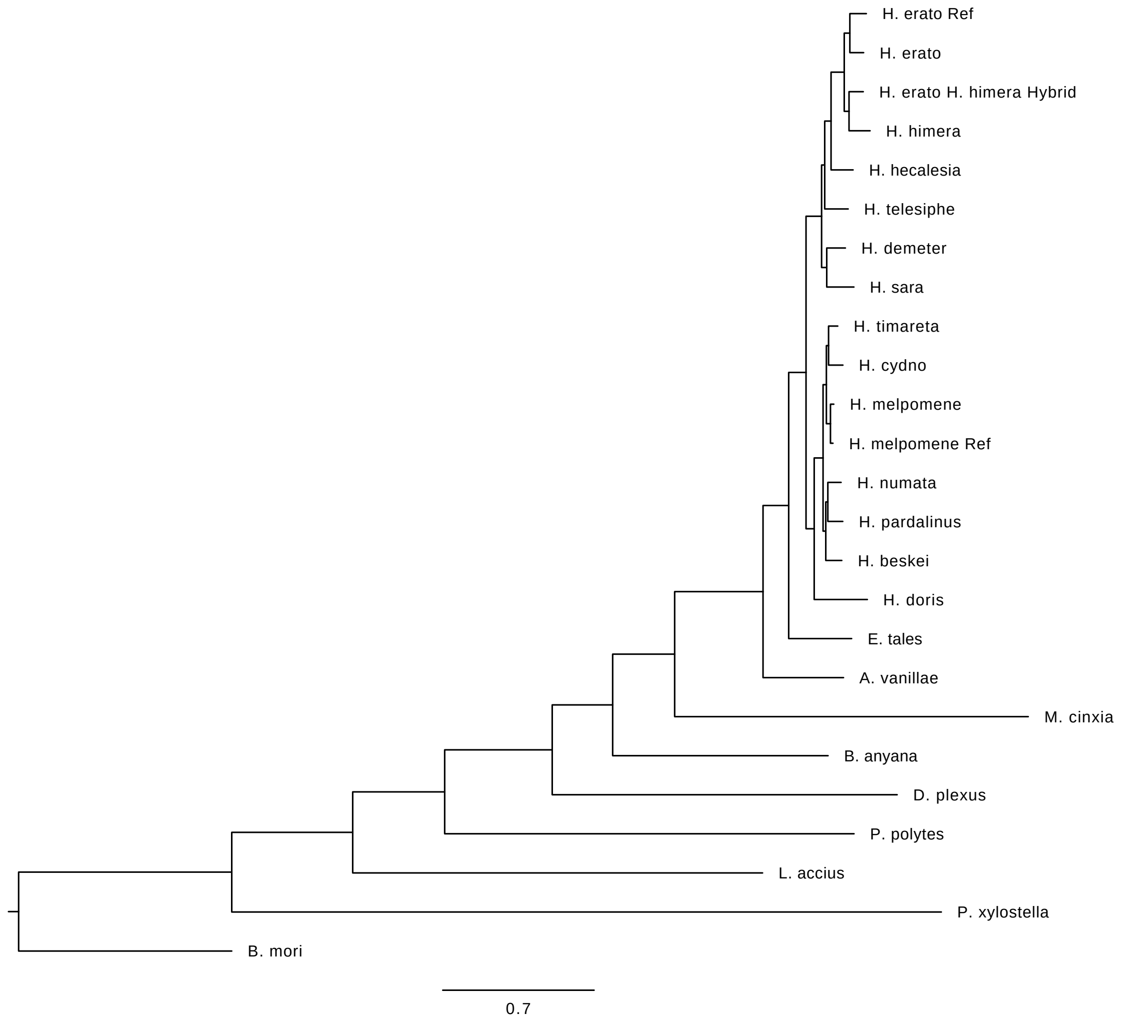

**Figure S4.1. Concatenated tree for genes.** All nodes had 100% bootstrap values.

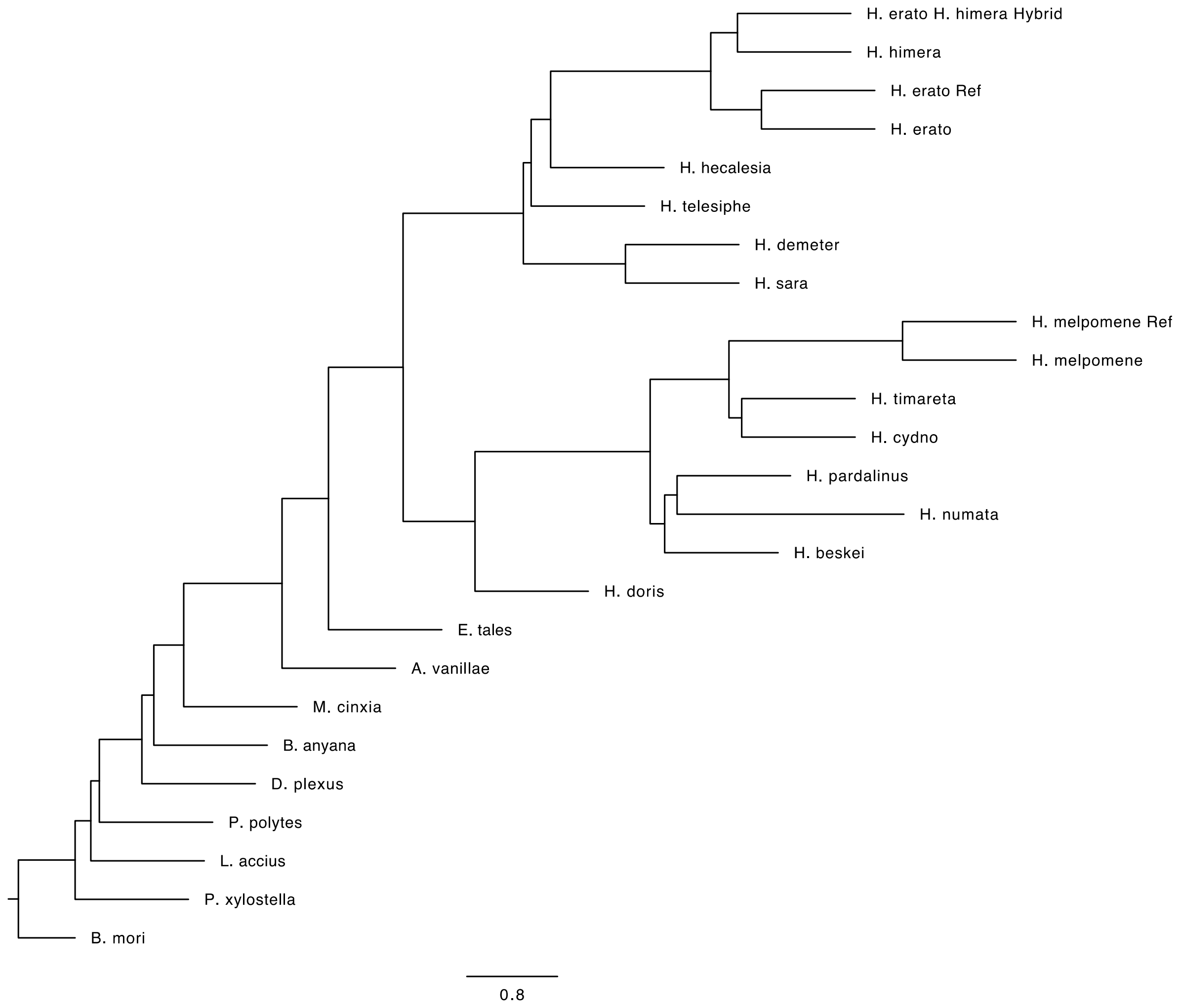

**Figure S4.2. ASTRAL tree for genes**

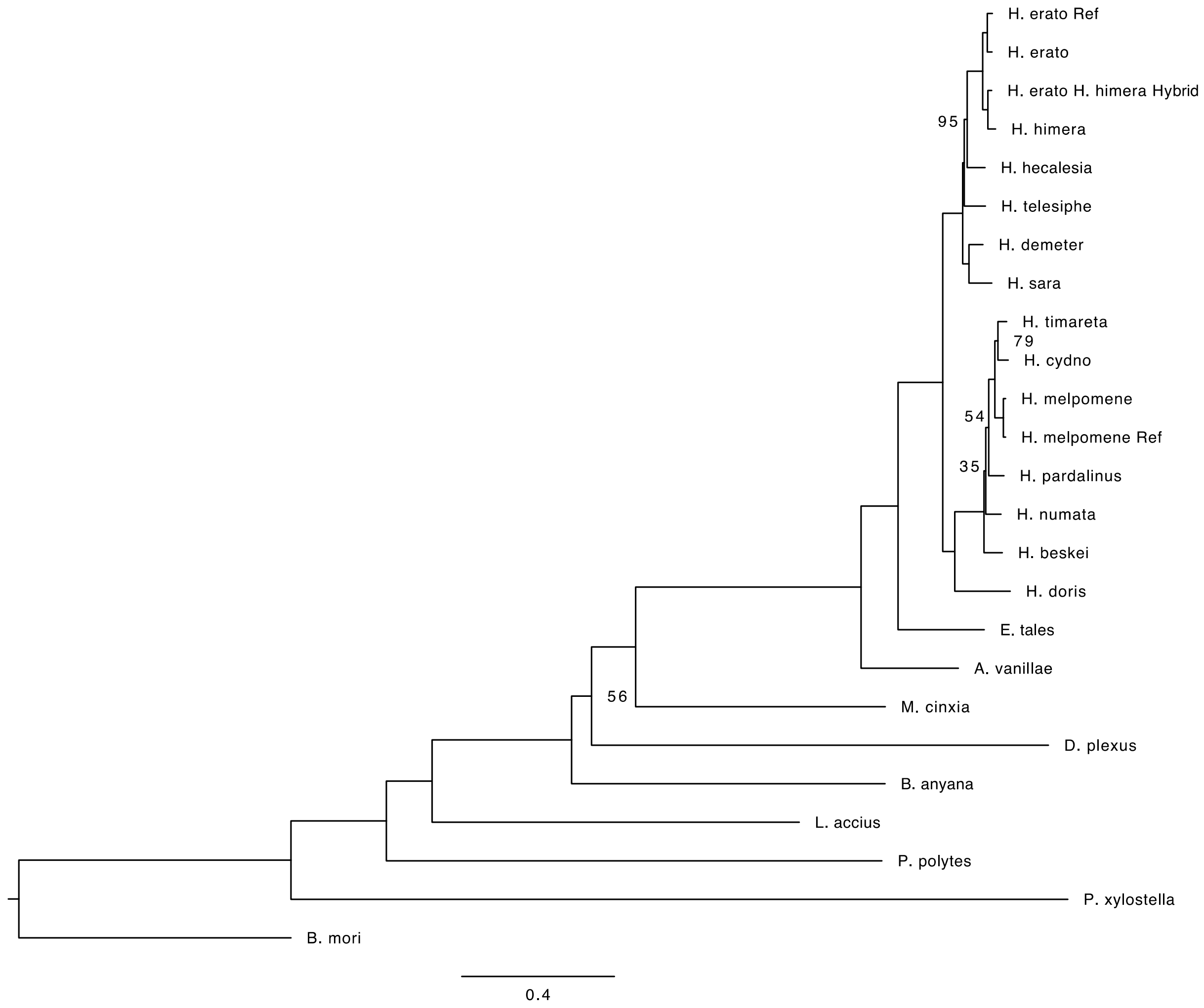

**Figure S4.3. Concatenated tree for noncoding, fully aligned blocks greater than or equal to 100 bp.** Bootstrap values are shown for nodes with less than 100% support.

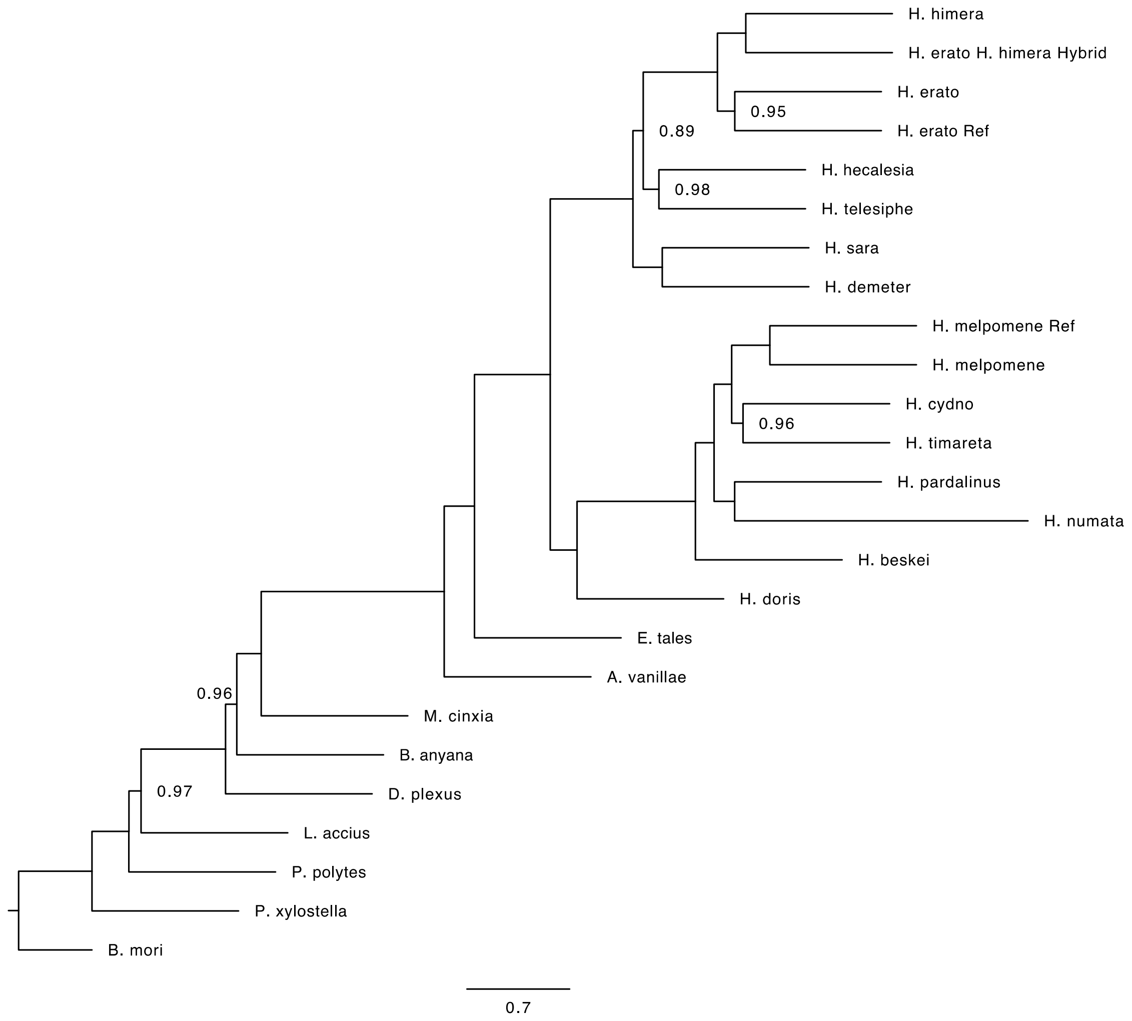

**Figure S4.4. ASTRAL tree for noncoding, fully aligned blocks greater than or equal to 100 bp.**

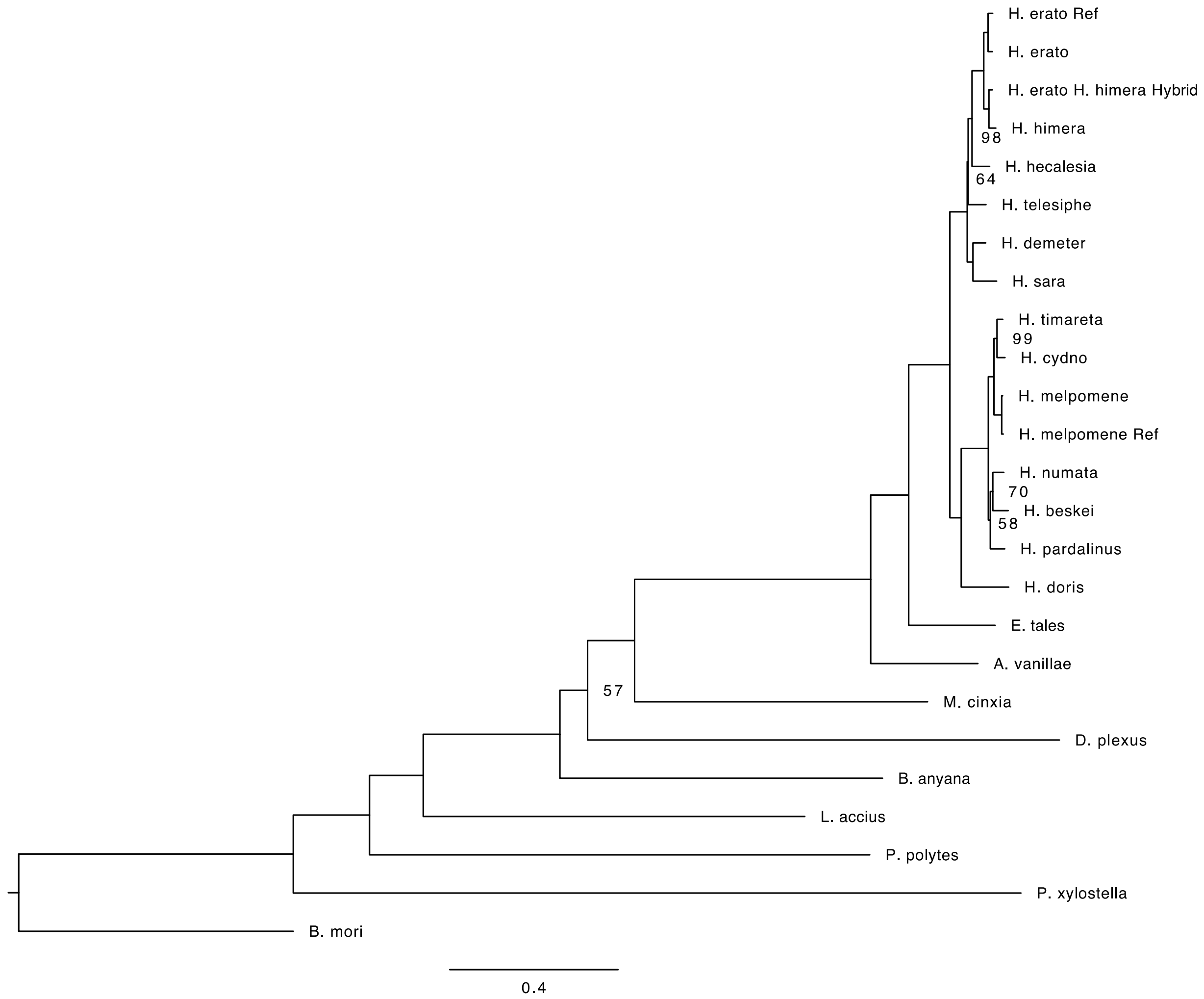

**Figure S4.5. Concatenated tree for noncoding, fully aligned blocks greater than or equal to 150 bp.** Bootstrap values are shown for nodes with less than 100% support.

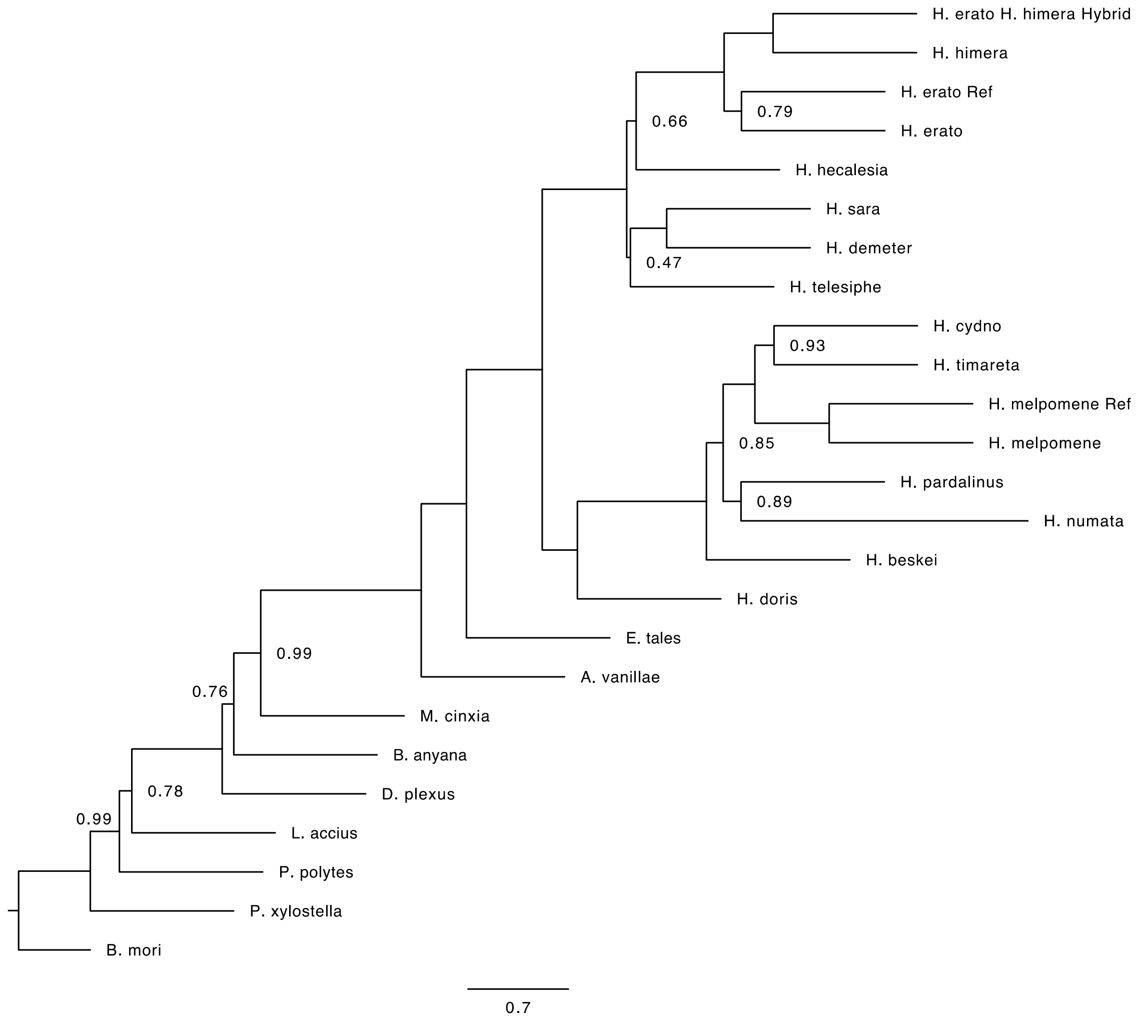

**Figure S4.6. ASTRAL tree for noncoding, fully aligned blocks greater than or equal to 150 bp.**

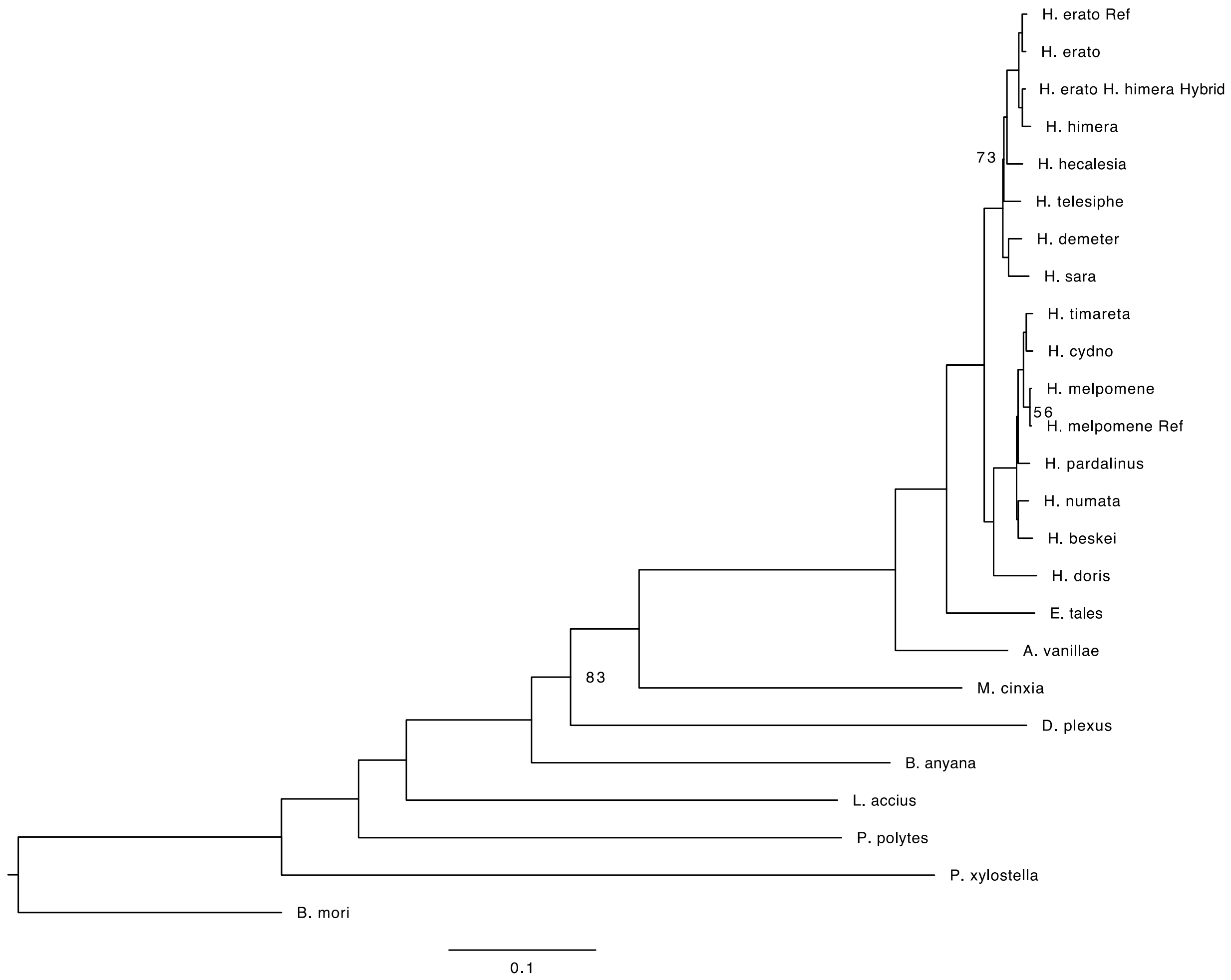

**Figure S4.7. Concatenated tree for coding, fully aligned blocks greater than or equal to 100 bp.** Bootstrap values are shown for nodes with less than 100% support.

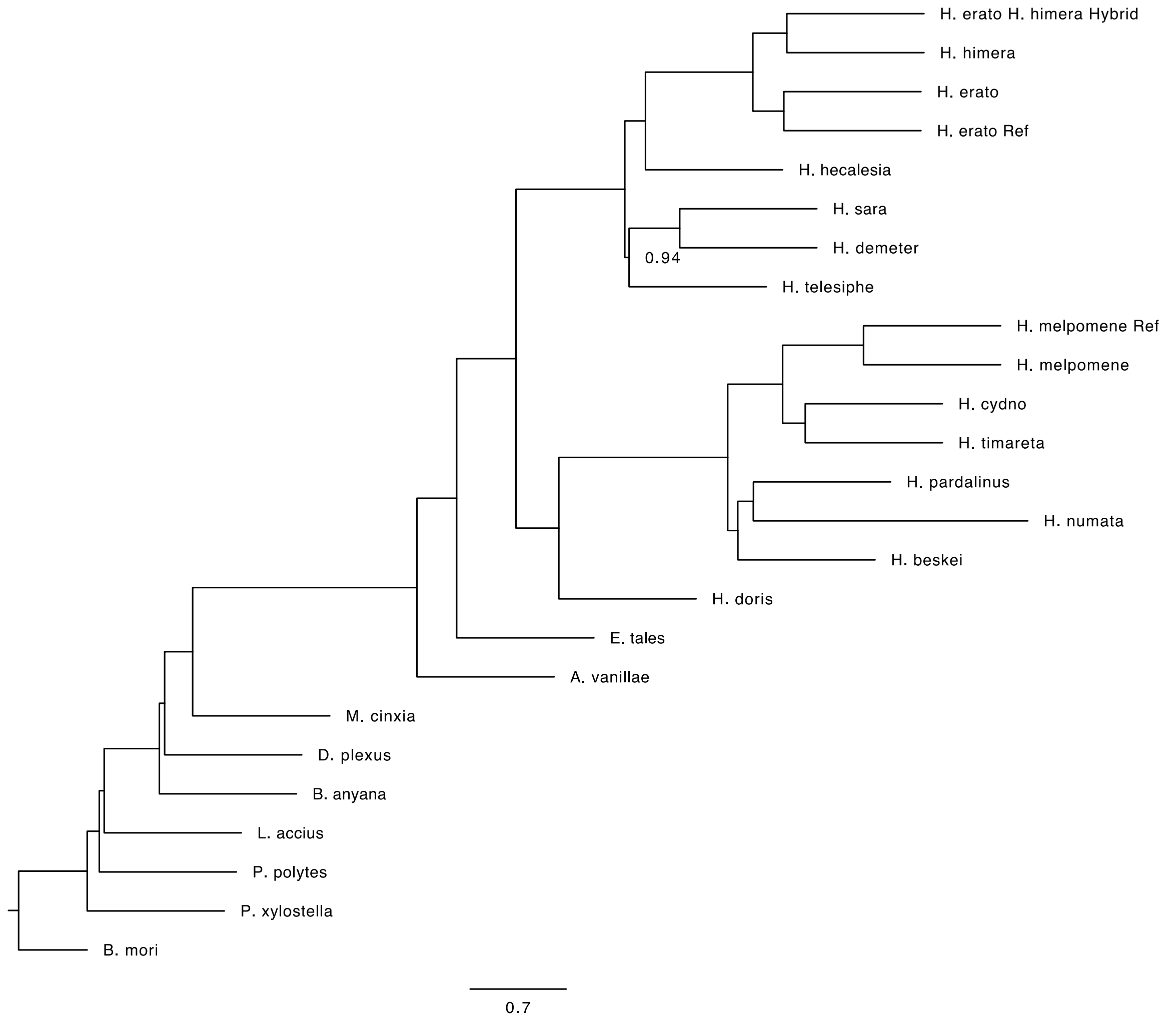

**Figure S4.8. ASTRAL tree for coding, fully aligned blocks greater than or equal to 100 bp.**

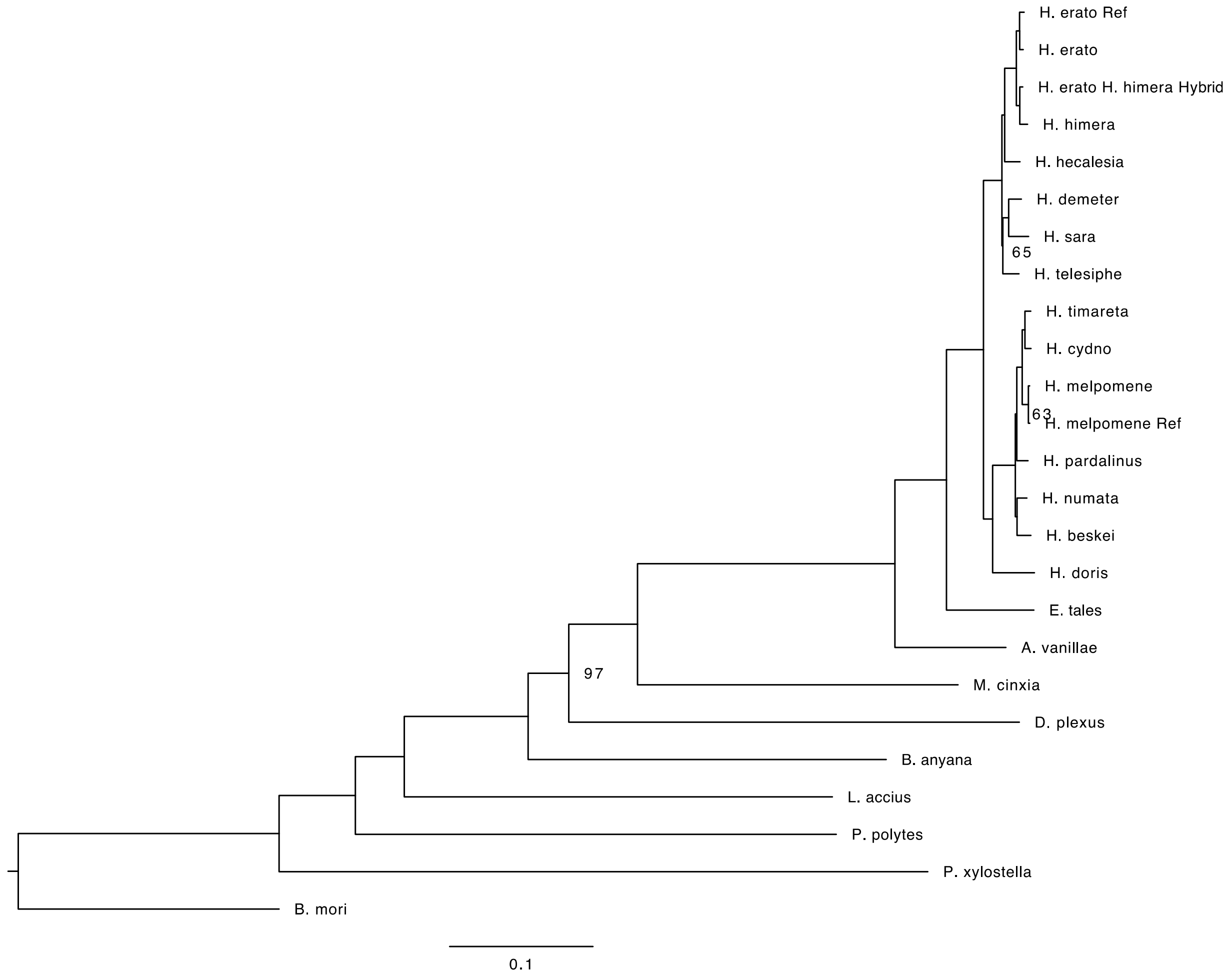

**Figure S4.9. Concatenated tree for coding, fully aligned blocks greater than or equal to 150 bp.** Bootstrap values are shown for nodes with less than 100% support.

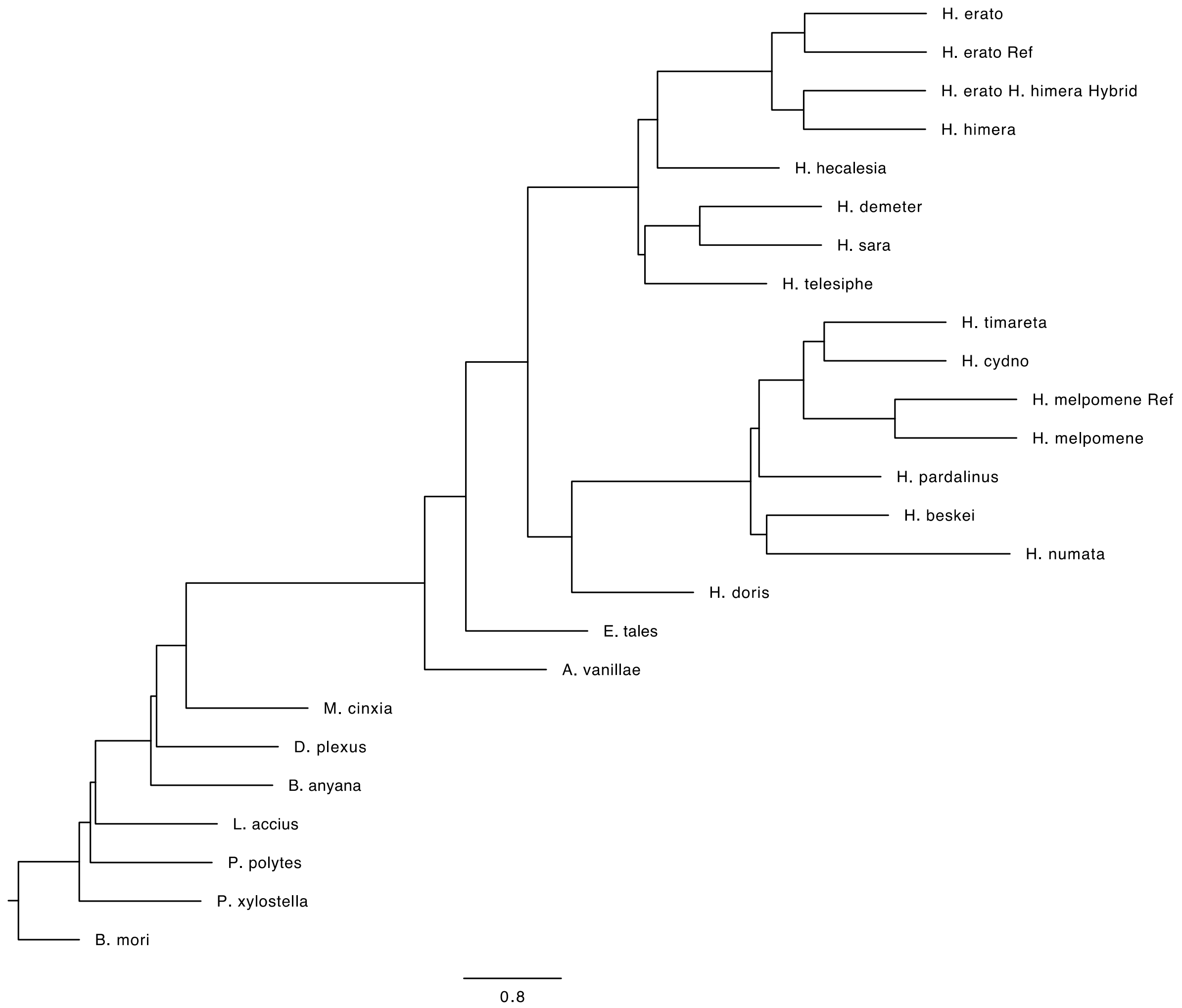

**Figure S4.10. ASTRAL tree for coding, fully aligned blocks greater than or equal to 150 bp.**

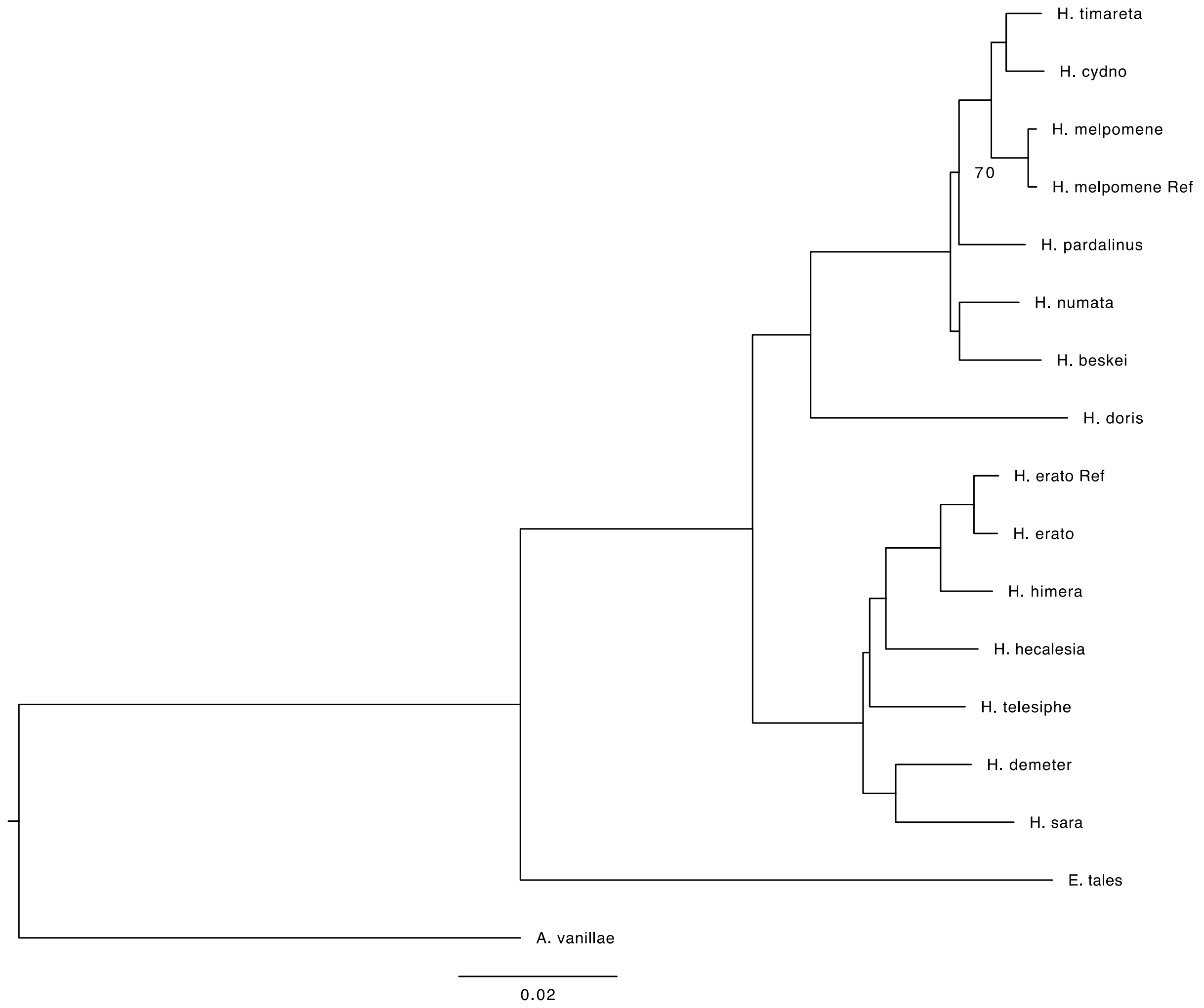

**Figure S4.11. Concatenated tree for noncoding, fully aligned blocks among Heliconiini that are greater than or equal to 100 bp.** Bootstrap values are shown for nodes with less than 100% support.

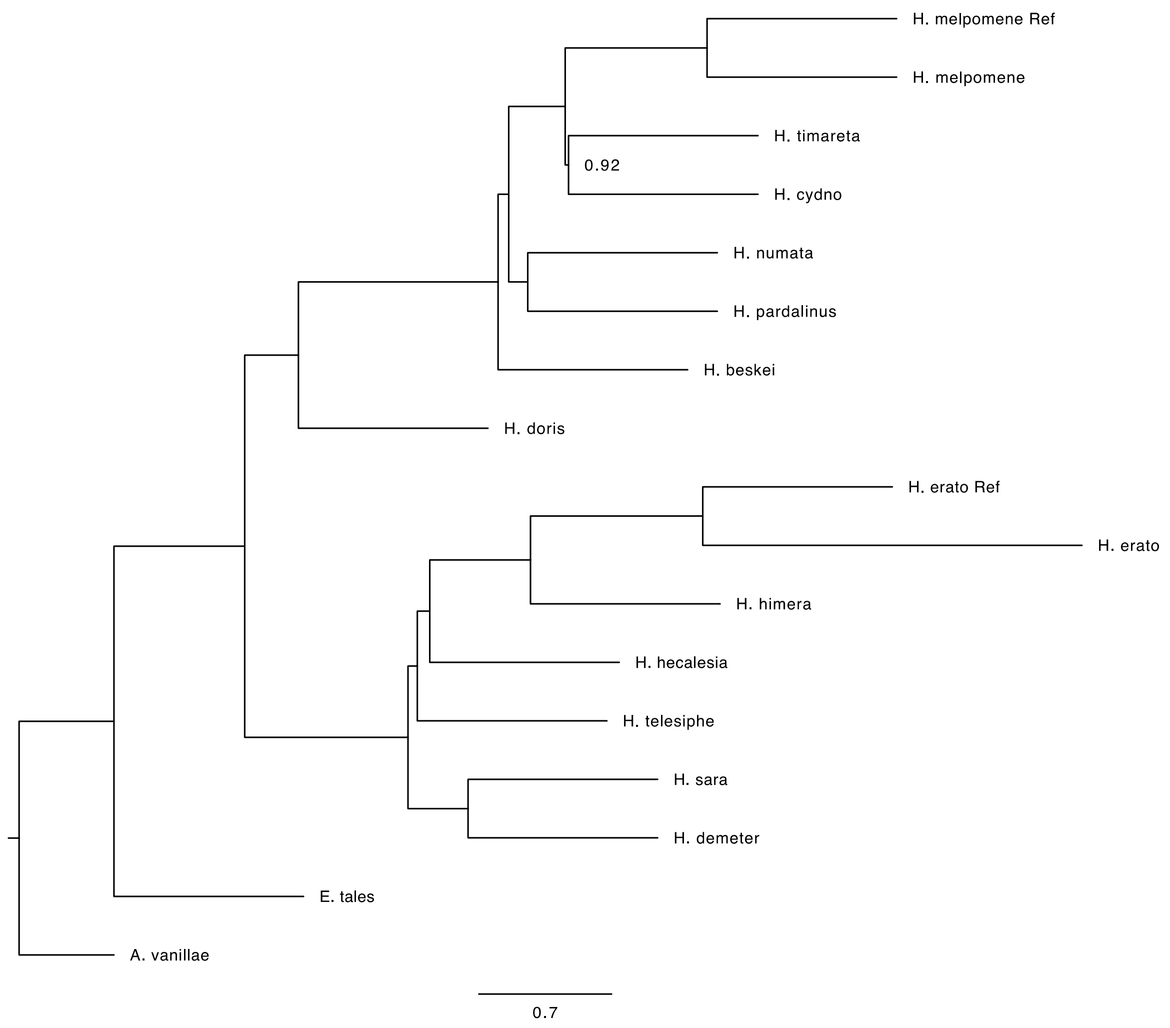

**Figure S4.12. ASTRAL tree for noncoding, fully aligned blocks among Heliconiini that are greater than or equal to 100 bp.**

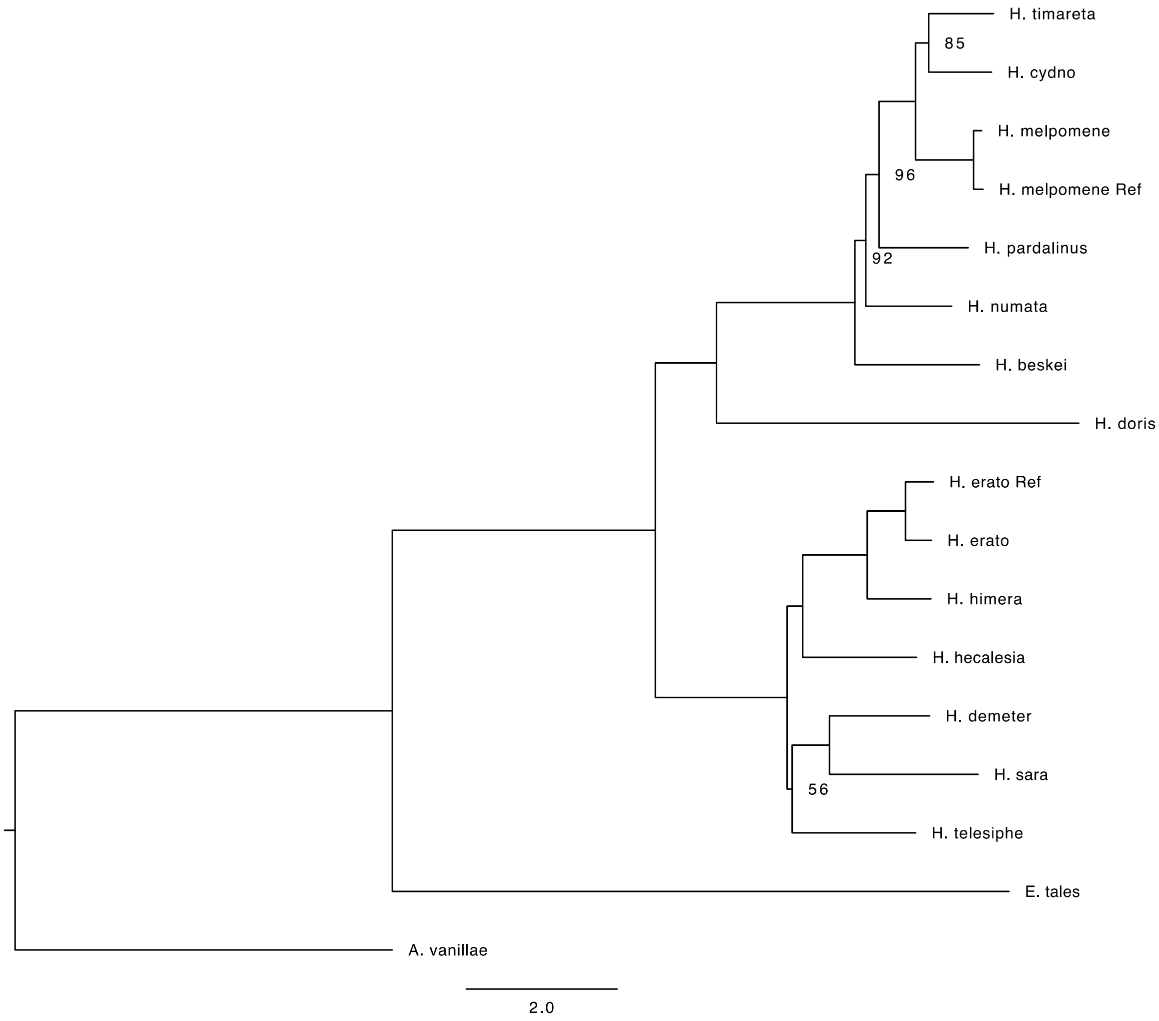

**Figure S4.13. Concatenated tree for noncoding, fully aligned blocks among Heliconiini that are greater than or equal to 150 bp.** Bootstrap values are shown for nodes with less than 100% support.

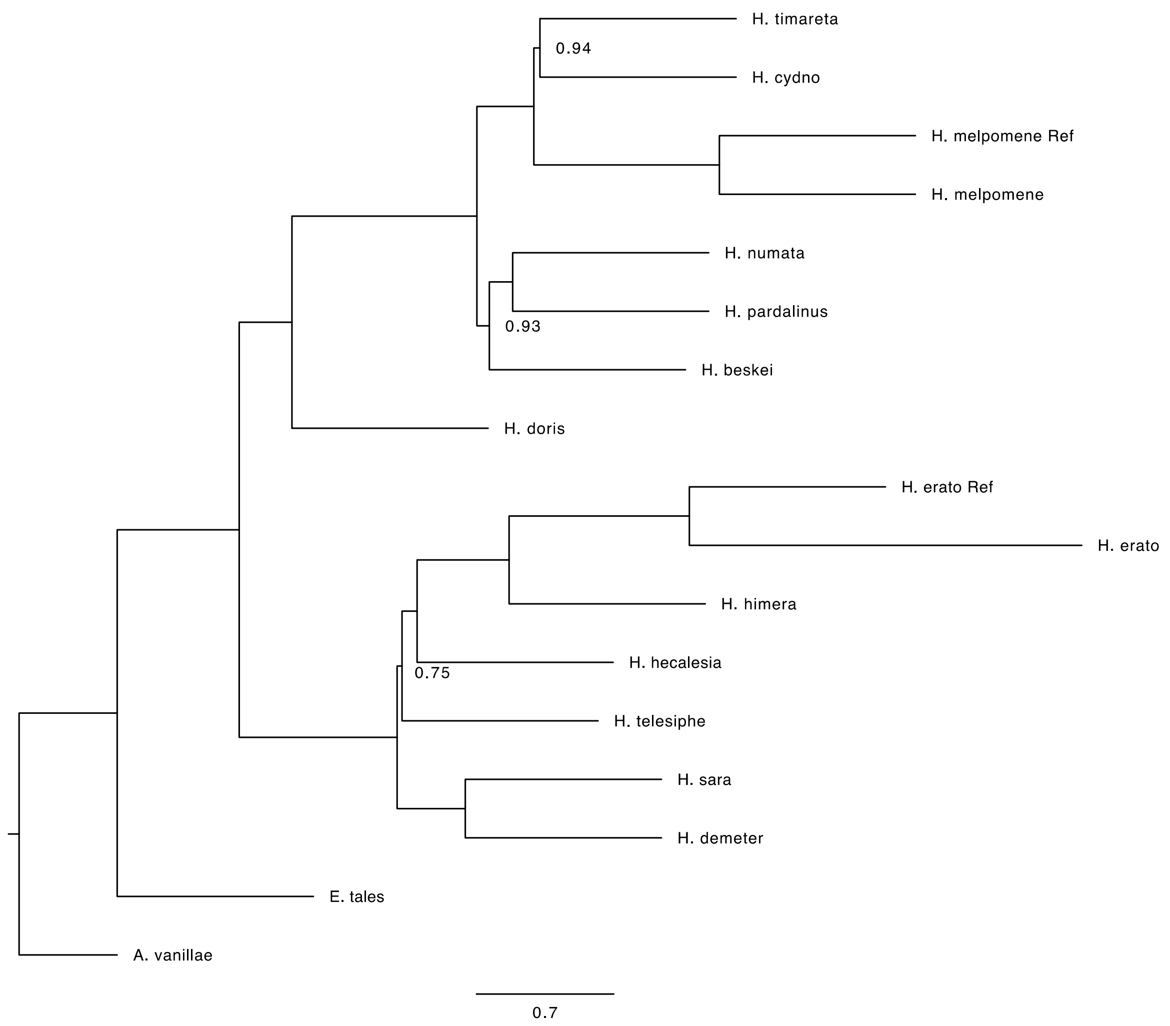

**Figure S4.14. ASTRAL tree for noncoding, fully aligned blocks among Heliconiini that are greater than or equal to 150 bp.**

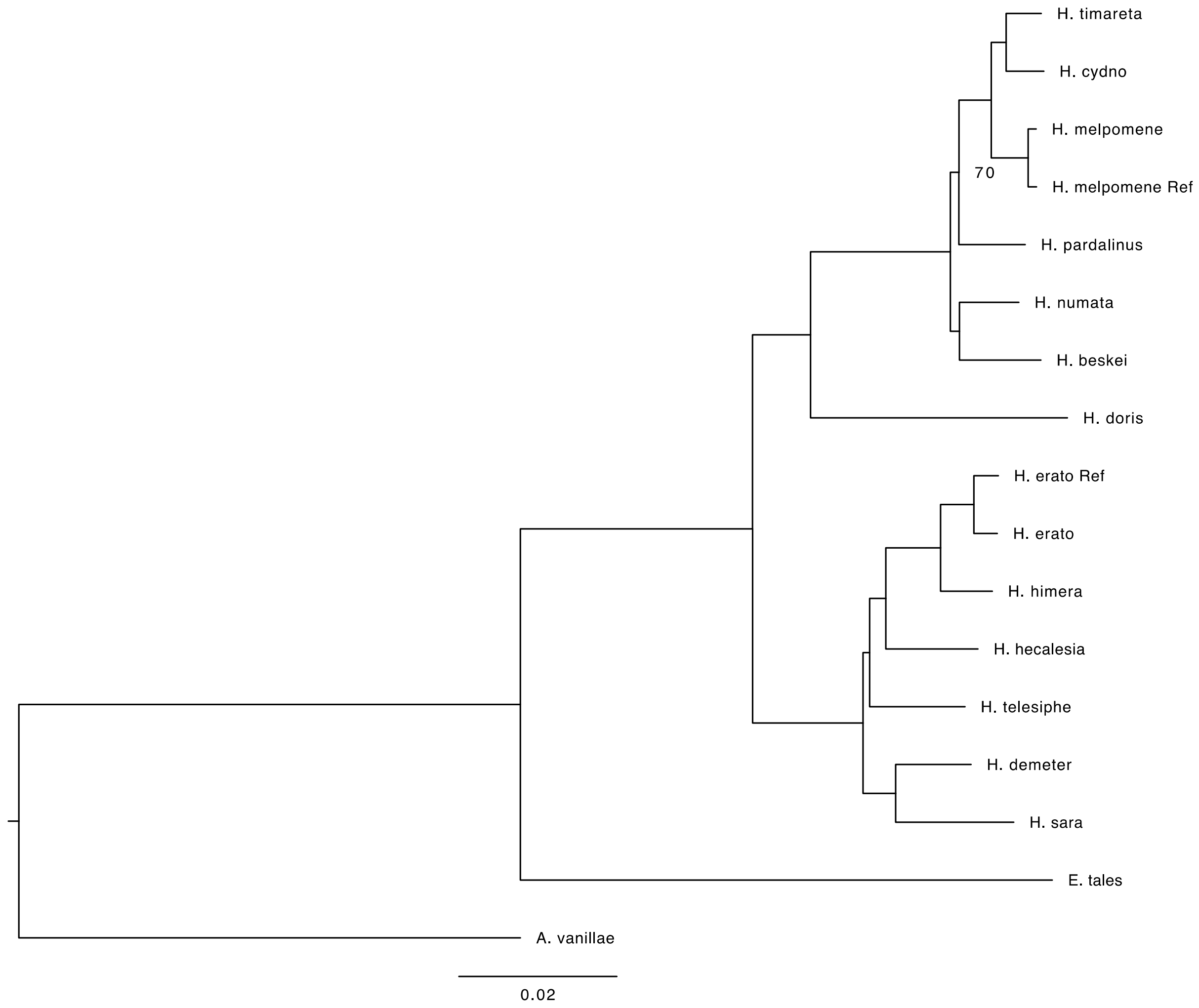

**Figure S4.15. Concatenated tree for coding, fully aligned blocks among Heliconiini that are greater than or equal to 100 bp.** Bootstrap values are shown for nodes with less than 100% support.

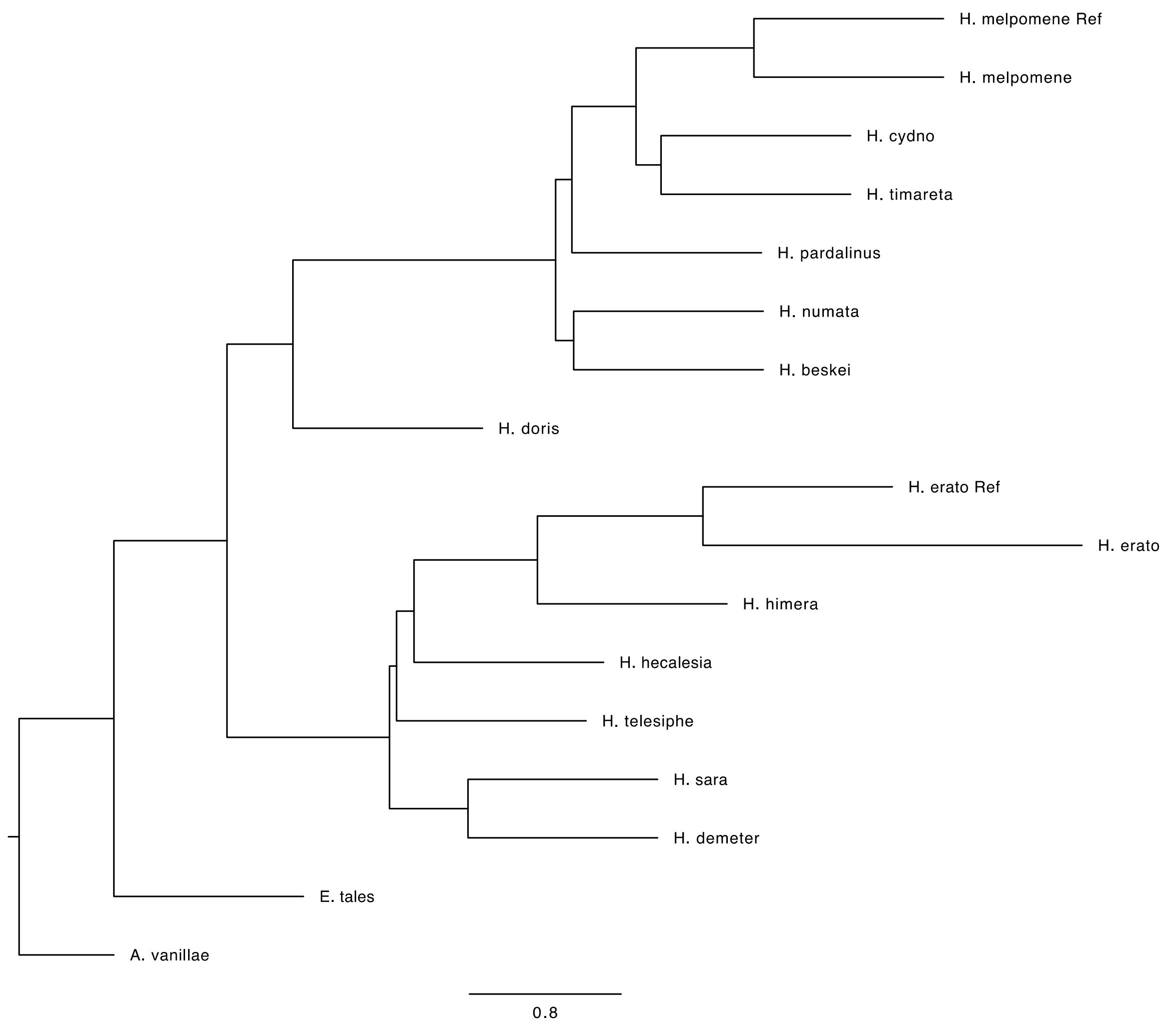

**Figure S4.16. ASTRAL tree for coding, fully aligned blocks among Heliconiini that are greater than or equal to 100 bp.**

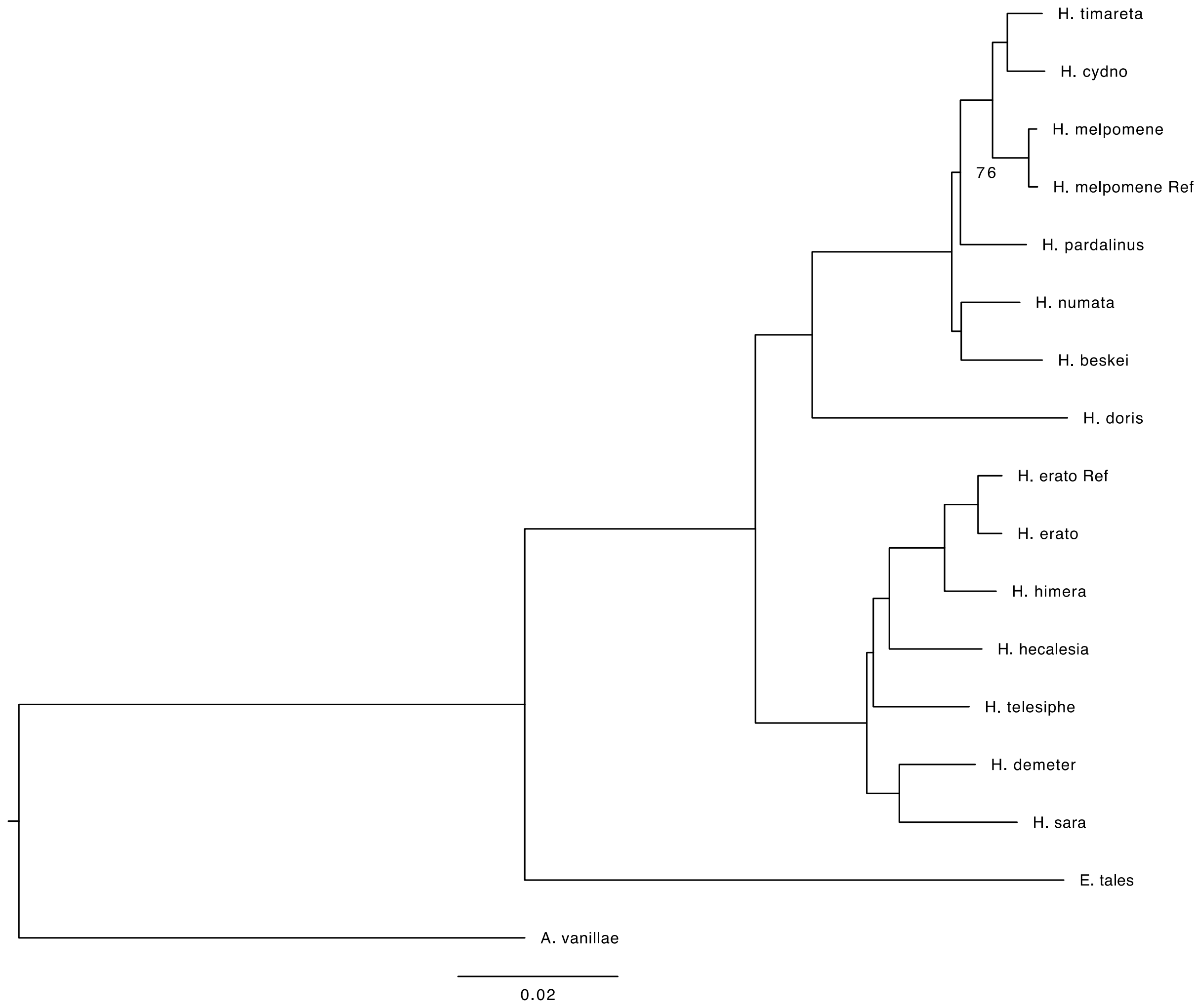

**Figure S4.17. Concatenated tree for coding, fully aligned blocks among Heliconiini that are greater than or equal to 150 bp.** Bootstrap values are shown for nodes with less than 100% support.

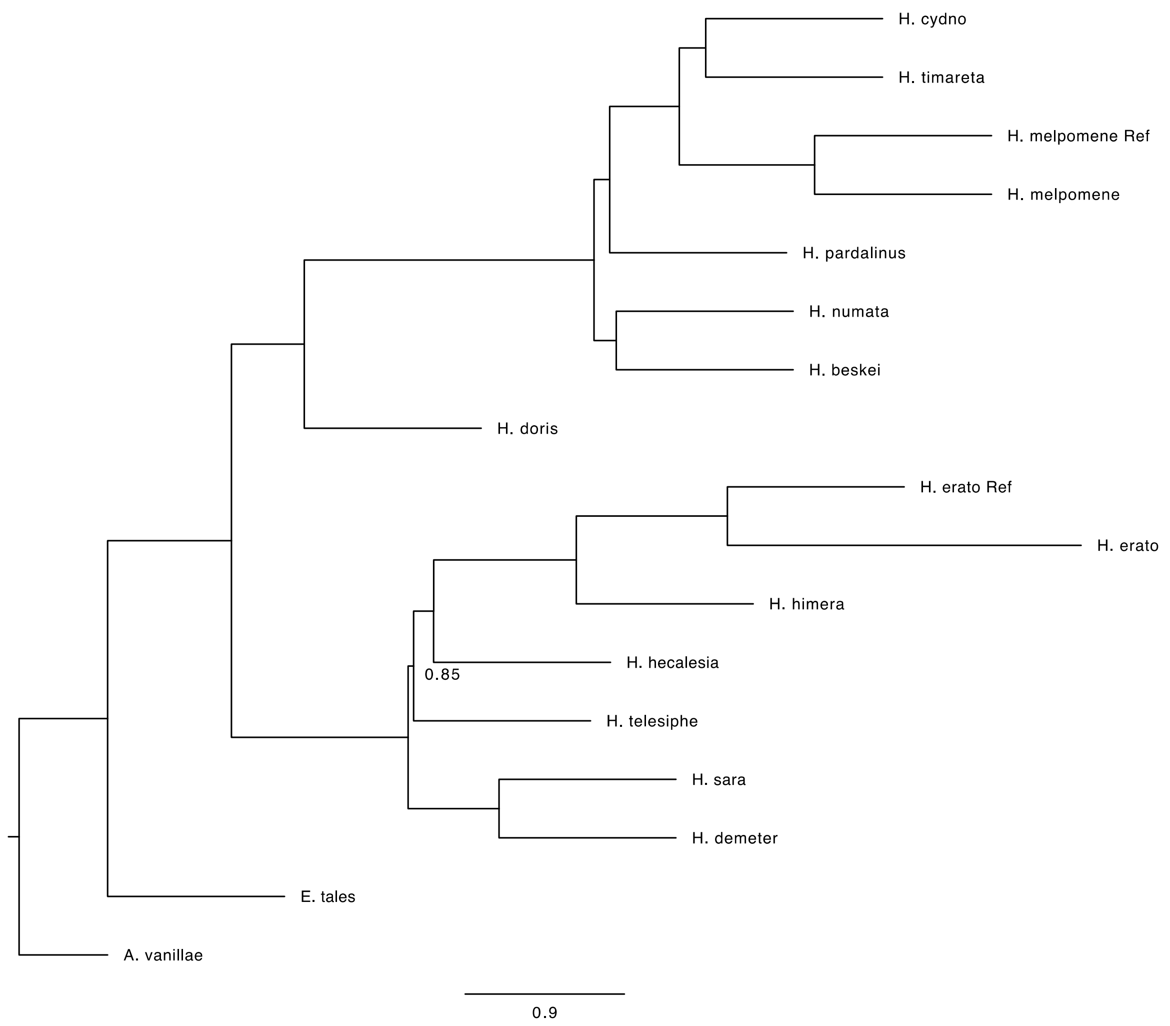

**Figure S4.18. ASTRAL tree for coding, fully aligned blocks among Heliconiini that are greater than or equal to 150 bp.**

**Section 5: D-statistics**
[return to top](#)

As stated in the main text, we calculated D-statistics for all triplets of *Heliconius* species, holding *Eueides tales* constant as the outgroup. For each triplet, we calculated the D-statistic for all three possible species topologies. The distribution of the lowest D-statistic score for each triplet are shown in Figure S5.1, and the corresponding Z-scores are shown in figure S5.2.

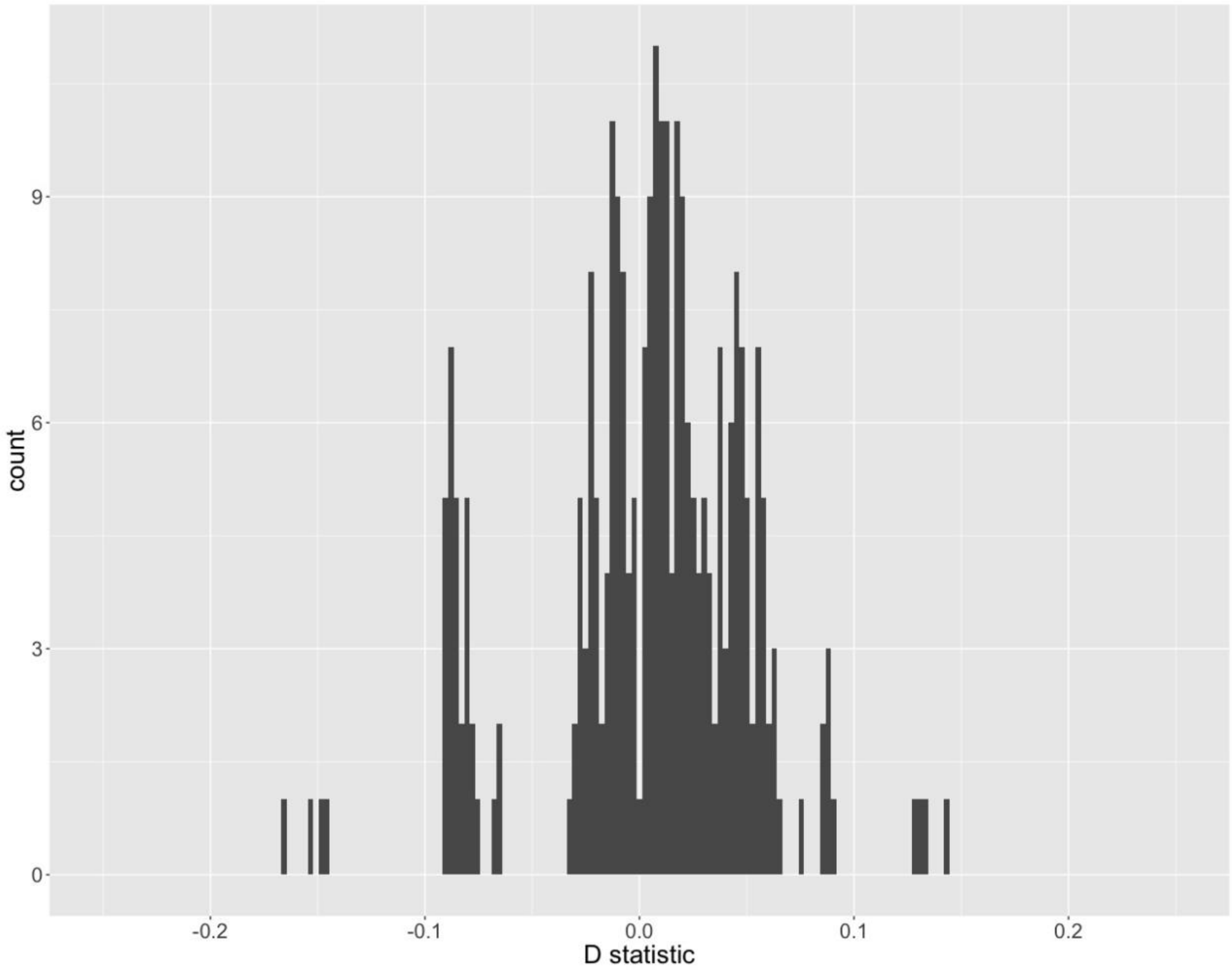

**Figure S5.1: D-statistic values for all triplets.**

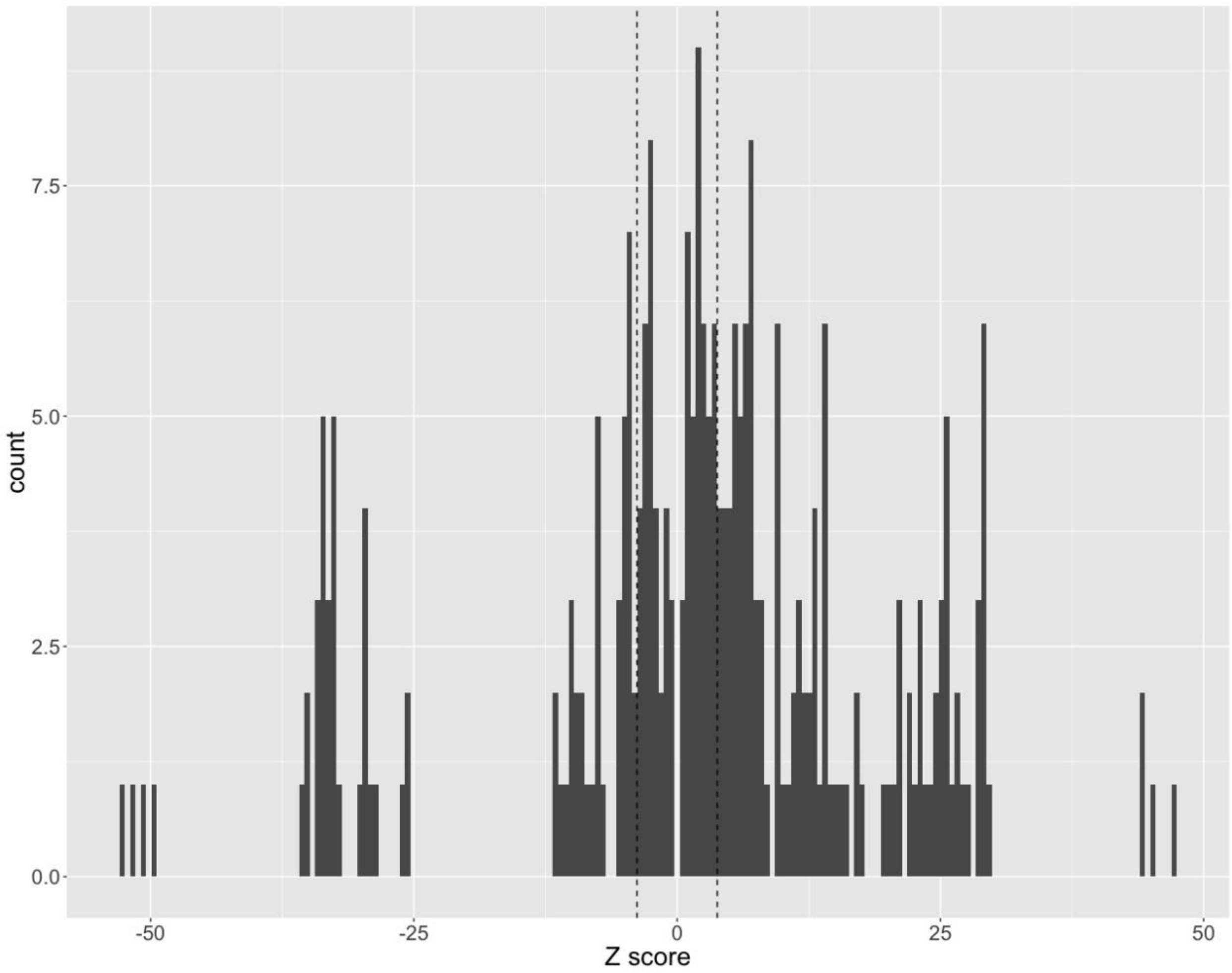

**Figure S5.2: Z-values for D statistics for all triplets**
This histogram shows the Z-values for all comparisons shown in Figure S5.1. The dotted lines represent significance at a .05 level after applying a bonferroni correction.

To ensure our analysis was not biased due to aligning the genomes with a guide tree, we calculated D-statistics for the same set of species using an alternative progressiveCactus alignment with a guide tree that reflects a different order of branching in the uncertain silvaniform clade[1]. In addition, the two alignments differed in species composition. The "LepBase" alignment included all genome assemblies, including those of low quality (accessed from lepbase.org). The values for the two alignments within each clade were very highly correlated, indicating no strong effect of the input guide tree on these results (Figure S5.3, green points). Across clades, the correlation was not perfect, particularly when the *H. doris* genome was one of the samples (Table S5.1, Figure S5.3, blue points). Anomalies over longer evolutionary distances to be expected, as the D statistic was built to examine very similar populations and assumes only a single mutation per site. In addition, the deviation did not affect our downstream analyses, as we focused on within-clade alignments.

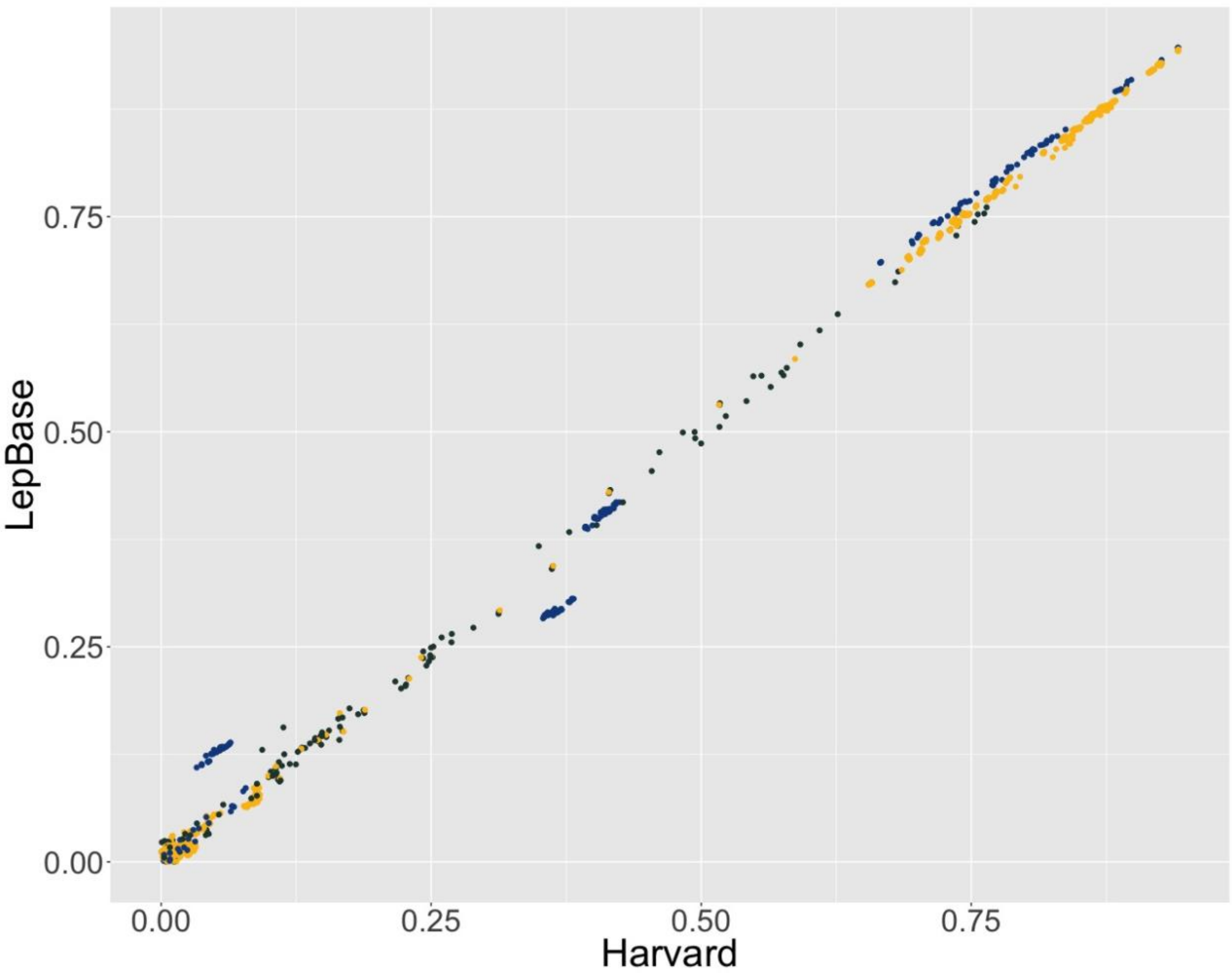

**Figure S5.3: D statistics are consistent between alignments**
These plots show a comparison of the absolute value of D-statistics computed from the full-Lepidoptera alignment described here (x-axis, "Harvard"), and the *Heliconiini* alignment generated by lepbase.org (y-axis, "LepBase"). Dark green points are inter-clade comparisons, yellow are between clade, and blue are those involving *H. doris*. Both groups of *H. doris* points off the 1:1 line are inter-clade comparisons. Values are shown for all triplet topologies, not only the lowest value per triplet.

**Section 6: Phylogenetic Networks**

*return to top*

As stated in the main text, we used the MCMC_Seq method of PhyloNet to generate phylogenetic networks for the *erato* and *melpomene* clades separately. We started by running 100 iterations of the program using 100 randomly sampled loci each time. We considered all runs in which the final output network had greater than 50% posterior probability, and summarized the results by generating a correlation matrix of those networks based on Luay Nakhleh's metric of reduced phylogenetic network similarity[5]. We calculated each pairwise similarity score, and then used the R heatmap function to group all trees by similarity (Figures S6.1-S6.2. For the *melpomene* clade, there was a single region of tree-space that was strongly supported (main text, Figure 1C). However, the *erato* clade was less clear, and we therefore repeated the analysis using 150 loci per run and doubled the chain length. Still, several regions of tree space seemed equally supported. Upon close examination, many of the differences are simply in direction or timing of introgression events. The clearest structural difference (i.e. change to the backbone tree structure) was that roughly half of the networks infer a base tree that recovers *H. telesiphe* and *H. hecalesia* as a monophyletic clade that admixed with the ancestor of *H. erato* and *H. himera.* The other half recovers *H. hecalesia* as sister to *H. erato* and *H. himera*, and identifies an introgression event either from *H. telesiphe* into *H. hecalesia* or vice versa. The remainder of the uncertainty lies in the timing of the introgression event into the ancestor of *H. sara* and *H. demeter.* Various networks place it at all possible branches of the tree with the exceptions of the *H erato* and *H. himera* terminal branches.

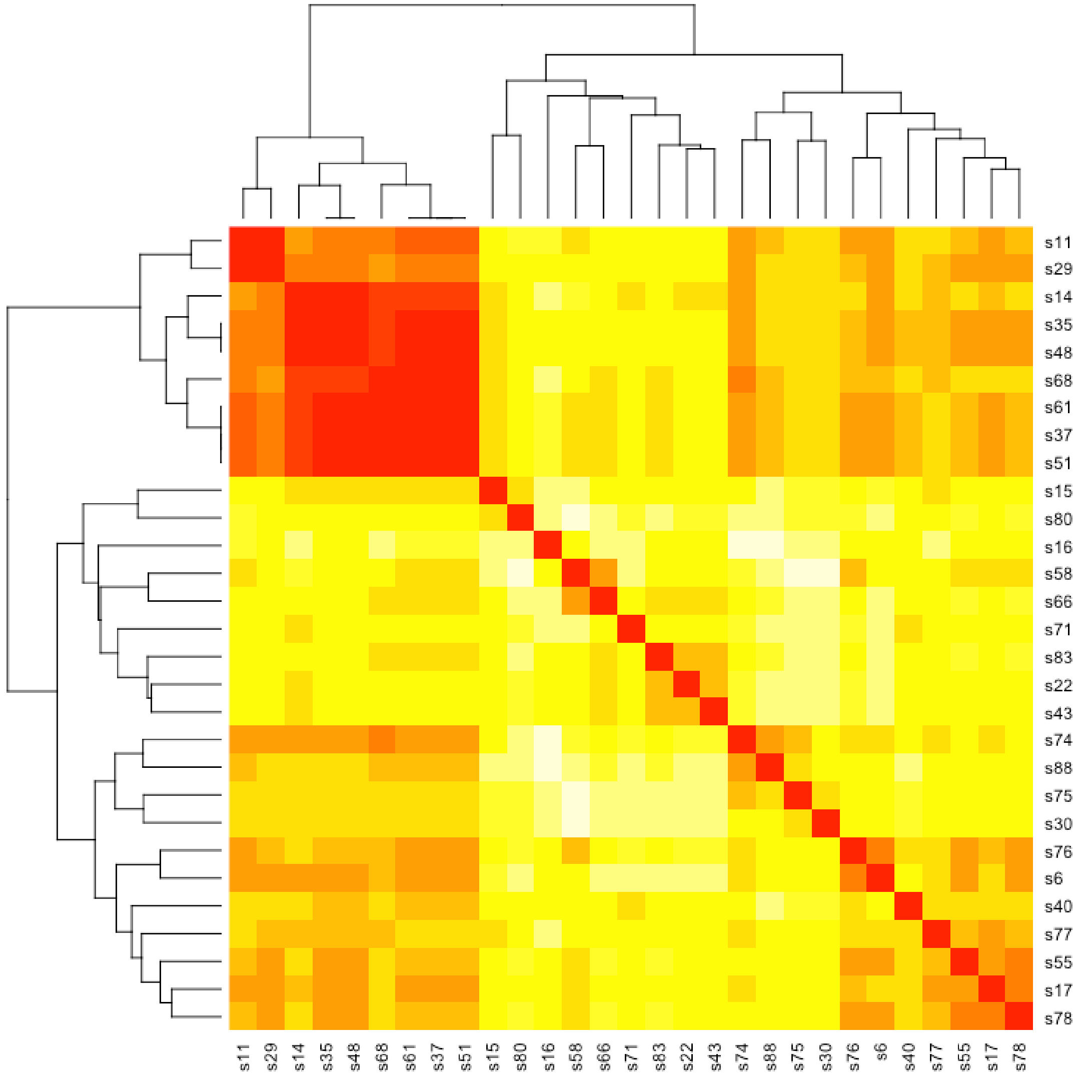

**Figure S6.1 *melpomene*-silvaniform clade phylogenetic network clustering.**
Matrix is symmetrical, and each row or column represents the most likely network in a single run of PhyloNet MCMC_Seq. The branch lengths of the dendrograms represent Luay Nakhleh's reduced phylogenetic network distance metric, where 0 means the networks are identical. The top left section is populated by 9 very similar networks, while the remaining 20 networks are quite different from one another. The consensus network shown in the main text Figure 1C was generated by summarizing the contents of the networks in the top left corner.

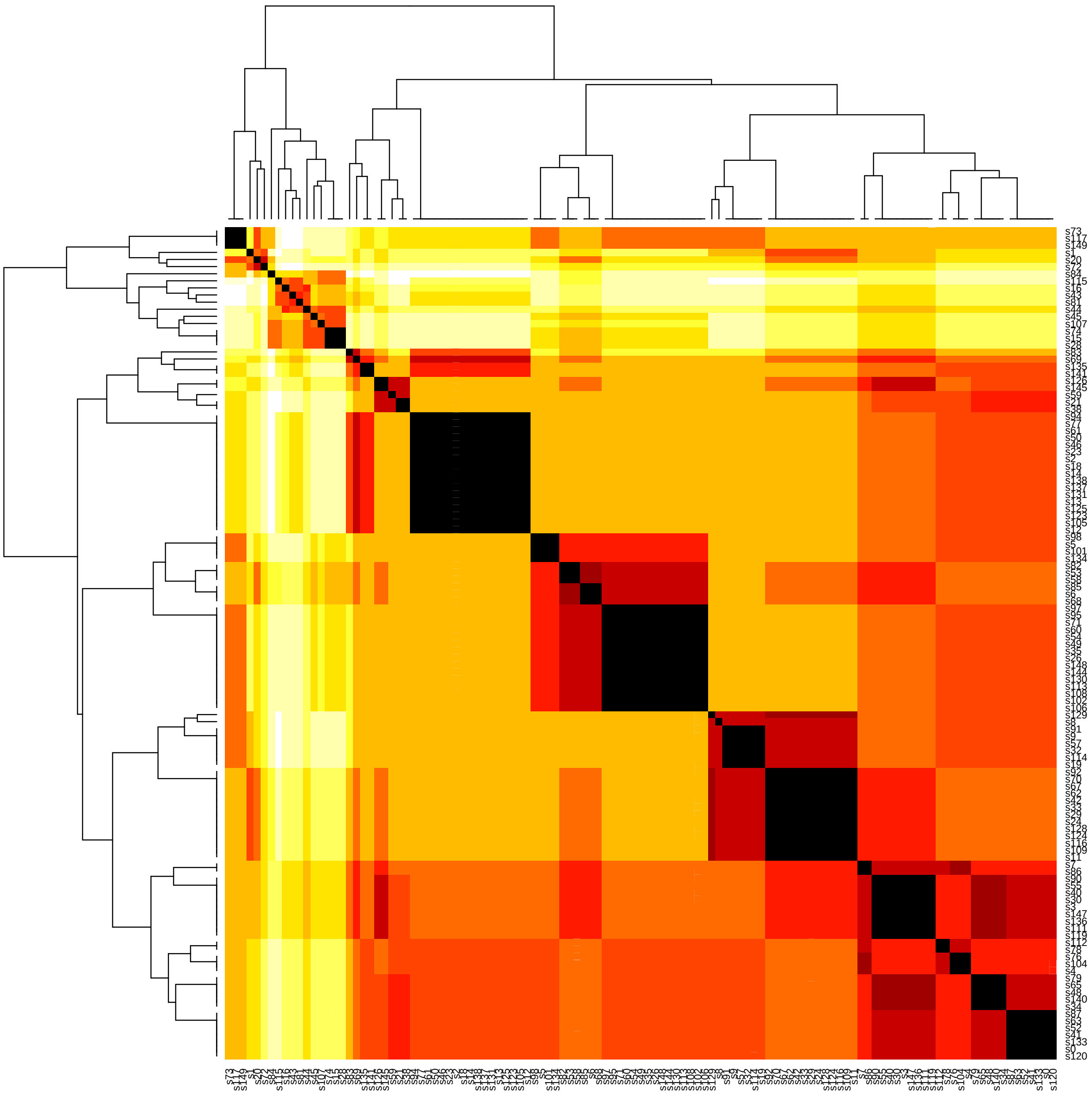

**Figure S6.2 *erato-sara* clade phylogenetic network clustering.**
Format of the matrix is identical to that in Figure S6.1. We identify four well-represented regions of tree space among *erato* clade PhyloNet MCMC_Seq runs.

We generated consensus phylogenetic networks by analyzing all networks in the supported region(s) of tree space. For the *melpomene*-silvaniform clade, any connection present in a majority of networks in the single well-represented region was drawn as a solid line, and any connection present in a minority of those networks was drawn as a dashed line. In the *erato*

clade, we drew the most common network as solid lines, and added dashed lines to represent introgression events supported by other well-represented regions of tree space.
We next aimed to test the robustness of the network results by repeating our analysis with other inference methods. A major drawback of the MCMC_Seq method above is its limitation to small datasets, so we only used a subset of available data in each run, and were only able to combine them after running each set. We therefore used another PhyloNet method, Infer_Network_MPL, to corroborate the result. This method uses all loci, but uses inferred gene trees as input data as opposed to sequence alignments[19,20]. The method estimates the network by maximizing pseudolikelihood scores based on those input trees. We inferred gene trees using PhyML[18] from the same set of loci used in the MCMC_Seq method. In both clades, the major introgression events identified by MCMC_Seq were also found by Infer_Network_MPL, though not always in the same direction (i.e. *H. besckei-H. numata* in the *melpomene*-silvaniform clade; *H. sara/H. demeter - H. telesiphe/H.hecalesia* in the *erato-sara* clade) (Fig. S. 6.3).

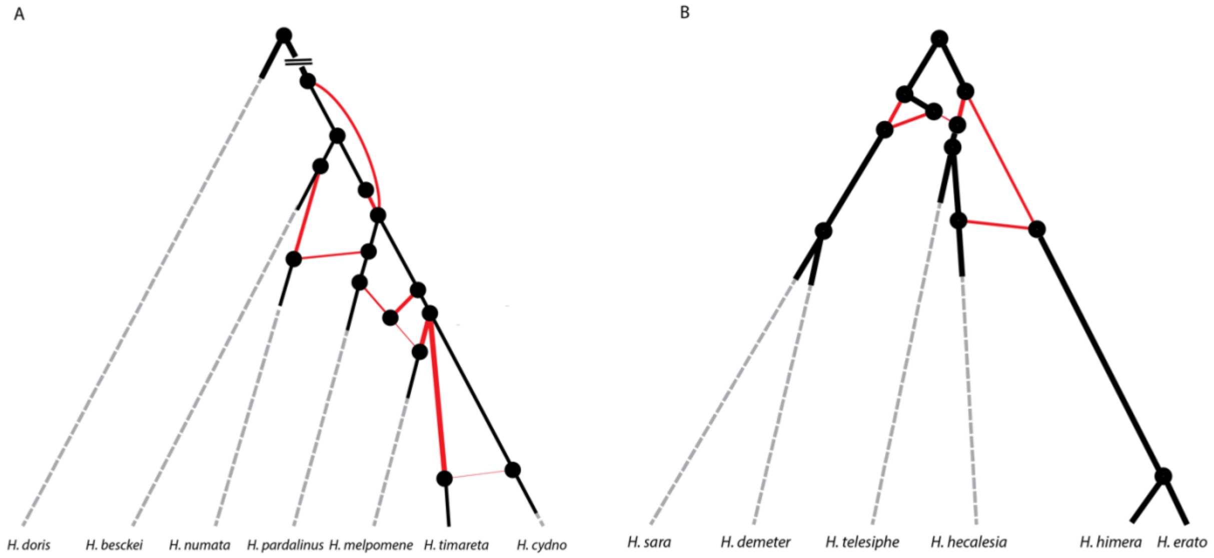

**Figure S6.3: phylogenetic networks using PhyloNet Infer_Network_MPL.**
***A.*** *melpomene*-silvaniform clade network.
**B.** *erato-sara* clade network

Finally, we used the biallelic SNP dataset generated for our D-statistic analysis to generate a phylogenetic network with qpGraph[21]. We iteratively added taxa to a model according to the following algorithm:

*1. Begin with 3 species that have a well-defined, bifurcating history.*
*2. Add one species ("test taxon") to the tree in its most likely location based on previously published phylogenies and D-statistic values.*
*3. Run qpGraph.*

*4. If all statistics are equal to 0, return to 2. If not, continue to 5.*
*5. If the test taxon can be moved to a different location without adding a reticulation node, continue to 6. If not, move to 8*
*6. Move the test taxon to a different location without adding a reticulation node. The position should be informed by the output of qpGraph.*
*7. Return to 3.*
*8. Add a reticulation node whose position is informed by the output of qpGraph.*
*9. Return to 3.*

Continue until all species are included in the phylogenetic network.

This method was successful for the *erato* clade (Figure S6.4), because sister species relationships between *H. erato* and *H. himera*, as well as *H. sara* and *H. demeter*, were generally stable. This allowed us to build a backbone tree and vary the placements and admixture events involving *H. hecalesia* and *H. telesiphe* until we had a reasonably well-supported tree. These admixture events included the previously supported events between *H. hecalesia* and *H. telesiphe*, and between *H. telesiphe* and *H. sara+H. demeter*. The backbone tree differs slightly from that inferred by either PhyloNet method, most notably by the placement of *H. hecalesia* as sister to *H. sara+H. demeter*. However, a significant amount of *H. hecalesia* genetic material is inferred to trace its ancestry to its more classical placement as sister to *H. erato+H. himera.* We were also able to infer small admixture events involving *H. demeter.* This differs from the PhyloNet methods, which inferred admixture events involving the ancestor of *H. sara* and *H. demeter.*

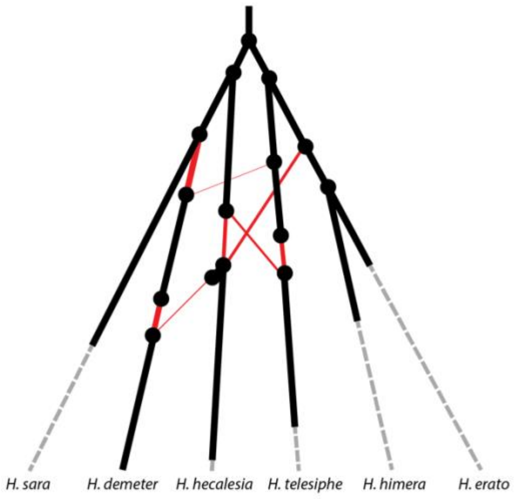

**Figure S6.4: *erato* clade phylogenetic network using qpGraph**

Unfortunately, we were not able to use the qpGraph method with the *melpomene* clade. No sister species relationships were stable, and the space of possible models proved too large to search manually.

**Section 7: Evolutionary Rate**
*return to top*

Our goal in this section was to identify candidate genes potentially involved in key innovations that catalyzed the *Heliconius* radiation. We therefore focused on changes that occurred in the branch leading from the common ancestor of *Heliconius* and *Eueides* to the common ancestor of *Heliconius.* We were interested in both coding sequence and regulatory evolution, so we opted to use the tool dless from the PHAST suite, which identifies changes in evolutionary rate along each branch of a phylogeny. Within *Heliconius*, there is uncertainty as to the correct branching order in several parts of the phylogeny (Supplementary Section 4), but the genus itself is strongly supported as a monophyletic group, and the branch of interest is stable. We generated a null model of evolutionary rate with Phylofit, another tool in the PHAST suite, using all annotated 4-fold degenerate sites in the genome. Significance in dless is calculated as a LOD score, the probability of the designated alignment segment under the “foreground model” (e.g., clade-specific conservation or acceleration) divided by the probability under the null model.

As stated in the main text, we identified 38,490 genomic regions that evolved at significantly faster or slower rates specifically along the branch leading to *Heliconius*, and their size distribution is shown in Figure S7.1. dless detected roughly equal numbers of changes in evolutionary rate (both conserved and accelerated regions) in intergenic and genic regions (Figure S7.2). Importantly, “genic” here refers to both exonic and intronic regions, as a single identified locus might cover both. However, the split between accelerated and conserved regions was far from equal. On the contrary, the vast majority of loci identified by dless as having changed evolutionary rate at the base of *Heliconius* displayed an accelerated rate (Figure S7.3). This may be due to a bias in the power of this analysis when projecting the alignment onto *H. melpomene*. When most *Heliconius* species appear conserved for a locus but the outgroups do not, it may be difficult to distinguish between true enhanced conservation and poor alignment due to evolutionary distance. In contrast, regions that are strongly conserved among all outgroups but less conserved within *Heliconius* are more easily interpretable.

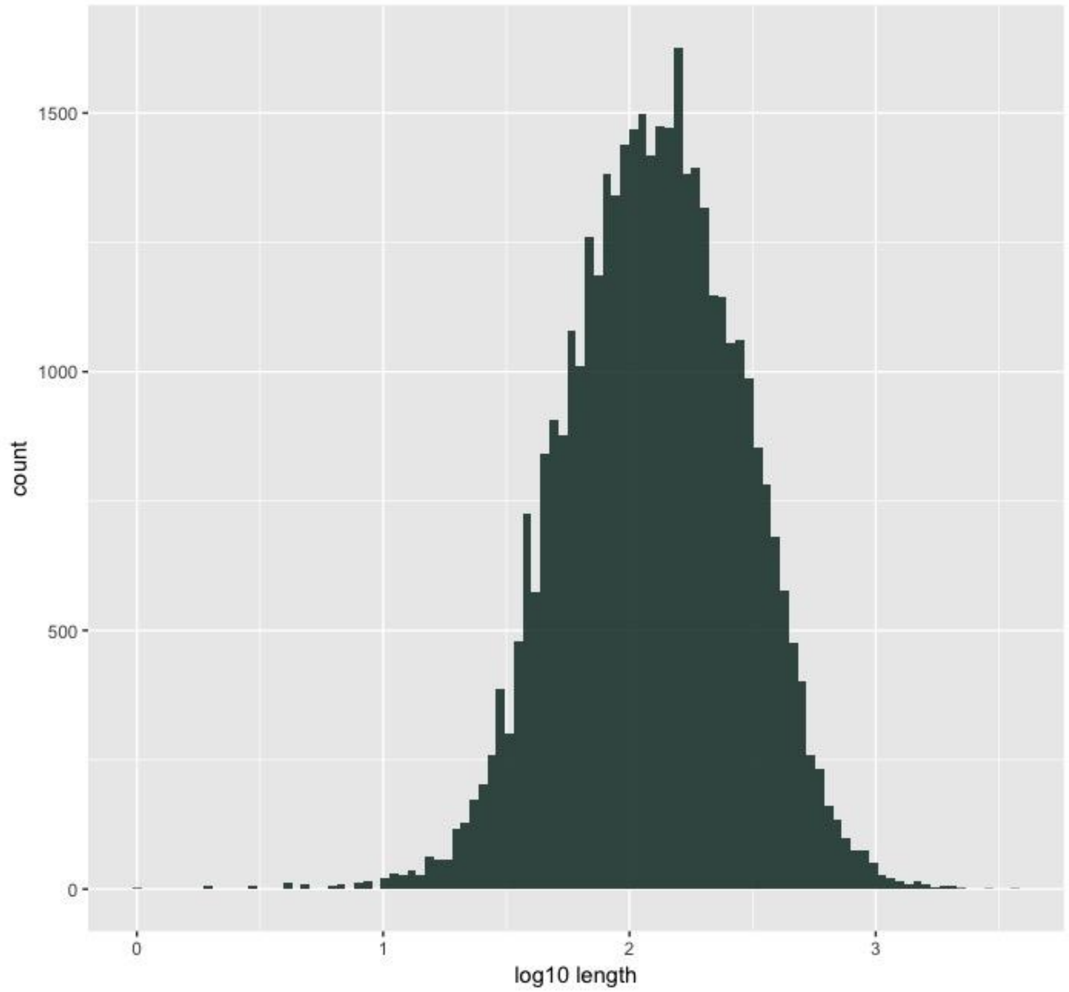

**Figure S7.1: Length of rate-changed loci**
This graph displays the distribution of locus lengths for regions identified by dless as having an altered evolutionary rate at the base of *Heliconius*. The x-axis is the $\log_{10}$ length of the locus.

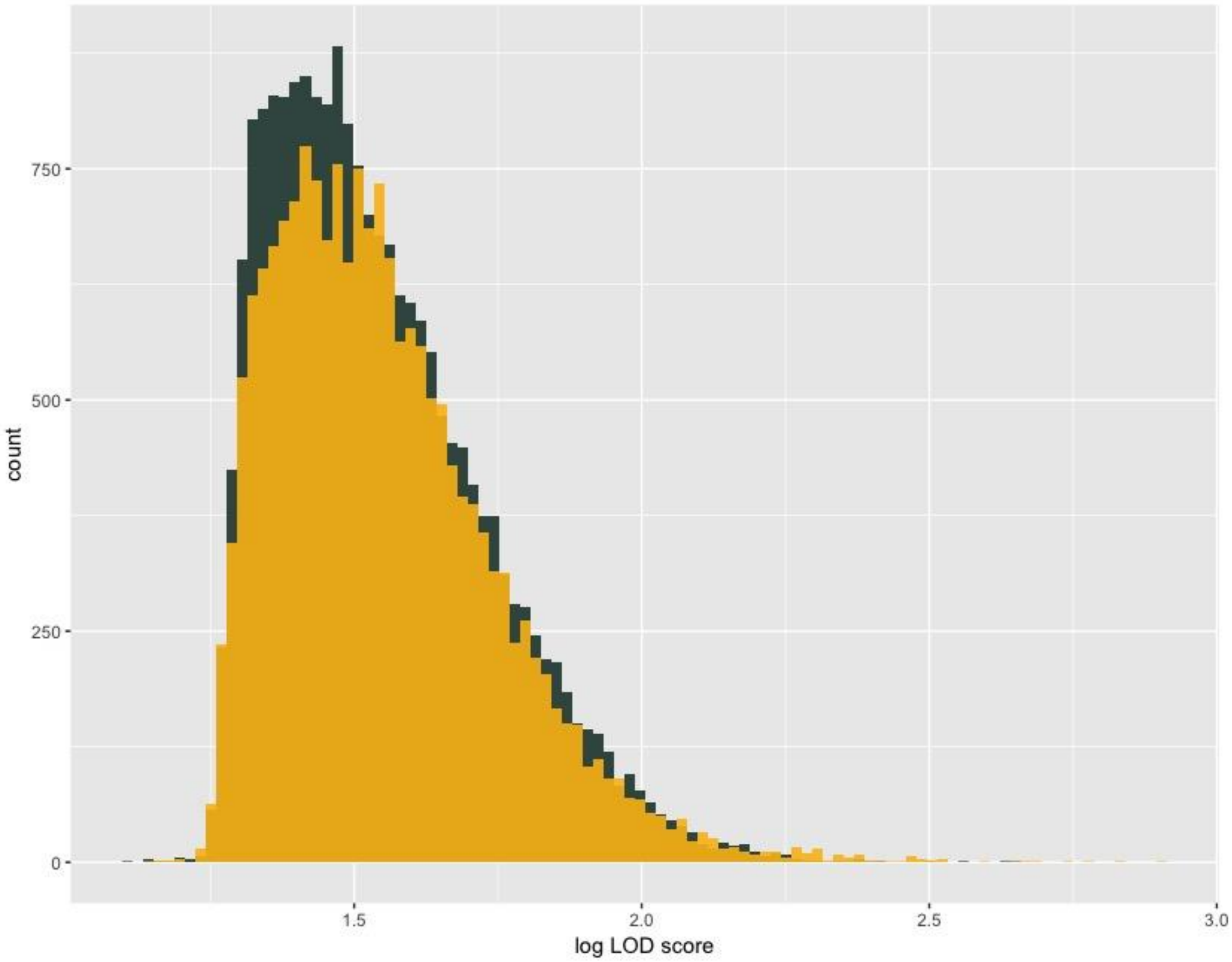

**Figure S7.2: Genic and intergenic loci have similar distributions of evolutionary rate change**
The x-axis is the $\log_{10}$ transformation of the LOD score (in order to better visualize the skewed distribution), calculated as the probability of the designated alignment segment under the "foreground model" (e.g., clade-specific conservation or acceleration) divided by the probability under the null model.

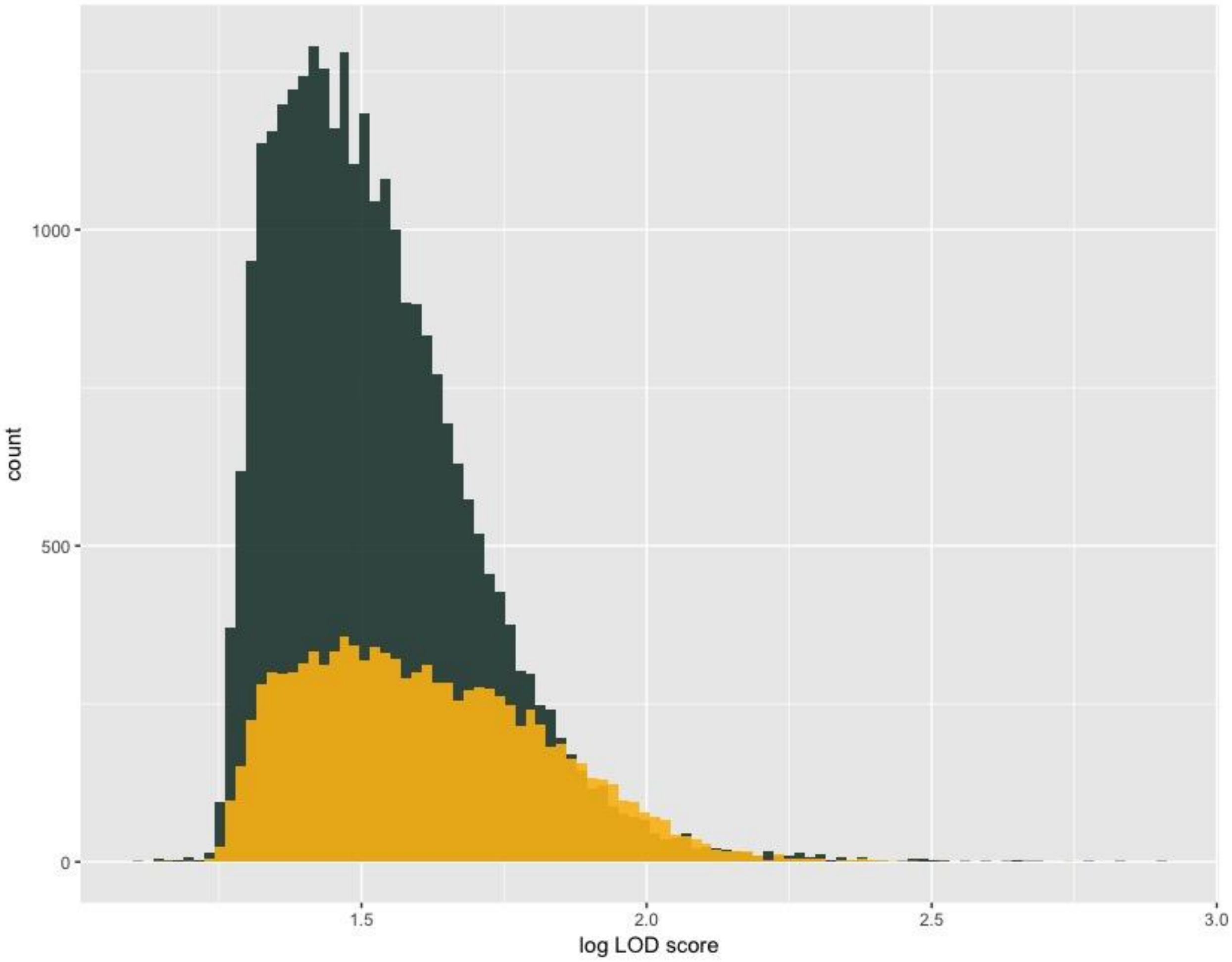

**Figure S7.3: Distribution of accelerated vs conserved regions**
Accelerated regions are shown in green, and conserved in gold. The x-axis is the $\log_{10}$ of the LOD score.

Finally, we examined regions around known color pattern loci in more detail. While there are regions that show changes in evolutionary rate around *wntA*, *cortex*, and *optix*, note the scale is on the order of 10-100, while the highest conserved and accelerated genomic regions have LOD scores above 300, with a maximum of 817. Nevertheless, significant loci identified in non-coding regions are candidates for *Heliconius*-specific gene expression evolution, and should be studied further. A list of dless hits with the highest LOD scores for genic, intergenic, accelerated, and conserved loci is shown in Table S7.1, and a full list of hits can be found on Dryad.

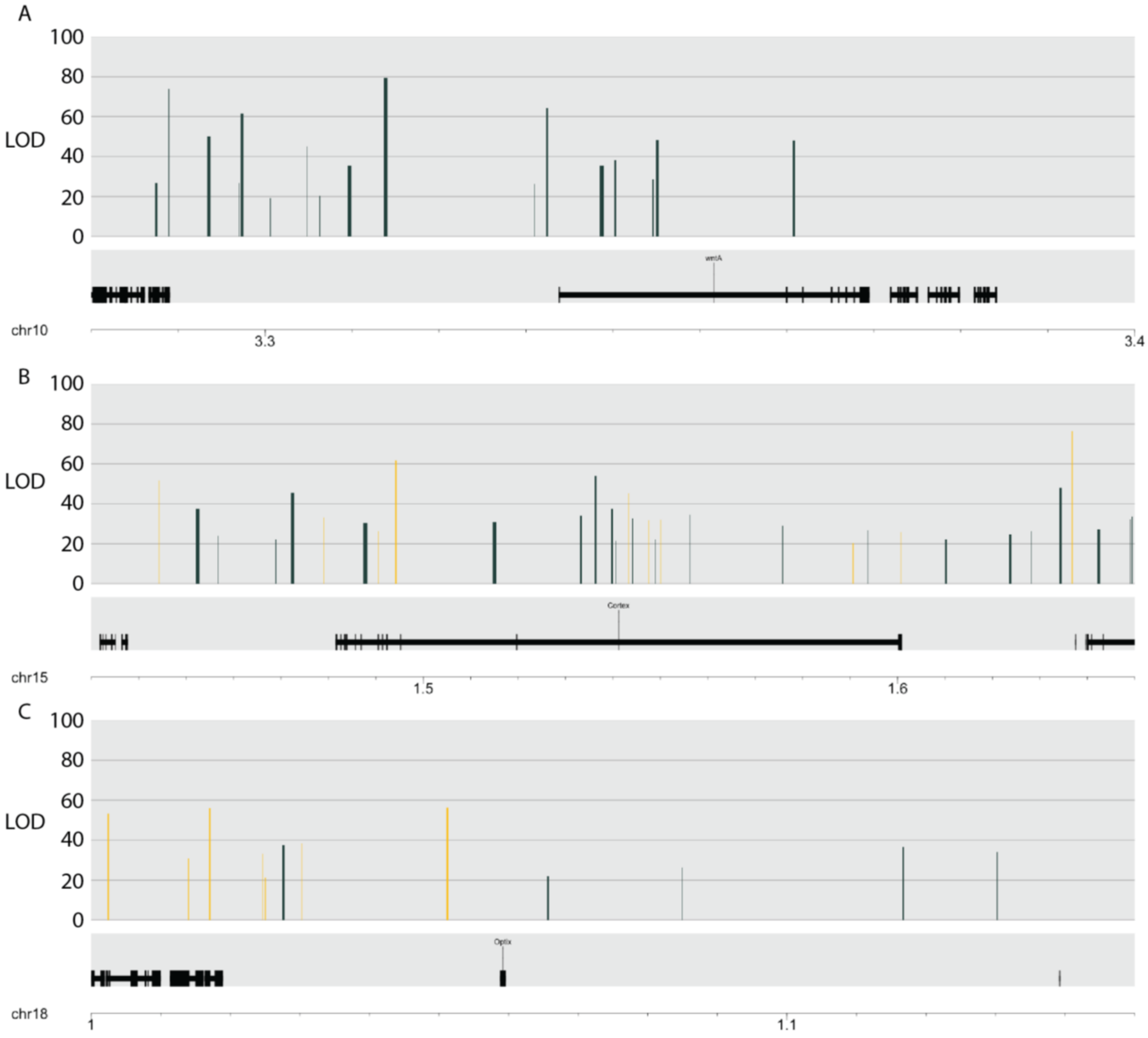

**Figure S7.4: *Heliconius*-specific evolution around known color pattern loci**
Ticks are *Heliconius* chromosomal coordinates, with scaffold order corresponding to Hmel2.5. Conserved regions are shown in yellow, while accelerated regions are shown in green. All chromosomal conservation maps are available on Dryad.

**Section 8: Topology distribution**
*return to top*

We calculated regional topologies in sliding windows across the genome in a number of different configurations. Interestingly, although we have a single multi-genome alignment, the fraction of windows that recovered each topology varied depending on whether the *H. erato demophoon* v1 or Hmel2.5 coordinates were used as reference. This may be due to weaker alignability between the *erato-sara* clade genomes and the Hmel2.5 assembly relative to *H. erato demophoon*, or may be due to the fact that the *H. erato demophoon* genome is approximately 100MB (33%) larger than the Hmel2.5 genome. The genome topology map using the Hmel2.5 coordinates is shown in Figure S8.1 A, and the most common topologies, their relative fractions, and block length distributions are shown in Figure S8.1 B, C. We also used 10KB non-overlapping windows instead of 50KB when mapping onto the *H. erato* genome. Those data are shown in Figure S8.2 and were used for all subsequent analyses of topology distributions.

We next aimed to elucidate structural genomic features associated with regional evolutionary histories. As stated in the main text, the fraction of a given topology recovered in each chromosome was strongly correlated with that chromosome's total size in base pairs, and relationships for all 8 of the most common topologies are shown in Figure S8.3, and their correlation coefficients are listed in Table S8.1. The Z chromosome was excluded when computing correlation coefficients, as the values recovered from it are clear outliers, almost certainly because the Z chromosome is often involved in F1 hybrid female sterility, in accordance with Haldane's Rule[22,23]. Values from the chromosomes that contain inversions (chr2 and chr15) are also outliers, but as they form a minority of each chromosome, we included their chromosomes in this analysis; thus reported values are conservative estimates. Tree labels in Table S8.1 correspond to those in the main text Figure 2B. All 8 most common topologies had significant associations with chromosome size ($p<.05$), with correlation coefficients ($r^2$) ranging from 0.197 to 0.883. We also examined the correlation between topology and binned position along the chromosome – Figure S8.4 reflects the per-bin breakdown (all topologies within a bin sum to 1), while Figure S8.5 reflects the per-tree breakdown (all measures of each tree type sum to 1). Regardless of how we split the data, we find an increase in "species tree" topology in the low-recombination regions at the edges of chromosomes. We then binned windows by their recombination rate as determined by linkage disequilibrium[4], and again saw a very strong negative correlation between species tree topology and recombination rate. Finally, we also predicted that topology would be associated with the density of coding sequence per window if incompatibility alleles were more likely to be in coding sequence. That relationship is much weaker, but we detect a minor decrease in species tree topologies in regions with very low coding sequence density(Figure S8.7).

Although we have evaluated the effect on topology of chromosome size, chromosomal position, local recombination rate, and coding density separately, these measures are not independent. Recombination rate is higher in regions of low gene density (Figures S8.8, S8.9), and is lower at the edge of chromosomes (Figure S8.10). However, gene density is constant across chromosomal position (Figure S8.11). There is an unexpected, slightly positive relationship between chromosome size and gene density (Figure S8.12), and a strongly negative relationship between chromosome size and average recombination rate (Figure S8.13), though this correlation is actually weaker than expected based on data from *H. melpomene*[24,25], though the sample size for *H. erato* is much lower than that for *H. melpomene.*

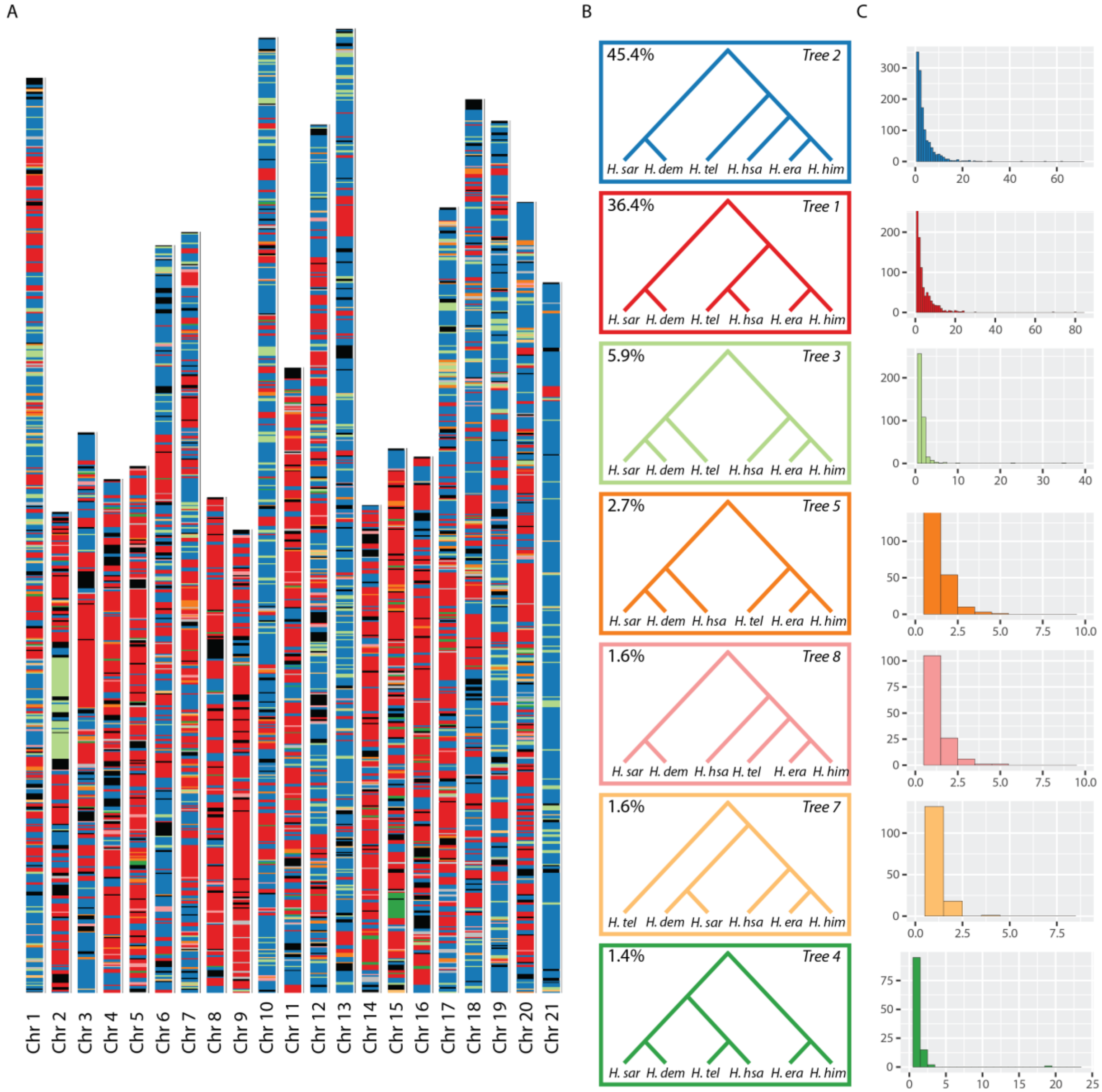

**Figure S8.1: Heterogeneity of *erato-sara* clade evolutionary history across the *melpomene* genome**

***A. Distribution of topologies across the genome.*** *For each 50kb window, the colored bar represents the topology recovered from that region. Colors correspond to topologies in* ***B.*** *Coordinates are in terms of the H. melpomene Hmel2.5 reference, and topologies for all windows were reconstructed with PhyML. Black regions are missing data.* ***B. Common***

***topologies.*** *The eight most common phylogenies are shown. The value in the top left corner of each topology is the percentage of all 50kb windows that recovers that topology. The tree labels correspond to those in the main text Figure 2B* ***C. Tree block length distribution*** *Each histogram corresponds to the topology of the same color in* ***B****, and shows the distribution of the number of consecutive 50kb windows that recover that topology.*

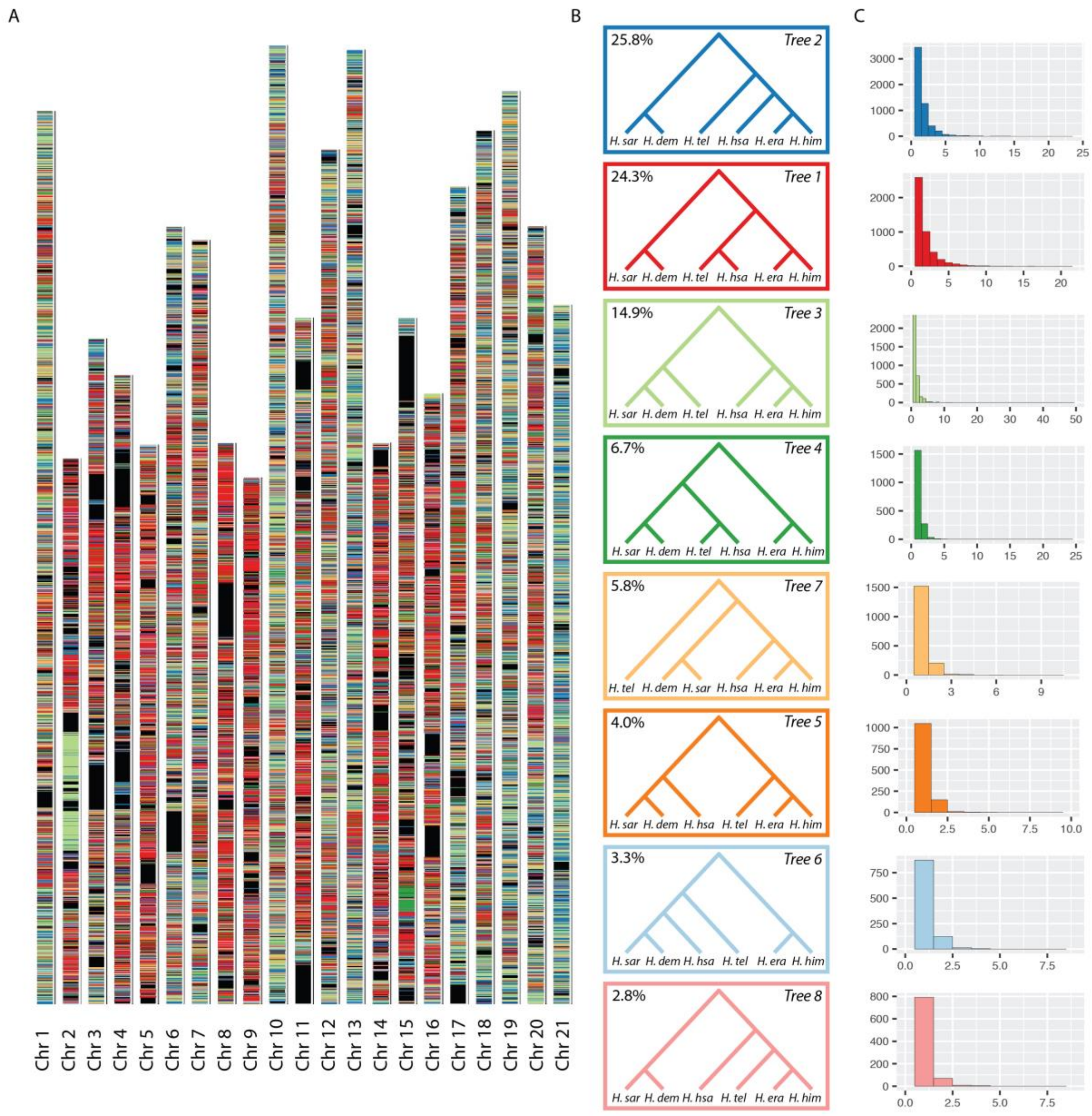

**Figure S8.2: Heterogeneity of evolutionary history across the *erato* genome, 10KB windows**
***A. Distribution of topologies across the genome.*** *For each 10kb window, the colored bar represents the topology recovered from that region. Colors correspond to topologies in* ***B.*** *Coordinates are in terms of the H. erato demophoon v1 reference, and topologies for all windows were constructed with PhyML. Black regions are missing data.* ***B. Common topologies.*** *The eight most common phylogenies are shown. The value in the top left corner of each*

*phylotopology is the percentage of all 10kb windows that recovers that topology. The tree labels correspond to those in the main text Figure 2B.* ***C. Tree block length distribution*** *Each histogram corresponds to the topology of the same color in* ***B****, and shows the distribution of the number of consecutive 10kb windows that recover that topology.*

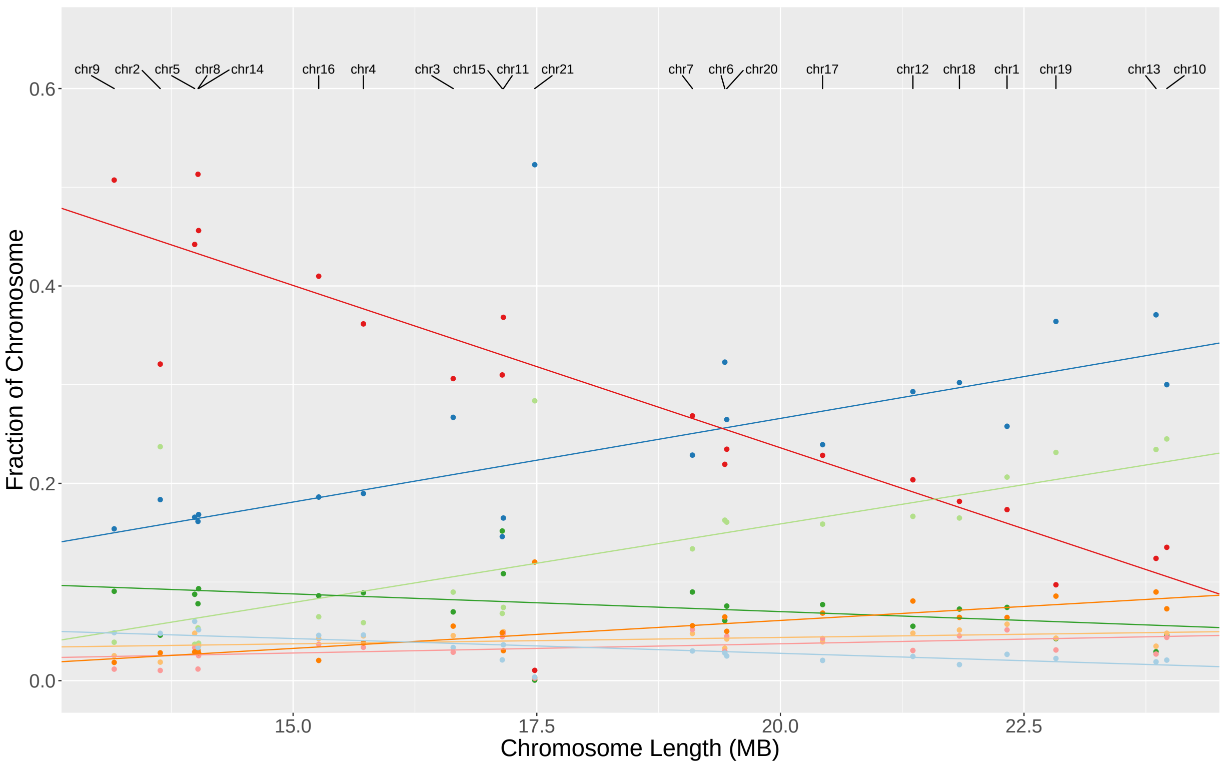

**Figure S8.3: Relationship of topology to chromosome length**

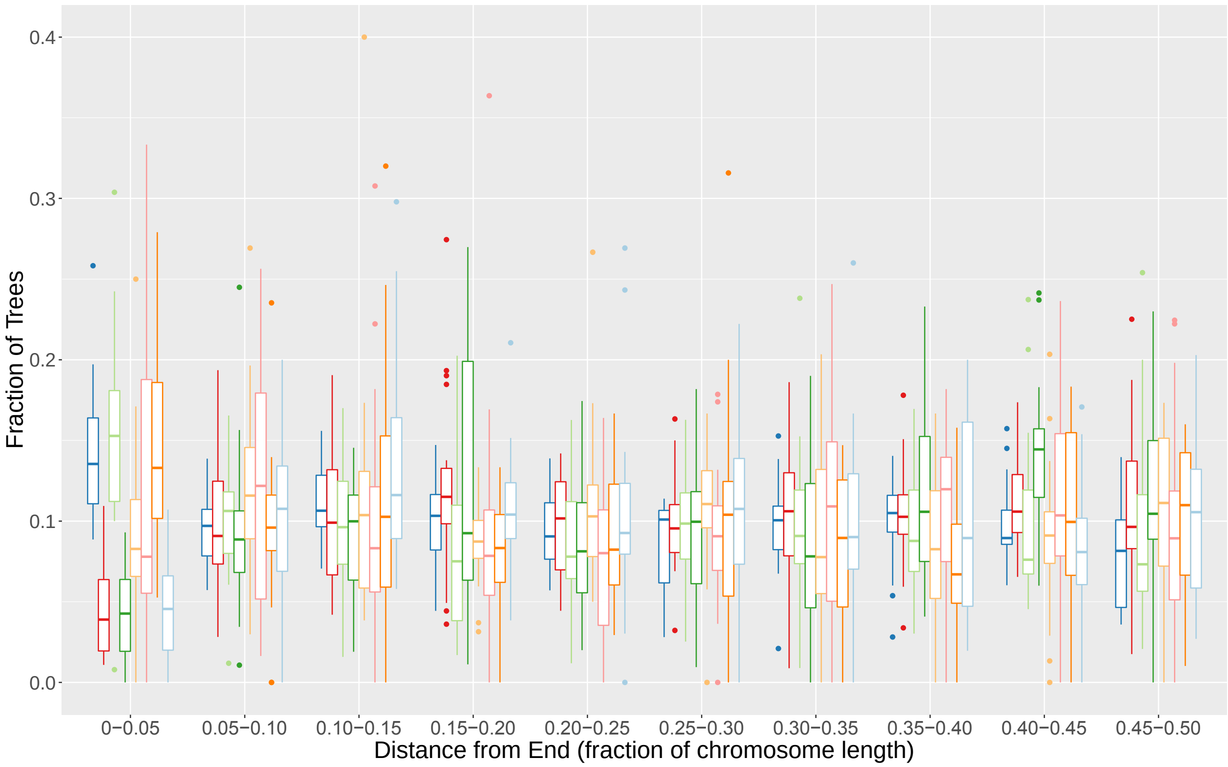

**Figure S8.4: Relationship of topology to chromosomal position, per tree**

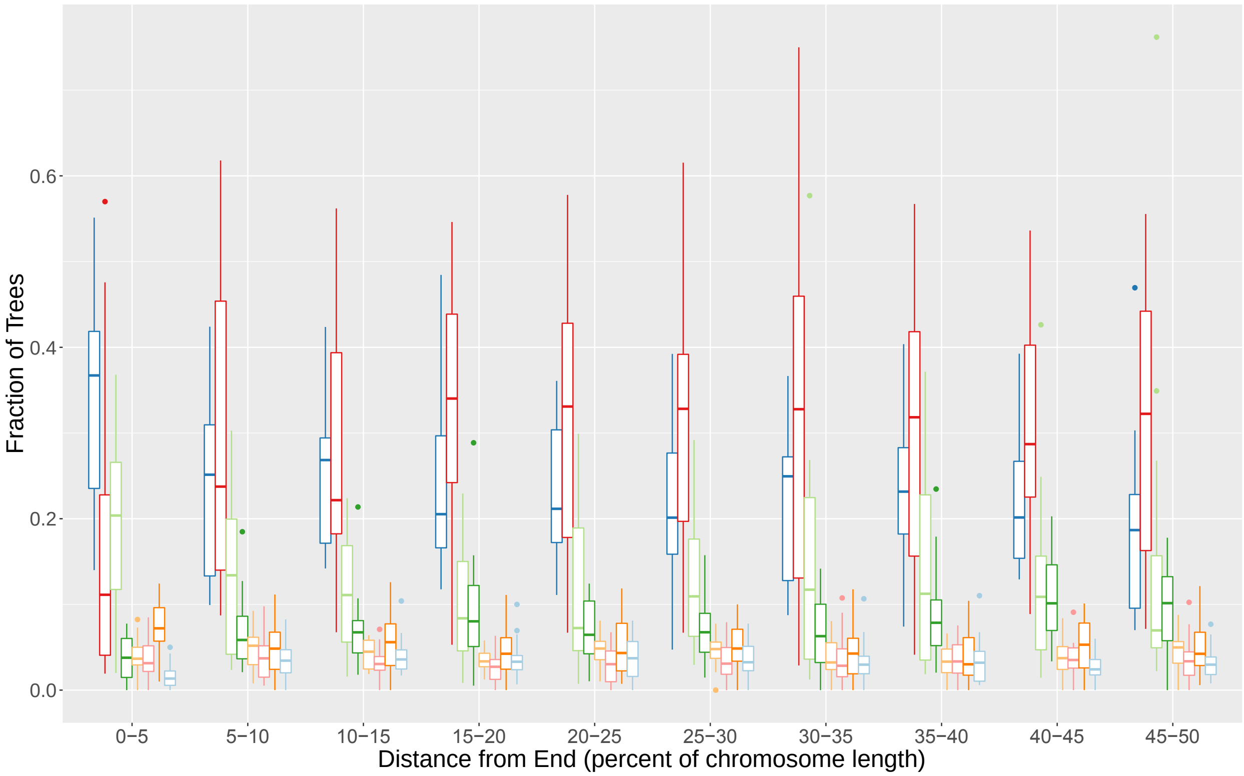

**Figure S8.5: Relationship of topology to chromosomal position, per position bin**

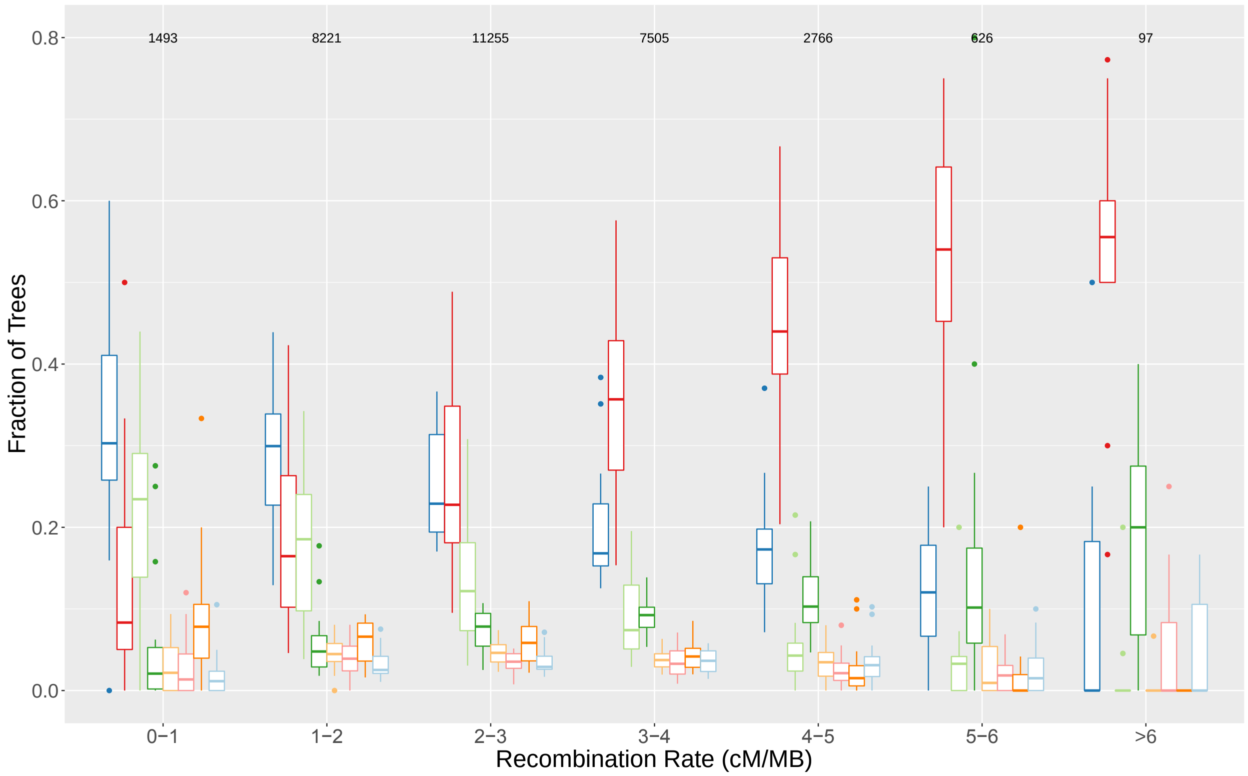

**Figure S8.6: Relationship of topology to recombination rate**

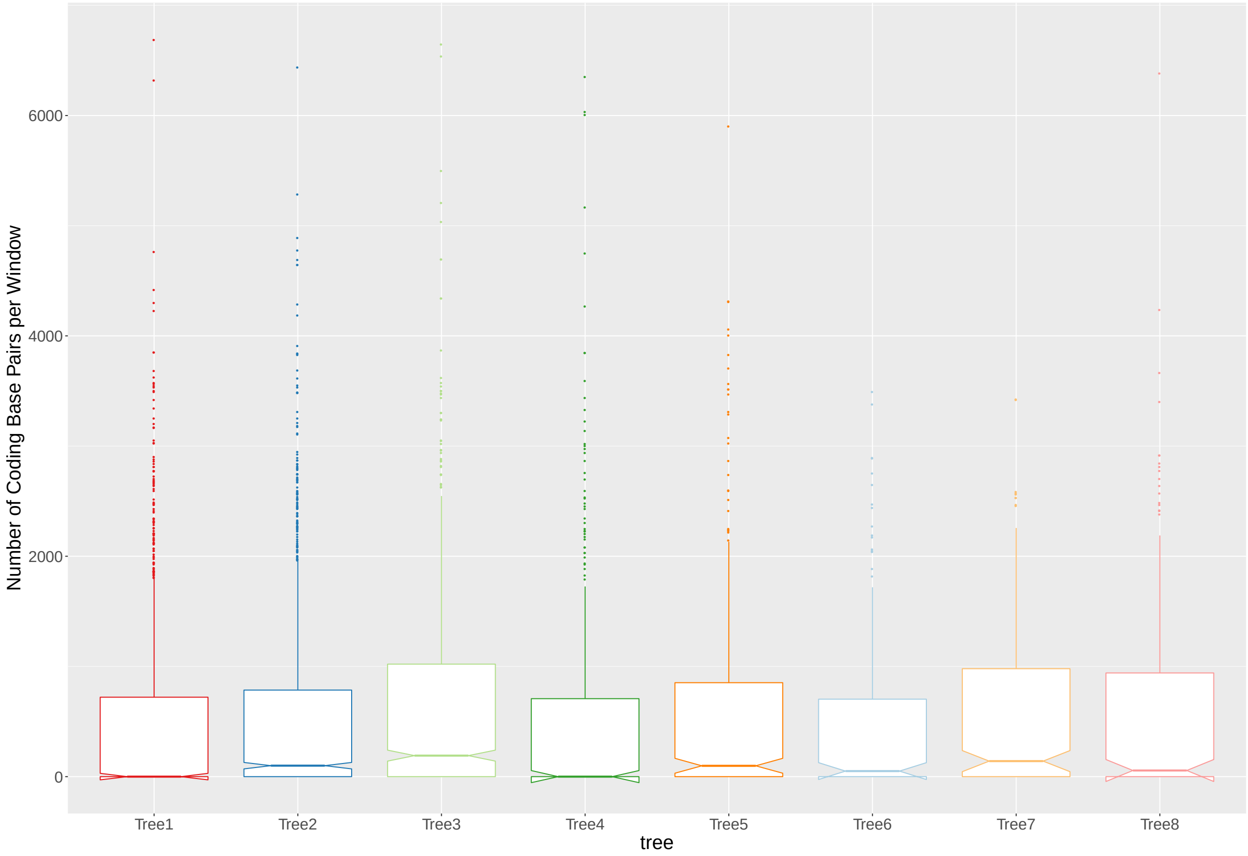

**Figure S8.7: Relationship of topology to number of coding base pairs per window**
Central lines show median number of coding bases per window that recovers given topology. Box edges correspond to the inter-quartile range (IQR), and the minimum value they can take is 0. The notches represent $1.58 * \frac{\mathrm{IQR}}{\sqrt{n}}$ , and can therefore take values less than zero{Wickham:2016tn} as seen in Trees 1, 4, 6, and 8. Tree labels correspond to those in main text Figure 2B.

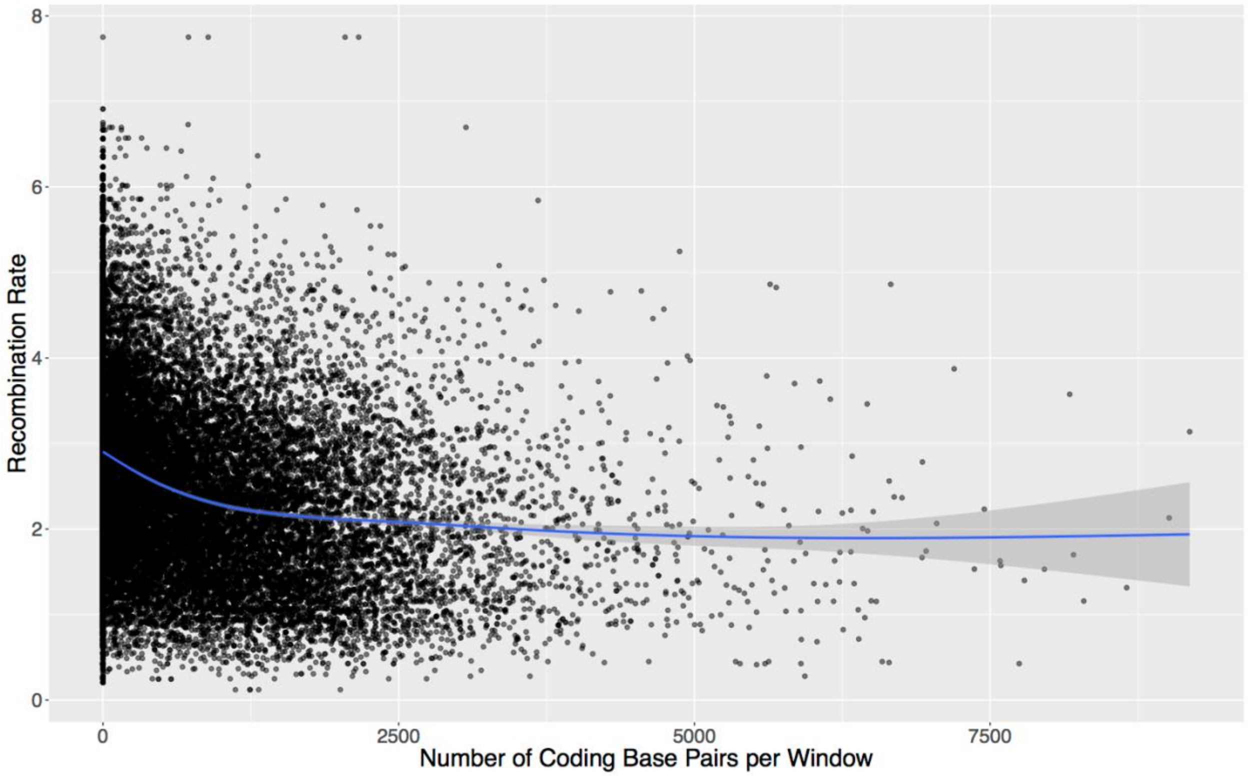

**Figure S8.8: Relationship of recombination rate to number of coding base pairs per window**

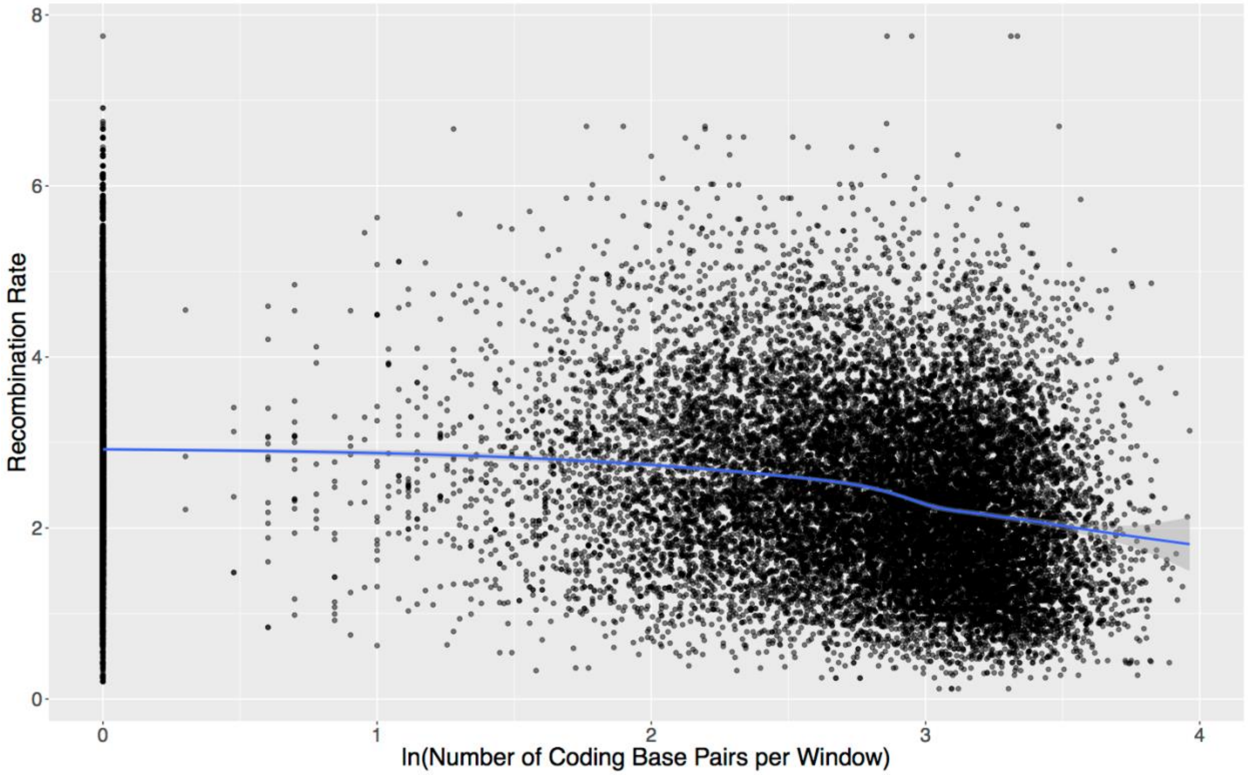

**Figure S8.9: Relationship of recombination rate to log number of coding base pairs per window**

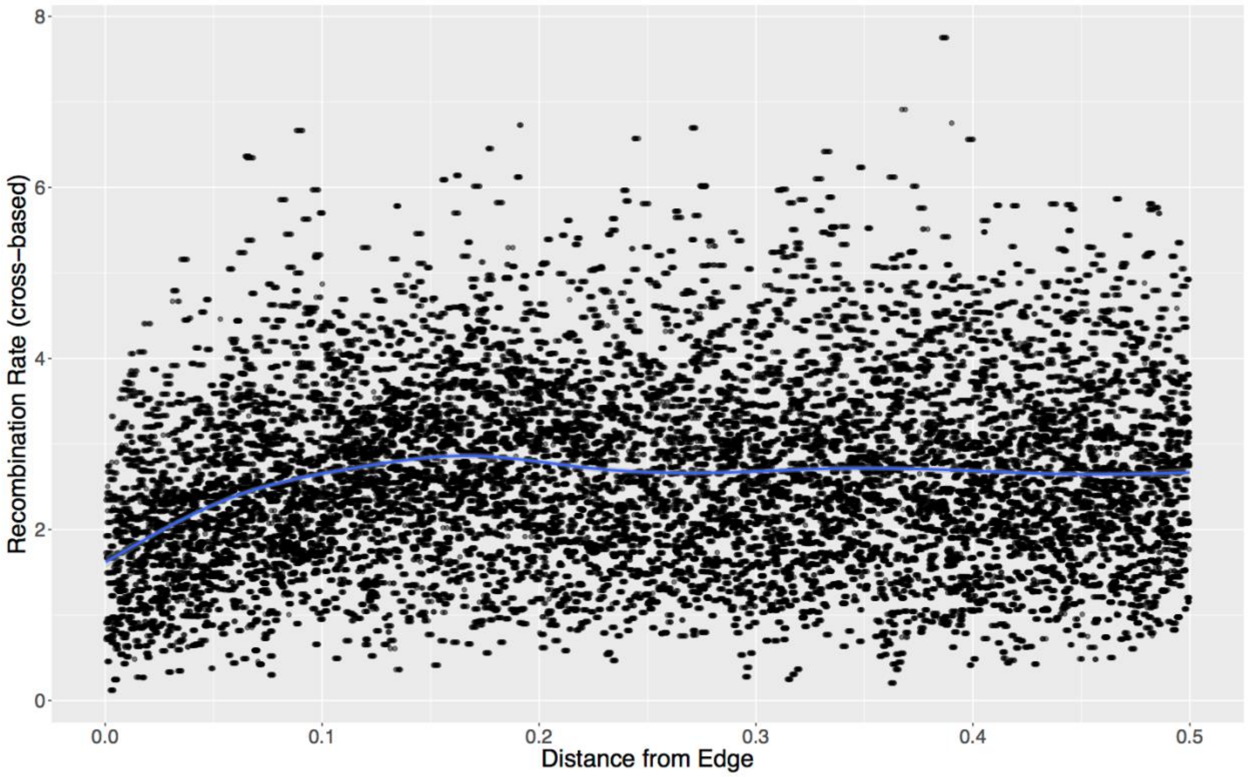

**Figure S8.10: Relationship of chromosomal position to recombination rate**
Local recombination rates for the H. erato demophoon genome were calculated in 50KB windows. Because topologies in this study were reconstructed in 10KB windows, adjacent windows often share the same recombination rate estimate.

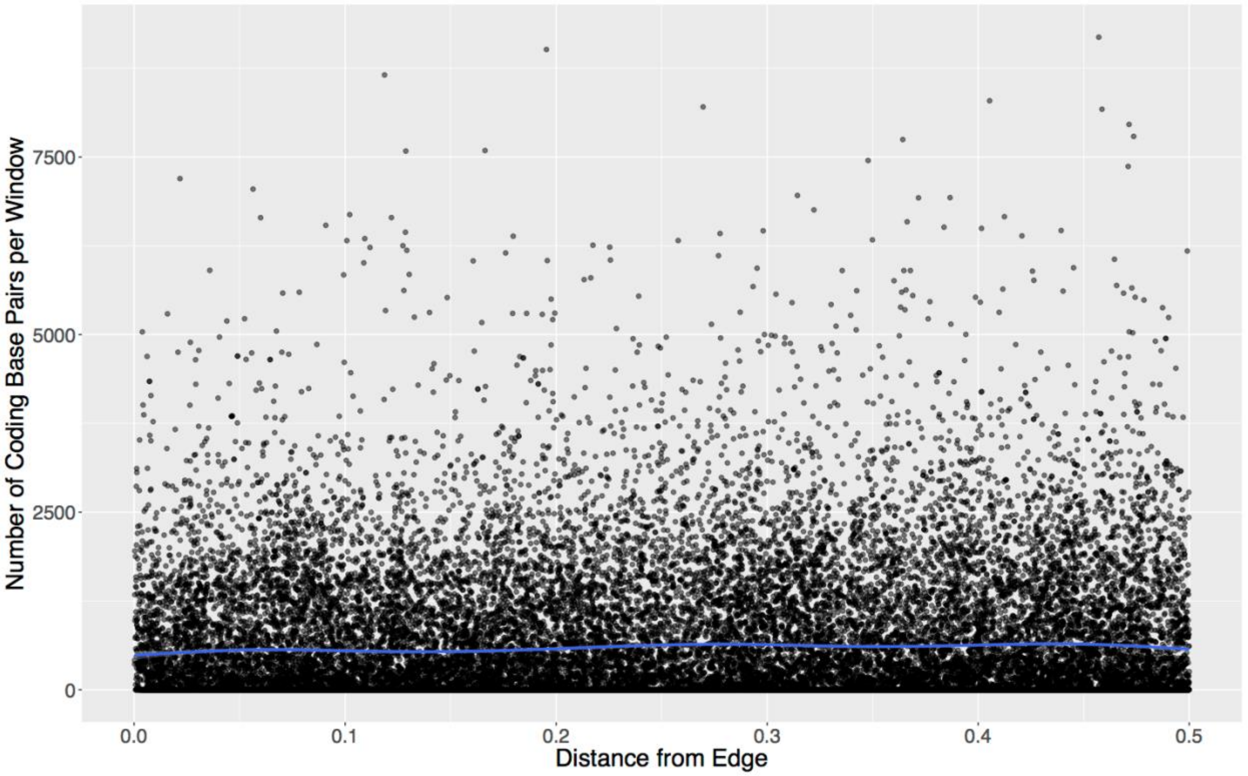

**Figure S8.11: Relationship of chromosomal position to number of coding bases per window**

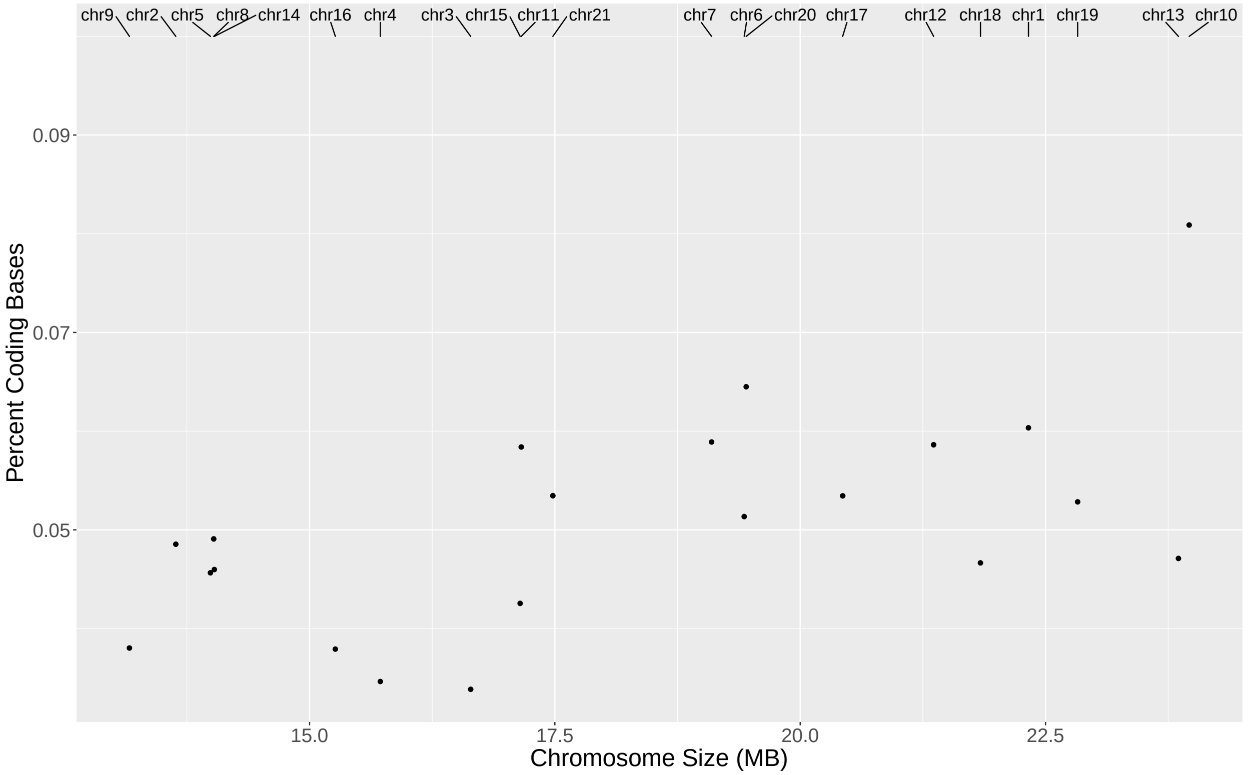

**Figure S8.12: Relationship of chromosome size to fraction of coding bases**

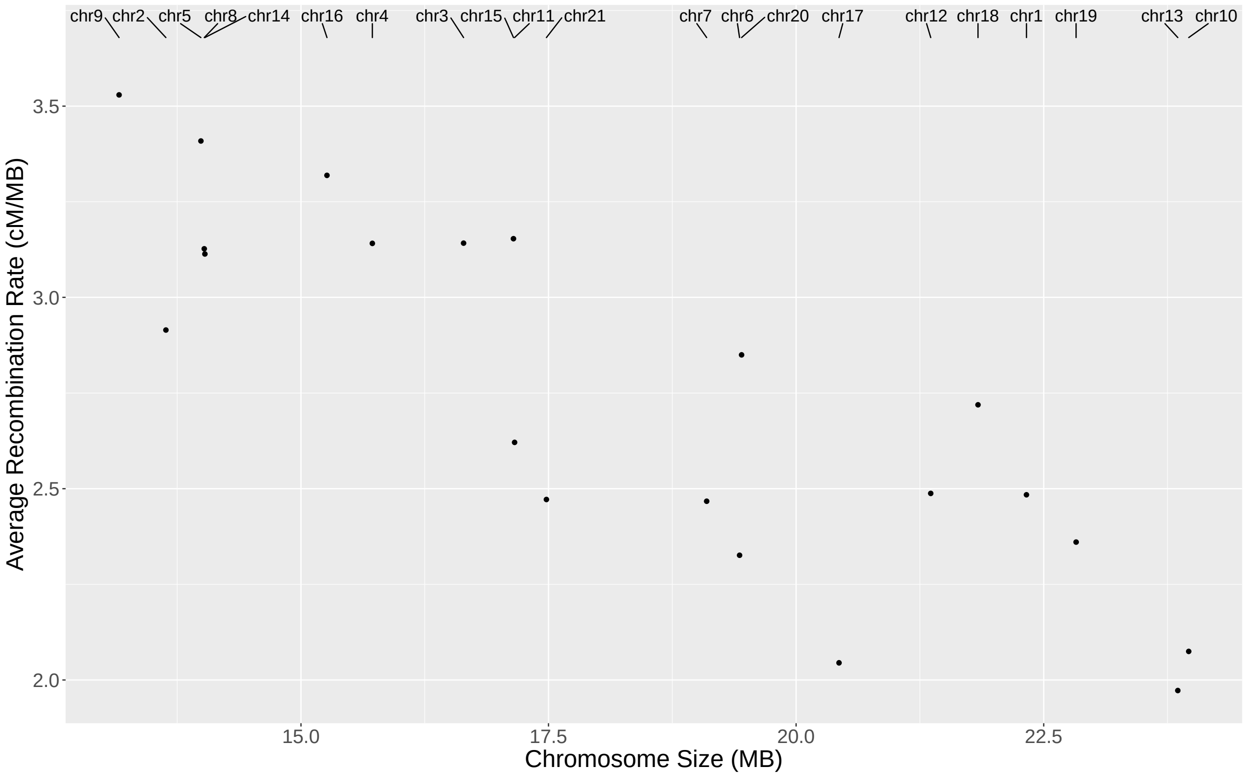

**Figure S8.13: Relationship of chromosome size to average recombination rate**

Interestingly, some evidence arising from this recombination-based method suggests that the species tree presented in the main text is not entirely correct. When examining the topology restricted to the triplet *H. demeter – H. telesiphe – H. erato*, windows with the "species tree topology", (*H. demeter*, *(H. telesiphe*, *H. erato*)) are slightly negatively correlated with chromosome size, while windows with the "introgression topology" (*H. erato*, *(H. demeter*, *H. telesiphe*)) show a slight positive correlation (Figure S8.14). A small subset of our species trees also support this topology, but the relationships are weak and require further study before any strong case can be made to overturn the prevailing evidence of ASTRAL and concatenation species tree methods.

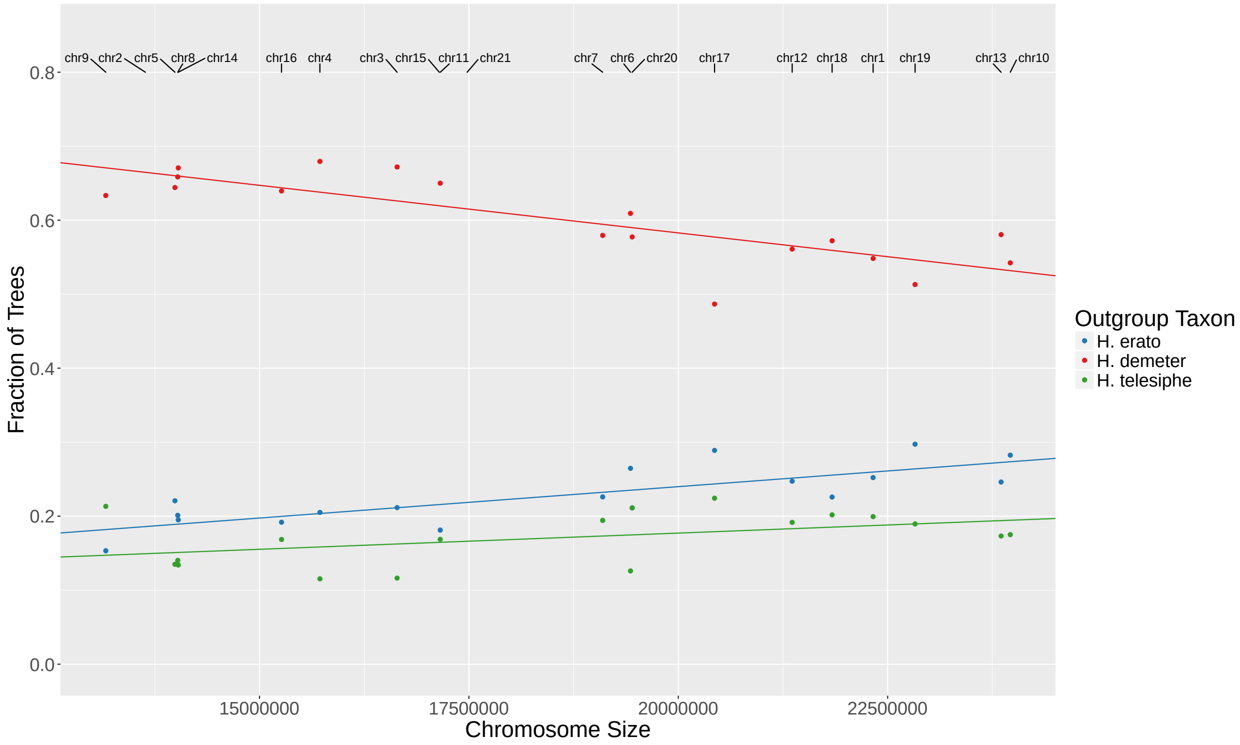

**Figure S8.14: Evidence for *H. telesiphe* as sister to *H. demeter* and *H. sara***

### Section 9: Triplet Internal Branch Length Test

#### 1 Introduction

Identifying the genetic signatures of introgression is of interest due to its impact on patterns of adaptation and speciation. However, random features such as deep coalescence (also called incomplete lineage sorting, or ILS) can muddy the signal of introgression by generating gene trees which differ from the species tree without any interspecies gene flow. One robust genome-wide approach to detecting introgression is Patterson's $D$ statistic, which looks at imbalances in the frequency of discordant gene tree topologies [27] . This statistic and related $F$ statistics can be motivated from a coalescent standpoint [28], and can provide genome-wide estimates of introgression. We wish to extend these tests to locate introgressed loci or estimate the likelihood that a given region displays its gene tree topology due to introgression rather than ILS [29] through use of additional information in gene tree branch lengths.

We present a statistical framework for assessing whether a given set of gene trees shows evidence for introgression events or could be generated by ILS alone, as well as for calculating the odds that a given discordant locus was generated by ILS or introgression. These calculations are unwieldy on the full gene tree, so inspired by the approach of Zairis *et al.* [30], we focus on statistics of the set of three-leaf gene subtrees for each gene genealogy in our sample. Rather than consider every relationship in our sample at a particular locus, we take all sets of three terminals in the original tree and examine these triplet trees independently of each other. In particular, we compare the internal branch length of a given triplet at a locus to the genome-wide distribution to classify it as likely introgressed or not. We rely on the following notation.

| | |
|---|---|
| $N_e$ | Effective population size, assumed to be constant through time. |
| $u$ | Per-site, per-generation mutation rate. |
| $\theta$ | $4N_eu$, the population mutation parameter. |
| $EXP(\lambda)$ | Exponential distribution with scale parameter $\lambda$. |
| $TEXP(\lambda, C)$ | Truncated exponential distribution with scale parameter $\lambda$ and truncation point $C$. |
| $S_i$ | Time interval during which the species tree has exactly $i$ extant lineages, measured in units of $2N_e$ generations. |
| $H$ | Time back from the present until a hybridization event, measured in units of $2N_e$ generations. |

Table 1

#### 2 Intuition

We model the distribution of the length of the internal branch of a gene tree as a mixture of distributions resulting from the coalescence of two lineages within a branch of the species tree which only they share (e.g. introgression or speciation) and coalescence within the common ancestral population (e.g. ILS). Consider the simple case of a rooted species tree with three terminals. We can use coalescent theory to derive the expected branch length distribution of gene trees generated by that species tree. ILS alone will generate all three topologies with equal frequency and expected branch lengths, since the three lineages are exchangeable once they have reached the ancestral population. In addition to this, the gene tree topology consistent with the species tree is realized when the two sister lineages coalesce within the internal branch of the species tree, and as a result this matching gene tree topology will be both enriched in frequency and contain longer internal branches (Figure 1).

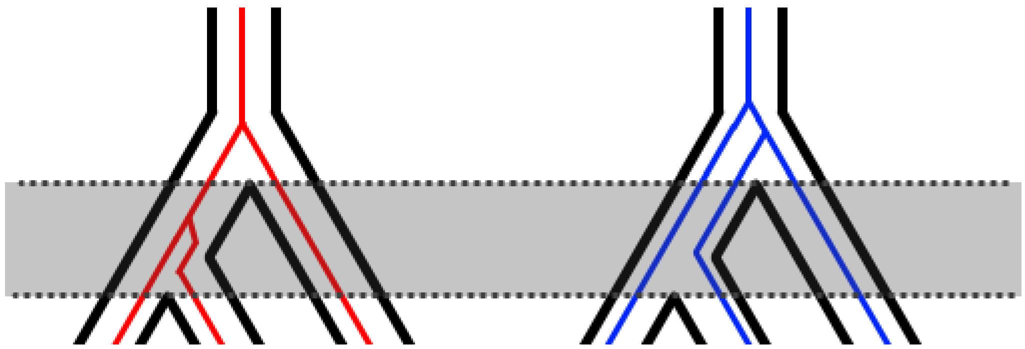

Figure 1: In gene trees matching the species tree topology (left, red), there is an additional window of time in between speciation events (highlighted in grey) where coalescence can occur. However, for non-matching topologies (right, blue) coalescence can only occur in the MRCA.

We condition on the gene tree topology, and focus on the information present in its internal branch. Gene trees generated by ILS will have an internal branch length that follows an exponential distribution with scale parameter equal to one when time is measured in units of $2N_e$ generations.

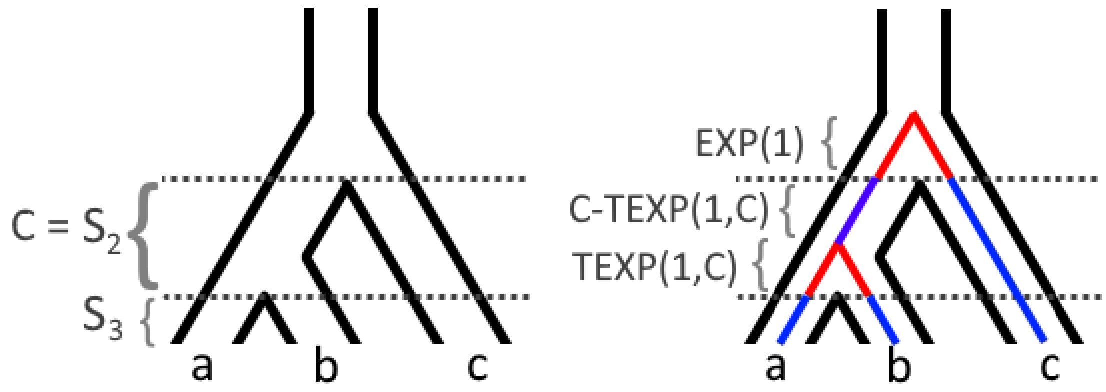

(a) One sample tree for three species, with the relevant times between speciation events labeled.

(b) Given a sample gene tree, we can split the branches into components based on their generating distribution.

Figure 2

If there is no introgression, all discordant gene tree topologies are generated by ILS alone, and will have internal branches sampled from an $EXP(1)$ distribution. In contrast, the concordant gene tree topology will have internal branches pulled from the distribution $EXP(1) + S_2 - TEXP(1, S_2)$, as shown in figure 2. Further, as shown in figure 3, if introgression has occurred at a locus, we can take the corresponding set of paths in the species network and obtain another distribution of the form $EXP(1) + C - TEXP(1, C)$, see figure 3.

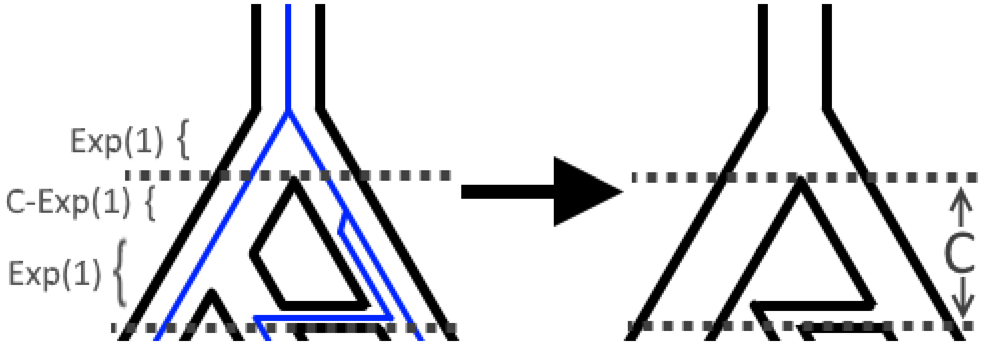

Figure 3: If a gene tree follows an introgression path in the species graph, the species history that particular locus experiences can still be described completely as a tree. As a result, the internal branch distribution is of the form $EXP(1) + C - TEXP(1, C)$. In this case, $C$ is the time from the introgression event back to the common ancestor of all three species. Using the notation in table 1, we have $C = S_2 + S_3 - H$ for this example.

The full branch length distribution for a given gene tree topology will be a mixture generated by multiple of these phenomena. Our goal is to use the observed branch length distributions to infer the mixture proportions of gene trees coming from each possible source.

### 3 The Mixture Distribution of Internal Branches of Triplets

So far we have described the internal branch length of a triplet gene tree as $EXP(1) + C - TEXP(1, C)$ in coalescent units. In reality, branch lengths must be inferred from genetic data, in units proportional to mutations rather than $N_e$. We assume that they are measured in units of substitutions per site. If we take mutations to be Poisson distributed along the branches of the genealogy we require a scaling factor of $\lambda = \theta/2$. This gives us the following:

$$f(x; C, \lambda) = \begin{cases} \frac{1}{2\lambda}(1 + e^{C})e^{-x/\lambda} & C\lambda \leq x \\ \frac{e^{-x/\lambda} - e^{x/\lambda}}{2\lambda(1-e^{c})} & C\lambda > x \\ 0 & x < 0 \end{cases}$$

Notably, this is non-differentiable at $C\lambda$, where it achieves its maximum, and reduces to an exponential distribution when $C = 0$, corresponding to the lineages being sampled from a single population. For any given gene tree triplet topology, the internal branch length distribution will be a mixture of distributions of this type (Figure 4). There will always be some ILS component that has $C = 0$, with a possible additional distribution with $C$ corresponding to the amount of time during which the two sister gene lineages are isolated. In reality, the mixture proportions between these distributions will be determined by the effective population sizes and lengths of each branch in the species tree. However, our implementation of expectation maximization to infer these parameters treats them as independent for computational simplicity, as it allows us to estimate the mixture proportions and parameters of each distribution iteratively (explained in greater detail in section 4.1). We find that our method still performs well despite this relaxation (see section 4.3).

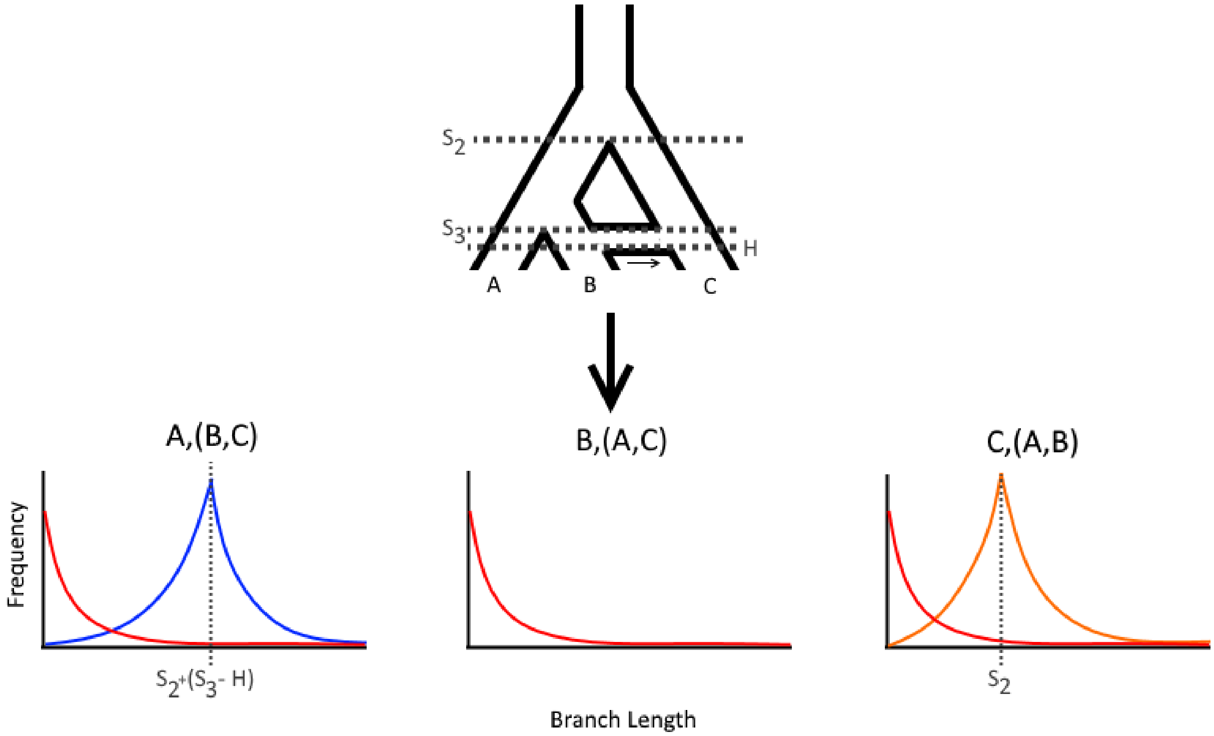

Figure 4: A species triplet with an introgression event indicated by the black arrow showing the direction of gene flow backwards in time generates a characteristic branch length distribution. For the topology generated by introgression, $A,(B,C)$, a fraction of gene trees result from introgression (with distribution shown in blue) with $C = S_2 + (S_3 - H)$. Similarly for the concordant gene tree topology, a portion of gene trees are generated by speciation with $C = S_2$ (shown in orange). In all three topologies, ILS generates trees following an exponential distribution (red).

### 4 Inference

Since we need to estimate both the assignment of each datapoint to a distribution as well as the parameters of the distributions themselves, we use an iterative inference algorithm called expectation maximization (EM), which we will describe using notation from table 2. Given a number of distributions to fit, $K$, EM works by alternating between two problems. First is the E-step, where we calculate the relative probability that a given value could be sampled from either of the distributions given values for the parameters $\{C_k\lambda\}$, $\lambda$, and$\{\pi_k\}$ (the peak locations, scaling, and mixing proportions respectively). Next is the M-step, where we fix the probabilities calculated in the E-step and try to find the values of $\{C_k\}$, $\lambda$, and $\{\pi_k\}$ that maximize the log-likelihood.
As we need a fixed $K$, we will define two competing models for each gene tree triplet topology, one with initial parameters $K = 1, C_1 = 0$ (an ILS model) and another with $K = 2, C_1 = 0, C_2 = 1$ (a model with some gene tree topology bias, whether due to speciation or introgression).

#### 4.1 Expectation Maximization

We use EM to infer the parameters of the mixture models (see algorithm 1). For the initial values, we take the mixture proportion of the $k^{th}$ distribution, $\pi_k$ to be $1/K$, $C_0 = 0$, and $C_1 = 1$. Given $\{C_k\}$, the optimal assignments of data to the $k^{th}$ distribution is the ratio $\pi_k f(x; C, \lambda)/\Sigma_k^K \pi_k f(x; C, \lambda)$, which we denote $q_{C_k\lambda}(x_i)$. We update this value in the E-step and split the M-step into two parts, updating first the the peak of the $k^{th}$ distribution $(C_k\lambda)$ followed by the scalar $\lambda$ conditioned on $C_k\lambda$. This repeats until the change in log-likelihood is below a specified precision threshold.

| | |
|---|---|
| $X$ | The data, in the form of internal branch lengths in units of substitutions per site, with length $n$. |
| $x_i$ | The $i^{th}$ branch length in $X$. |
| $K$ | The total number of contributing distributions in the observed triplet branch length mixture distribution, either 1 or 2 in our implementation. |
| $\lambda$ | Scaling factor for converting our observed distributions to coalescent units ($\frac{1}{2N_e\mu}$). |
| $C_k$ | Interval parameter $C$ of the $k^{th}$ distribution. |
| $\pi_k$ | Mixing proportion of the $k^{th}$ distribution. |
| $q_{C\lambda}(x_i)$ | $\pi_i f(x;C,\lambda)/\Sigma_k^K \pi_k f(x;C,\lambda)$, the soft assignment of the value $x_i$ to either of the two $K$ in $\{0,1\}$. |
| $\log(L(\{x_i\},\{C_k\},\lambda))$ | The log-likelihood, $\log \prod_{x_i\in X} \sum_{C_k} \pi_k f(x_i;C,\lambda)$. |

Table 2

**Algorithm:** Triplet Topology EM

**Input:** $X$ the set of branch lengths for a given gene tree triplet topology.

$K \leftarrow 2$
$C_1 \leftarrow 0$
$C_2 \leftarrow 1$
$\pi_k \leftarrow 1/K$
$\lambda \leftarrow \bar{x_i}$
$L_1, L_2 \leftarrow 0, |L(\{x_i\},\{C_k\},\lambda)|$
**while** $L_2 - L_1 > 10^{-6} * L_1$ **do**
    $q_{C\lambda}(x_i) \leftarrow \pi_k f(x;C,\lambda)/\Sigma_k^K \pi_k f(x;C,\lambda)$
    $C_2\lambda \leftarrow \text{argmax}_{C\lambda} \Sigma_{i=1}^n q_{C_k}(x_i) ln(f(x;C,\lambda))$ ▷ `Find` $C\lambda|\lambda$`, detailed in 4.1.1`
    $\lambda \leftarrow \text{argmax}_{\lambda} \Sigma_{i=1}^n q_{C_k}(x_i) ln(f(x;C,\lambda))$ ▷ `Find` $\lambda|C\lambda$`, detailed in 4.1.2`
    $\pi_k \leftarrow mean_{x_i}(q_{C_k}\lambda(x_i))$ ▷ `Update proportions to the mean of the soft assignments`
    $L_1 \leftarrow L_2$
    $L_2 \leftarrow |L(\{x_i\},\{C_k\},\lambda)|$
**end**
**return** $C_2, \lambda, \{\pi_k\}$

**Algorithm 1:** Our expectation maximization implementation for inferring the mixture components of a triplet gene tree branch length distributions.

##### 4.1.1 Picking $C\lambda$

In the M-step we want to maximize the log-likelihood by solving the following for each $k$:

$$C_k\lambda = \text{argmax}_{C,\lambda}\Sigma_{i=1}^{n} q_{C_k}(x_i) ln(f(x;C,\lambda))$$

However, since $f$ is not differentiable at $C\lambda$ we need to consider points on either side of each $C\lambda$ separately. We rewrite $\Sigma_{i=1}^{n} q_{C_k}(x_i)\log(f(x;C,\lambda))$ in terms of the data above and below $C\lambda$:

$$\begin{aligned}
\text{E}[\log(L)] &= \sum_{x_i<C\lambda} q_C(x_i)\log\left(\frac{-e^{\frac{x_i}{\lambda}}+e^{\frac{x_i}{\lambda}}}{2\lambda(-1+e^C)}\right) + \sum_{x_i\geq C\lambda} q_C(x_i)\Bigg(-\log(2)+\log\left(\frac{e^{\frac{-x_i}{\lambda}}(1+e^C)}{2\lambda}\right)\Bigg) \\
&= \sum_{x_i<C\lambda} q_C(x_i)(\log(\sinh(x_i/\lambda))) - \sum_{x_i>C\lambda} q_C \log(e^{\frac{-x_i}{\lambda}}/2\lambda) + \sum_{x_i>C\lambda} q_C(-\log(2)) \\
&+ \sum_{x_i>C\lambda} q_C(x_i)(\log(1+e^C)) - \sum_{x_i<C\lambda} q_C(x_i)(\log(e^C-1))
\end{aligned}$$

Conditioned on a particular value of $\lambda$, only the two terms on the last line differ between values of $C\lambda$ equal to sequential elements of the data. Furthermore, they are both convex, and consequently the whole sum is convex for values of $C\lambda$ between adjacent values in $X$. As a result, the expected likelihood is maximized by $C\lambda$ taking on a value realized by the branch lengths in the data. We need only search these finite possible values for our optimal $C\lambda$ and return it.

##### 4.1.2 Finding the Conditional $\lambda$ Using Gradient Ascent

To find $\lambda$ given $C_k\lambda$, we calculate the $\lambda$-derivative of $\log(L)$ at the given value for $\lambda$ and change $\lambda$ by an amount proportional to this quantity. In the space of mixture models we are considering, we only need this derivative for $k=2$, as when $k=1, C_1=0$ we only have an exponential and the maximum likelihood estimate of $\lambda$ is $\bar{x_i}$. For the former case, we still take $C_1=0$ as the exponential ILS component. This gives us the following for $k=2$:

$$\begin{aligned}
\log(L) &= \log\prod_{x_i\in X}\sum_{C_k}\pi_k * f(x_i;C_k,\lambda) \\
\frac{\partial\log(L)}{\partial\lambda} &= \partial\lambda\sum_{x_i\in X}\log\sum_{C_k}\pi_k * f(x_i;C_k,\lambda) \\
&= \sum_{x_i\in X}\partial\lambda\log(\pi_1 * f(x_i;C_1,\lambda)+\pi_2 * f(x_i;C_2,\lambda)) \\
&= \begin{cases} \dfrac{-C_2\lambda-\lambda+\frac{C_2\lambda(2-\pi_2)}{\pi_2 e^{C_2\lambda}+2-\pi_2}+x}{\lambda^2} & C_2\lambda\leq x \\[2ex] \dfrac{\frac{C_2\lambda}{e^{C_2\lambda}-1}-\lambda-\frac{C_2\lambda(2-\pi_2)+\pi_2(2x-C_2\lambda)e^{\frac{2x}{\lambda}}}{2(1-\pi_2)\left(e^{C_2\lambda}-1\right)+\pi_2\left(e^{\frac{2x}{\lambda}}-1\right)}+x}{\lambda^2} & C_2\lambda > x \end{cases}
\end{aligned}$$

We note that this can be extended to $K>2$, but we do not observe distributions with more than two modes in our dataset. For completeness, we include the $k=3$ case below. We use the following symbols for simplicity: $C_2\lambda=\beta$, $C_3\lambda=\gamma$, $\pi_2=p$, and $\pi_3=q$.

$$\begin{cases}
\dfrac{x\cosh\left(\frac{x}{\lambda}\right)\left(-\frac{p}{e^{\beta/\lambda}-1}-\frac{q}{e^{\gamma/\lambda}-1}\right)+\sinh\left(\frac{x}{\lambda}\right)\left(\frac{p\left(e^{\beta/\lambda}(\beta-\lambda)+\lambda\right)}{\left(e^{\beta/\lambda}-1\right)^2}-\frac{\lambda q}{e^{\gamma/\lambda}-1}+\frac{1}{4}\gamma q\text{csch}^2\left(\frac{\gamma}{2\lambda}\right)\right)+(p+q-1)\left(-e^{-\frac{x}{\lambda}}\right)(x-\lambda)}{\lambda^2\left(\sinh\left(\frac{x}{\lambda}\right)\left(\frac{p}{e^{\beta/\lambda}-1}+\frac{q}{e^{\gamma/\lambda}-1}\right)-(p+q-1)e^{-\frac{x}{\lambda}}\right)} & x_i<\beta,\gamma \\[3ex]
\dfrac{\frac{px\left(e^{\beta/\lambda}+1\right)e^{-\frac{x}{\lambda}}}{2\lambda^3}-\frac{\beta p e^{\frac{\beta}{\lambda}-\frac{x}{\lambda}}}{2\lambda^3}-\frac{p\left(e^{\beta/\lambda}+1\right)e^{-\frac{x}{\lambda}}}{2\lambda^2}+\frac{x(-p-q+1)e^{-\frac{x}{\lambda}}}{\lambda^3}-\frac{(-p-q+1)e^{-\frac{x}{\lambda}}}{\lambda^2}+\frac{\gamma q e^{\gamma/\lambda}\sinh\left(\frac{x}{\lambda}\right)}{\lambda^3\left(e^{\gamma/\lambda}-1\right)^2}-\frac{qx\cosh\left(\frac{x}{\lambda}\right)}{\lambda^3\left(e^{\gamma/\lambda}-1\right)}-\frac{q\sinh\left(\frac{x}{\lambda}\right)}{\lambda^2\left(e^{\gamma/\lambda}-1\right)}}{\frac{p\left(e^{\beta/\lambda}+1\right)e^{-\frac{x}{\lambda}}}{2\lambda}+\frac{(-p-q+1)e^{-\frac{x}{\lambda}}}{\lambda}+\frac{q\sinh\left(\frac{x}{\lambda}\right)}{\lambda\left(e^{\gamma/\lambda}-1\right)}} & \beta\leq x_i<\gamma \\[3ex]
-\dfrac{\gamma+\lambda+\frac{p(\beta-\gamma)e^{\beta/\lambda}+\gamma(p+q-2)}{p\left(e^{\beta/\lambda}-1\right)+q\left(e^{\gamma/\lambda}-1\right)+2}-x}{\lambda^2} & \beta,\gamma\leq x_i
\end{cases}$$

We then add a value proportional to the derivative by a factor $s$ to the current value of $\lambda$ and check the likelihood. If it increases, we keep the new value and repeat until convergence. If at any point the proposed step decreases the likelihood, we decrease $s$ by half and retry until we find the appropriate step size.

#### 4.2 Output Interpretation

Given the results of the independent triplet analyses, we can synthesize them by looking at the output for the three tree topologies in each triplet as a group. For two distribution models, $C_2$ corresponds to an estimate of the time in units of $2N_e$ generations that the two sister lineages were isolated from the third species. $\pi_2$ is an estimate of what fraction of the gene trees in the input were not generated via ILS. $\lambda$ corresponds to the single best estimate of $\theta/2$ for the branch in the species tree where the two sister lineages coalesced, averaging over both the ILS and non-ILS gene trees.
In order to make statements about the full gene trees, we set a significance threshold for the probability of introgression (90%), and considered how many triplets at a locus reported introgression probabilities above that.

#### 4.3 Tree Simulation Tests

To verify the behavior of the procedure, we simulated coalescent trees under a variety of parameters and observed the error in the parameter estimates (Figure 5). As we see only error in the form of false negatives (panel 5.b), and only at low mixing proportions (panel 5.c), these plots combined demonstrate that the error arises almost entirely as a result of the non-exponential introgression distribution being mixed in at low frequencies (< 65 trees), which is not observed in the *Heliconius* dataset. In addition, we find in larger tests that this threshold is a result of the number of points rather than the proportion itself.

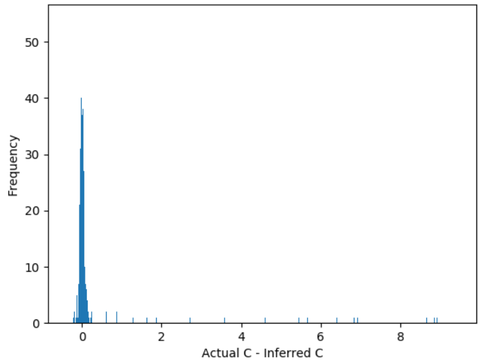

(a) Plot of the error in the estimate of the $C$.

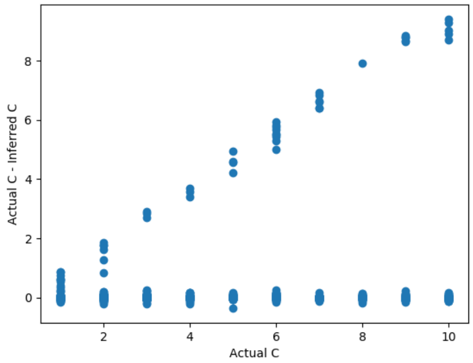

(b) Errors demonstrating the lack of false positives in $C$ estimates.

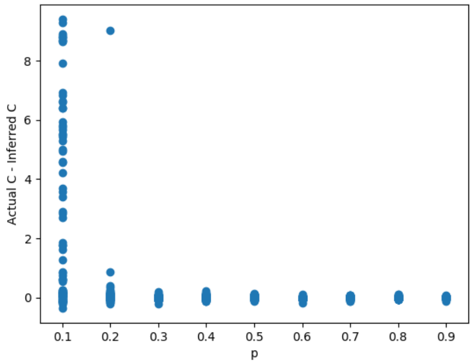

(c) Error in $C$ as a function of mixing proportion, showing the error is due to a detection threshold.

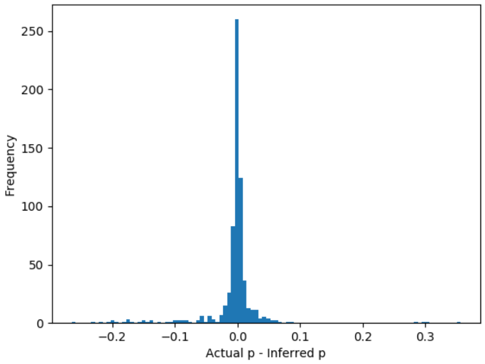

(d) Plot of the error in the estimate of the mixing proportion of the non-exponential distribution, $p$.

Figure 5: Set of plots describing the behavior of the error in parameter estimates for a set of 650 coalescent trees each for 100 parameter combinations with 10 replicates each.

**Section 10: Chromosome 2 Inversion**

*return to top*

As mentioned in the main text, a large region on Chromosome 2 displayed a topology discordant with the most common species relationships in the *H. erato* group. This region corresponds to a previously described inversion between *H. erato* and *H. melpomene.* We confirmed that this was in fact that inverted region by mapping all *erato* clade contigs onto Hmel2. We found clear evidence of an inversion in the *H. erato demophooon* reference genome, as well as the *H. hecalesia de novo* assembly. We also found contigs in *H. sara* and *H. demeter* that mapped across one of the inversion breakpoints, indicating that they are in the same orientation as *H. melpomene.* However, the breakpoints of this inversion are flanked by repetitive sequence that was difficult to align among all species, especially the left-side breakpoint. Therefore, we infer the orientation of the inversion in *H. himera* and *H. telesiphe* based on the local topology (Figure S10.1A; Topology is Tree3 in main text Figure 2B). As was done for the Chromosome 15 inversion in the main text, we used the branch length method in combination with $D_{XY}$ to evaluate whether the discordant history seen in this region was due to ILS, inversion, or some other process. Again, we used the *H. sara, H. telesiphe, H. erato* triplet, and again we found that the internal branch length in the inversion was much larger than expected under simple ILS (Figure 10.1E). However, in contrast to the Chromosome 15 inversion, normalized $D_{XY}$ between *H. sara* and *H. telesiphe* was very close to the genome-wide average (Figure 10.1F). This combination of statistics supports a history in which the inversion originated in the ancestor of the *erato-sara* clade and remained polymorphic for some time before speciation of the taxa we sampled.

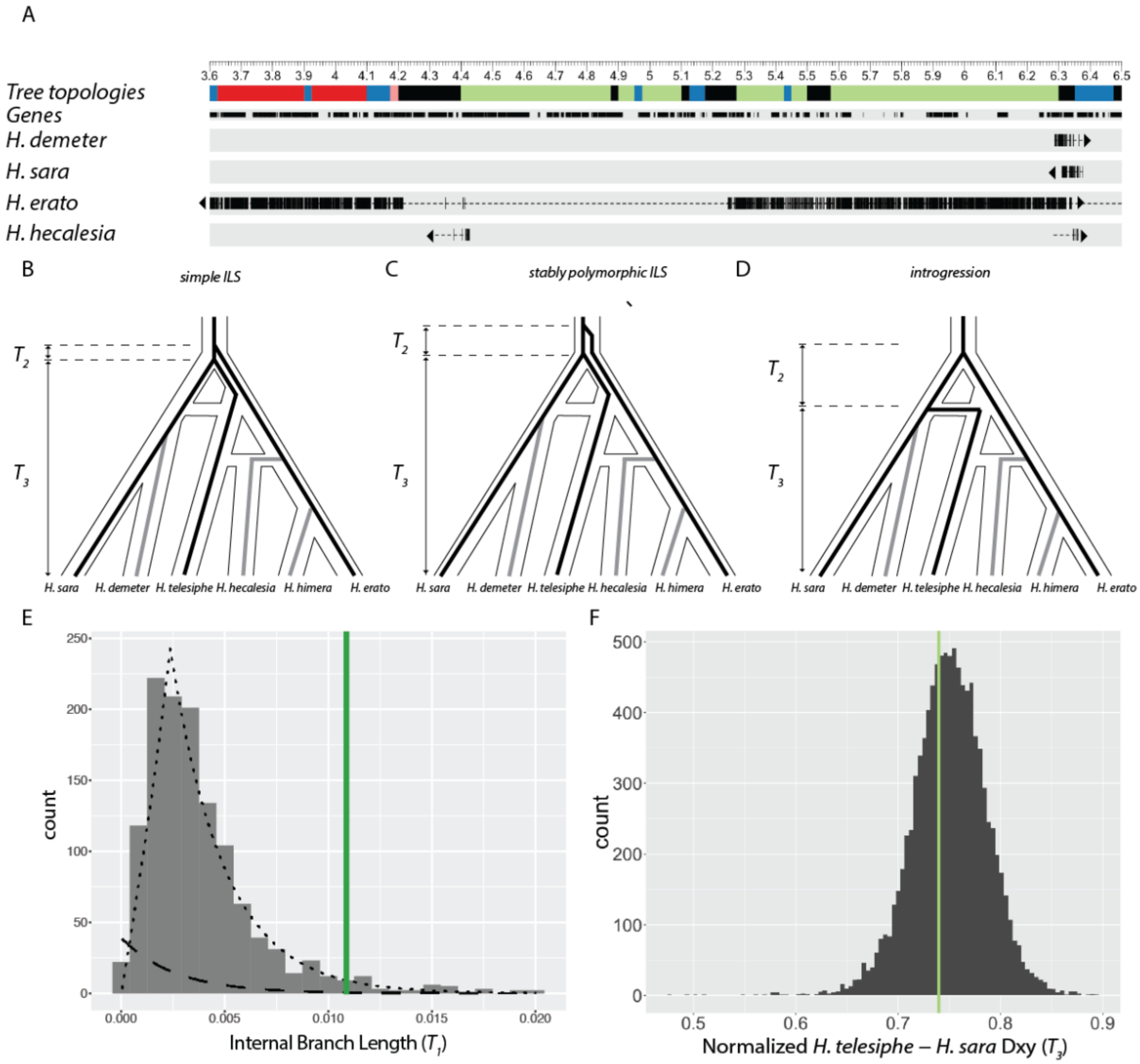

**Figure S10.1: An inversion on chromosome 2**

***A.*** *Map of 3 Mb region on Chromosome 2. Coordinates are in terms of the Hmel 2.5 reference order, and ticks are in Mb. Colors correspond to those in figure 2B. Genes are shown as black rectangles. Each line below shows the mapping of a single scaffold. Aligned sections of each scaffold are shown as thick bars, while unaligned sections are shown as dotted lines to indicate the relative position of the alignment within the scaffold. Only regions that were not identified as multiple-copy or repetitive are shown. Arrows indicate direction of alignment.* ***B-D. Hypothetical evolutionary histories.*** *In all cases, the histories of the three species used in the "triplet gene tree" method – H. erato, H. telesiphe, and H. sara – are shown as black lines, while lineages not included are shown as grey lines.* ***B*** *shows the scenario expected in the case of simple ILS.* ***C*** *shows another scenario of ILS, but in this case the inversion is polymorphic for some time in the common ancestor, leading to a longer internal branch for the inversion.* ***D*** *shows the case of introgression from the H. sara+H. demeter ancestor into the H. telesiphe+H. hecalesia ancestor.* ***E. Distribution of internal branch lengths****. Histogram is of internal branch* *lengths ($T_2$) in the H. erato,(H. telesiphe, H. sara) topology. The inferred ILS distribution is shown as a dashed line, and the inferred introgression distribution is shown as a dotted line. The average internal branch length in the inversion is shown as a green vertical line.* ***F. Normalized H. telesiphe-H. sara $D_{XY}$.*** *Normalized $D_{XY}$ ($T_3$) is calculated as H. telesiphe-H. sara $D_{XY}$ divided by the mean pairwise $D_{XY}$ among all species in each region. Mean normalized $D_{XY}$ in the inversion is shown as a green vertical line.*

**References**
[return to top](#)

1. Smit, A., Hubley, R. & Green, P. *2013–2015. RepeatMasker Open-4.0*. (2013).
2. Vurture, G. W. *et al.* GenomeScope: fast reference-free genome profiling from short reads. *Bioinformatics* **33,** 2202–2204 (2017).
3. Davey, J. W. *et al.* Major improvements to the *Heliconius melpomene* genome assembly used to confirm 10 chromosome fusion events in 6 million years of butterfly evolution. *G3* **6,** 695–708 (2016).
4. Van Belleghem, S. M. *et al.* Complex modular architecture around a simple toolkit of wing pattern genes. *Nat. Ecol. Evol.* **1,** 52 (2017).
5. Simão, F. A., Waterhouse, R. M., Ioannidis, P., Kriventseva, E. V. & Zdobnov, E. M. BUSCO: assessing genome assembly and annotation completeness with single-copy orthologs. *Bioinformatics* **31,** 3210–3212 (2015).
6. Hickey, G., Paten, B., Earl, D., Zerbino, D. & Haussler, D. HAL: a hierarchical format for storing and analyzing multiple genome alignments. *Bioinformatics* **29,** 1341–1342 (2013).
7. Hubisz, M. J., Pollard, K. S. & Siepel, A. PHAST and RPHAST: phylogenetic analysis with space/time models. *Brief. Bioinformatics* **12,** 41–51 (2011).
8. *https://github.com/glennhickey/progressiveCactus*. Available at: httpsgithub.comglennhickeyprogressiveCactus. (Accessed: 1st May 2016)
9. Paten, B. *et al.* Cactus: Algorithms for genome multiple sequence alignment. *Genome Research* **21,** 1512–1528 (2011).
10. Ahola, V. *et al.* The Glanville fritillary genome retains an ancient karyotype and reveals selective chromosomal fusions in Lepidoptera. *Nat. Commun.* **5,** 4737 (2014).
11. Nowell, R. W. *et al.* A high-coverage draft genome of the mycalesine butterfly *Bicyclus anynana*. *Gigascience* **6,** 1–7 (2017).
12. Zhan, S. *et al.* The genetics of monarch butterfly migration and warning colouration. *Nature* **514,** 317–321 (2014).
13. Nishikawa, H. *et al.* A genetic mechanism for female-limited Batesian mimicry in *Papilio* butterfly. *Nat. Genet.* **47,** 405–409 (2015).
14. Cong, Q., Borek, D., Otwinowski, Z. & Grishin, N. V. Skipper genome sheds light on unique phenotypic traits and phylogeny. *BMC Genomics* **16,** 639 (2015).
15. The International Silkworm Genome Consortium. The genome of a lepidopteran model insect, the silkworm *Bombyx mori*. *Insect Biochem. Mol. Biol.* **38,** 1036–1045 (2008).
16. You, M. *et al.* A heterozygous moth genome provides insights into herbivory and detoxification. *Nat. Genet.* **45,** 220–225 (2013).
17. Davey, J. W. *et al.* No evidence for maintenance of a sympatric *Heliconius* species barrier by chromosomal inversions. *Evolution Letters* **1,** 138–154 (2017).
18. Hoang, D. T., Chernomor, O., Haeseler, von, A., Minh, B. Q. & Vinh, L. S. UFBoot2: Improving the ultrafast bootstrap approximation. *Mol. Biol. Evol.* **35,** 518–522 (2018).
19. Kalyaanamoorthy, S., Minh, B. Q., Wong, T. K. F., Haeseler, von, A. & Jermiin, L. S. ModelFinder: fast model selection for accurate phylogenetic estimates. *Nat. Methods* **14,** 587–589 (2017).

20. Nguyen, L.-T., Schmidt, H. A., Haeseler, von, A. & Minh, B. Q. IQ-TREE: a fast and effective stochastic algorithm for estimating maximum-likelihood phylogenies. *Mol. Biol. Evol.* **32,** 268–274 (2015).
21. Mirarab, S. *et al.* ASTRAL: genome-scale coalescent-based species tree estimation. *Bioinformatics* **30,** i541–8 (2014).
22. Jiggins, C. D. *et al.* Sex-linked hybrid sterility in a butterfly. *Evolution* **55,** 1631–1638 (2001).
23. Naisbit, R. E., Jiggins, C. D., Linares, M., Salazar, C. & Mallet, J. Hybrid sterility, Haldane's rule and speciation in *Heliconius cydno* and *H. melpomene*. *Genetics* **161,** 1517–1526 (2002).
24. Pinharanda, A., Martin, S. H., Barker, S. L., Davey, J. W. & Jiggins, C. D. The comparative landscape of duplications in Heliconius melpomene and Heliconius cydno. *Heredity* **118,** 78–87 (2017).
25. Martin, S. H., Davey, J., Salazar, C. & Jiggins, C. Recombination rate variation shapes barriers to introgression across butterfly genomes. *bioRxiv* 297531 (2018). doi:10.1101/297531
26. Wickham, H. *ggplot2: elegant graphics for data analysis*. (Springer, 2016).
27. Martin, S. H., Davey, J. W. & Jiggins, C. D. Evaluating the use of ABBA-BABA statistics to locate introgressed loci. *Mol. Biol. Evol.* **32,** 244–257 (2014).
28. Patterson, N. *et al.* Ancient admixture in human history. *Genetics* **192,** 1065–1093 (2012).
29. Peter, B. M. Admixture, population structure, and F-statistics. *Genetics* **202,** 1485–1501 (2016).
30. Zairis, S., Khiabanian, H., Blumberg, A. J. & Rabadan, R. Genomic data analysis in tree spaces. *arxiv.org* (2016).
